# Characterization of LC-MS based urine metabolomics in healthy subjects across the life span

**DOI:** 10.1101/2020.11.12.379131

**Authors:** Xiaoyan Liu, Xiaoyi Tian, Qinghong Shi, Haidan Sun, Jing Li, Xiaoyue Tang, Zhengguang Guo, Ying Liu, Jie Ma, Na Ren, Fang Jin, Chengyan He, Wenqi Song, Wei Sun

## Abstract

Previous studies reported that gender and age could influence urine metabolomics, which should be considered in biomarker discovery. As a consequence, for the baseline of urine metabolomics characteristics, it becomes critical to avoid confounding effects in clinical cohort studies. In this study, we provided a comprehensive lifespan characterization of urine metabolomics in a cohort of 348 healthy children and 315 adults aged 1 to 70 years using liquid chromatography coupled with high resolution mass spectrometry. Our results suggested that gender-dependent urine metabolites are much greater in adults than in children. The pantothenate and CoA biosynthesis and alanine metabolism pathways were enriched in early life. Androgen and estrogen metabolism showed high activity during adolescence and youth stages. Pyrimidine metabolism was enriched in the old stage. Based on the above analysis, metabolomic characteristics of each age stage were provided. This work could help us understand the baseline of urine metabolism characteristics and contribute to further studies of clinical disease biomarker discovery.

## Introduction

In recent years, urine metabolomics has been widely used in disease biomarker discovery (1, 2). Previous studies have reported that gender and age could influence urine metabolomics, which should be considered during biomarker discovery (3). Therefore, understanding the baseline of urine metabolomics characteristics would be helpful for better understanding metabolism status under healthy conditions and discovering disease-specific metabolism disorders.

Investigation of urine metabolomics variation in a healthy population has been performed in children and adults using various approaches, including nuclear magnetic resonance (NMR) and mass spectrometry coupled to either gas (GC-MS) or liquid chromatography (LC-MS). In 2016, urinary metabolites from 30 healthy children were assessed at 6 months and 1, 2, 3, and 4 years of age by using NMR spectroscopy. Amino acid metabolism was significantly different between infants aged 6 months and 1 year, whereas carbohydrate metabolism was significantly different between children aged 2 and 3 years (4). In 2018, a six European centers research on metabolome differences in children aged 6 to 11 were performed using NMR and targeted LC-MS/MS. Both urine and serum metabolomes were found to be associated with age, sex, BMI and dietary habits (5). For the metabolic phenotypes in adults, in 2015, Etienne A. et al. characterized the urinary metabolomes of 183 healthy subjects using an LC-MS platform. A total of 108 metabolites related to amino acid metabolism, the carnitine shuttle and the TCA cycle were found to be affected by age, gender or BMI (2). Recently, in 2018, Sili Fan et al. applied metabolomics profiling on urine samples from 60 healthy males and females using GC-TOF-MS. Saturated fatty acids, TCA cycle intermediates, and butyrate were found to be significantly related to the effect of gender (6). In 2018, we conducted a urine metabolomics study in 203 healthy adults to find age- and gender-dependent urine metabolites. Metabolic pathways, such as tryptophan metabolism, citrate cycle, and pantothenate and CoA biosynthesis, were found to be related to gender and age (7). Despite the large efforts in characterizing urine metabolomics in healthy subjects, most of the previous studies were focused on the confounding factors of the urine metabolome. While the metabolic characterization of a population at each stage across the life span was unavailable until the present study.

In the present study, a large sample size including 348 children (aged from 1 to 18 years) and 315 adults (aged from 20 to 70 years) with balanced gender from China was enrolled. We aimed to characterize urine metabolic features in different age stages for males (boys) and females (girls). HRLC-MS-based metabolic profiling was utilized to characterize the urine metabolome of each individual. In addition to gender- and age-related metabolites and pathways, the key metabolites that played important roles during different gender or age stages were identified and analyzed. Identification of these significant metabolites and metabolite networks would be helpful for understanding the physiological functions during different life stages and would be useful for future disease biomarker discovery.

## Results

### Data quality control

The workflow for the present study is shown in Fig. 1. To reduce experimental variations from the sampling process, standard sampling procedures, including sampling time and sampling processing, were performed by a specialist. Data from the two centers’ sample were analyzed separately. Overall, 730 injections were carried out. Principal component analysis was performed to evaluate the variation of samples (Fig S1a). A good cluster of QC samples indicated good stability of the analytical platform.

**Fig 1.**
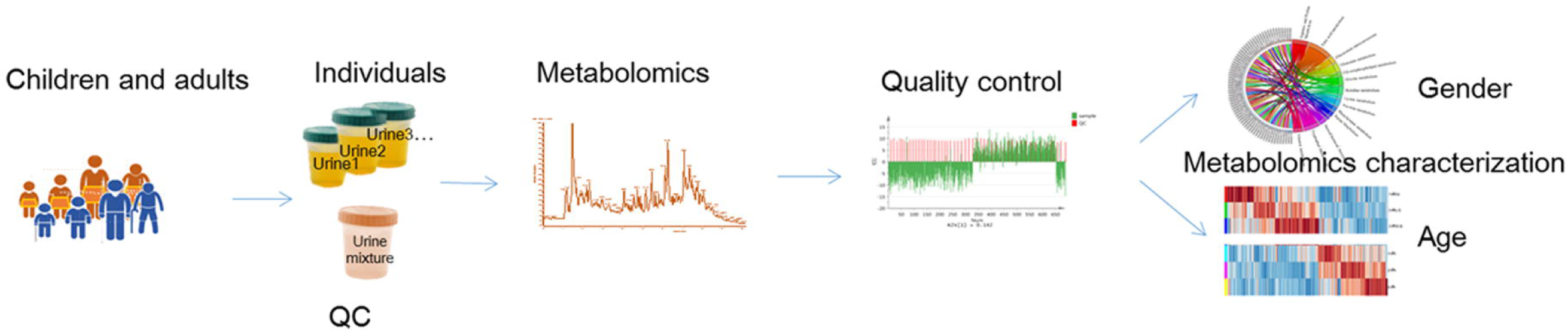
The workflow of this study.

### Gender-dependent metabolomics in children and adults

A total of 291 metabolites were identified and subjected to further analysis (Table S1). Children and adult urine samples were collected from two clinical centers, and gender-dependent metabolites were assessed in the children and adult samples, respectively.

Urine metabolic differences between males and females were assessed using PCA and OPLS-DA models. Scatter plots showed that gender differences in the urine metabolites of adults are more apparent than in children (OPLS-DA: children: R2Y = 0.599, Q2 = 0.379; adults: R2Y = 0.928, Q2 = 0.768. Fig. S1b and S1c). Metabolites with VIP values above the threshold value 1 and p values below the significance threshold were considered gender dependent. A total of 42 metabolites were found to be significantly different between boys and girls, and 98 metabolites were different between males and females (Fig. 2a and 2b, Tables S2 and S3). A total of 17.9% (21) of these metabolites showed gender differences in both children and adult populations with the same change trend (Fig. 2c). Guanidoacetic acid, 5-hydroxyindoleacetic acid, dopamine, 5’-methylthioadenosine and indoleacrylic acid showed higher levels in females. The metabolites deoxyinosine, cotinine glucuronide, dopamine glucuronide and L-formylkynurenine showed higher levels in males. Gender differences of these common differential metabolites in the adult population were larger than those in the children population (Fig. 2c). Specifically, the metabolites cortisol, uric acid, 18-hydroxycortisol, deoxycholic acid and glycine conjugate were found to be gender-dependent only in the children population. Metabolites of deoxyuridine, pantothenic acid, riboflavin, and 3-hydroxytetradecanedioic acid were gender-dependent only in the adult population. These metabolites suggested differential metabolic status between children and adults.

**Fig 2.**
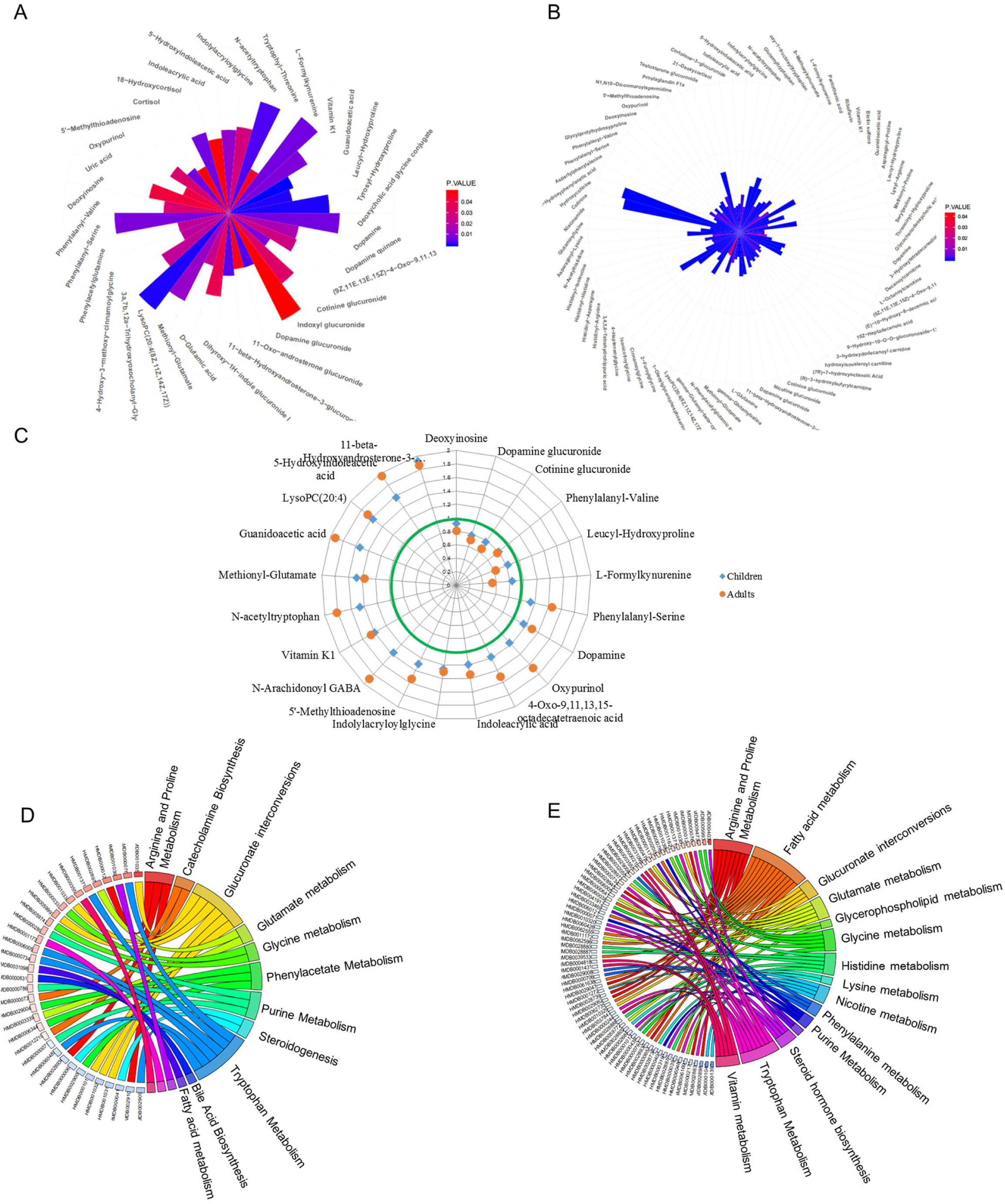
Representative gender-differential metabolites and metabolism pathways in children and adults. **a.** polar plot for fold change (Female/Male) and p value of gender-dependent metabolites in children. The length of the column indicates the fold change, and the color indicates the p value. **b.** Polar plot for fold change (Female/Male) and p value of gender differential metabolites in adults. **c.** Fold change value of 50 common gender-dependent metabolites in children and adults. Values inside the green circle represent metabolites higher in males (boys), and values outside the green circle represent metabolites higher in females (girls). **d.** Gender-dependent metabolism pathways in children. Metabolites involved in some pathway were connected using lines. **e.** Gender-dependent metabolism pathways in adults. Metabolites involved in some pathway were connected using lines.

Pathway enrichment analysis would provide an overview of gender-dependent metabolism status in children and adults. Arginine, proline, and tryptophan metabolism and glucuronate interconversions were found to be gender-dependent in both the children and adult populations (Fig. 2d and 2e). Additionally, several specific pathways were found to be gender-dependent in adults only, including nicotine metabolism, nicotinate and nicotinamide metabolism, steroid hormone biosynthesis, lysine metabolism and histidine metabolism. These results indicated the commonness and differences of gender-dependent metabolism characteristics in children and adults, probably resulting from different physiological characteristics, dietary habits or other environmental factors (11). The detailed gender-associated pathways in each age stage are listed in Table S4.

### Age-dependent metabolomics in children and adults

Age is another important factor in metabolism status. PCA was firstly performed, showing apparent separation of different groups (Fig. S2a). To determine metabolites related to age, PLS-DA modeling was performed on urine metabolite profiles of populations with different age stages. Since the metabolism status of the population was different between sexes, the analyses for age in children and adults were performed separately for males and females.

In the children population, the PLS-DA three-component score plot showed clear associations of metabolite profiles with age in boys (R2Y = 0.719, Q2 = 0.528, Fig. 3a, Fig. S2b), indicating significant metabolism differences among different age stages. The same trend was observed in girls (R2Y = 0.78, Q2 = 0.564, Fig. 3b, Fig. S2b). In both boys and girls, two clusters of metabolites were found, one cluster with metabolites decreasing with age (Component 1); the second cluster with metabolites showing the highest level in the primary school aging 7 to 12 (Component 2) (Fig. 3c). The first cluster contributes most to age variation in both girls and boys. In the adult population, clear separations were also observed for the three age groups in males and females (PLS-DA-male: R2Y = 0.652, Q2 = 0.17; PLS-DA-female: R2Y = 0.583, Q2 = 0.362, Fig. S2b, Fig. S2c and Fig. S2d). Similar to children, metabolites decreasing with age contributed most to age differences (Fig S2e). According to the significance threshold, a total of 98 and 76 metabolites were selected as age-dependent in boys and girls, respectively. For the adult population, 55 and 80 metabolites were age-dependent in males and females, respectively (Table S5, S6, S7 and S8). These metabolites were further submitted for further pathway analysis.

**Fig 3.**
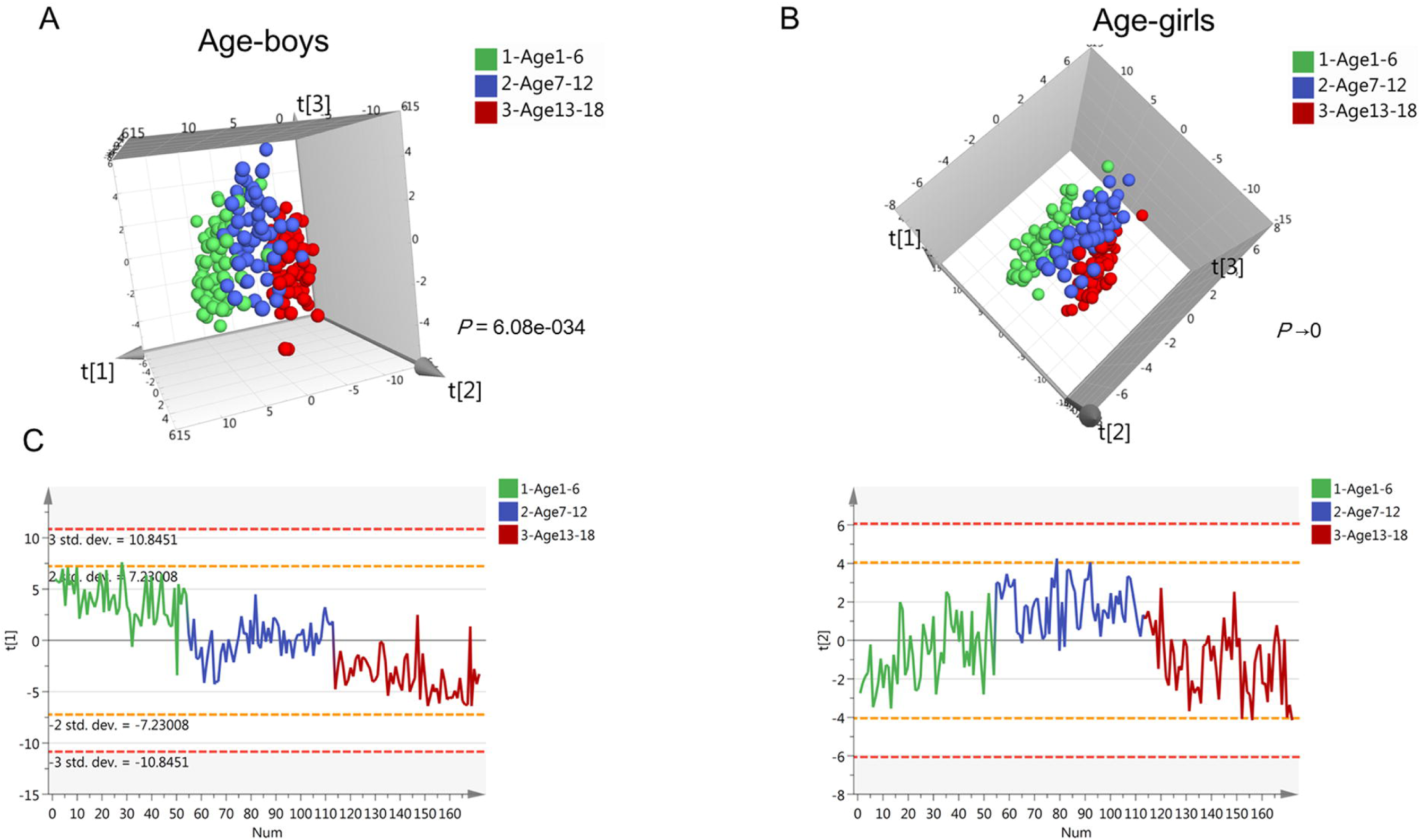
Urine metabolomics variation with age in children. a. PLS-DA score plot of urine metabolomics of different ages in boys. b. PLS-DA score plot of urine metabolomics of different ages in girls. c. Change trends of the first and second principal components of PLS-DA in children; these two principal components contribute most to the age separation in children.

Since the urine samples of children and adults were from two centers, we examined differential metabolism pathways in the two populations, respectively. In the children population, the metabolites involved in histidine metabolism, riboflavin metabolism, pantothenate and CoA biosynthesis and fatty acid biosynthesis were found to change with age in both boys and girls. Tryptophan metabolism was age-dependent in boys, indicating sexual dimorphism of age-associated metabolites. In the adult population, most differential pathways were the same as the children population. However, in adult males, steroid hormone biosynthesis was found to be age dependent, although this difference was found in girls rather than boys, partly resulting from later sexual development in boys. The detailed age-dependent metabolism pathways for males (boys) and females (girls) are listed in Table S4.

### Metabolic characteristics in males and females across life span

According to the above results, the urine metabolites of children and adults showed both gender and age dependent effects. Furthermore, we detailed the metabolic differences affected by both gender and age. The p values of OPLS-DA for gender separation during each stage were used to evaluate the significance of gender difference. The results showed a parabolic trend of gender differences during life span, with less significance in the pre- and primary school stages, high significance during the secondary school, youth and middle stages, and less significance during the old stage (Fig 4).

**Fig 4.**
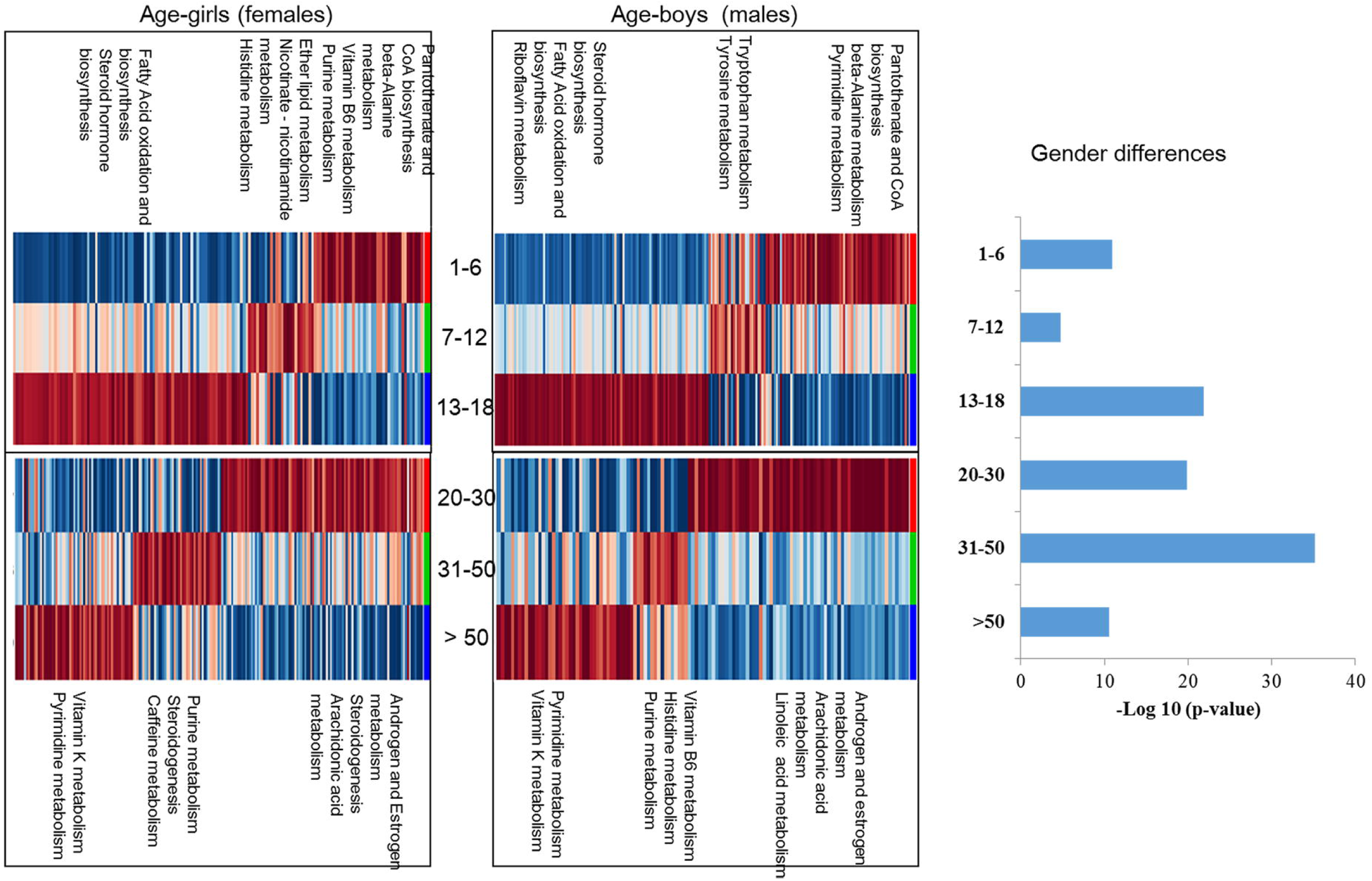
Metabolic characteristics for different gender and age stages. Age-dependent metabolic pathways were enriched based on metabolites with the highest level in each age group. The KEGG database was the background pathway database. The significance of gender differences was evaluated using the p value of OPLS-DA models. The cutoff for the p value was 0.05.

Second, we examined the main metabolic characteristics for each age stage in males (boys) and females (girls). Metabolites showing the highest level for each group were submitted to pathway analysis, indicating the specific metabolism characteristics during a certain life stage. For each age stage, representative pathways were listed for males (boys) and females (girls) (Fig 4 and Table 2). In the population aged 1-6, the main metabolic feature was energy metabolism for growth demand, showing high activity of pantothenate and CoA biosynthesis and beta-alanine metabolism. In the population aged 7-12, higher lipid, glucose and amino acid metabolisms were the main features. The activation of three major energy supply pathways indicated higher energy demand during this stage. In the population aged 13-18, the metabolic features in boys and girls showed some differences, most likely due to sexual development. Spermidine and spermine biosynthesis and riboflavin metabolism showed high activity in boys. In addition, fatty acid oxidation and biosynthesis were active in girls. In populations aged 20-30, organ metabolism was the most active. Androgen and estrogen metabolism, steroidogenesis and arachidonic acid/linoleic acid metabolism showed higher activity. During the 30- to 50-year-old stage, males and females showed different metabolic features. In males, vitamin B6 and purine metabolism were active. In females, steroidogenesis and caffeine metabolism were the main metabolic features. For the last age group, males and females showed similar metabolic features, with high levels of the metabolites vitamin K, pyrimidine and caffeine.

### Key metabolite variations with gender and age

In the present work, several important pathways were found to be significantly varied in males (boys) and females (girls) or with aging. For example, tryptophan metabolism showed gender differences, with metabolites of 5-hydroxyindoleacetic acid showing significant variation between males and females. Pantothenate and CoA biosynthesis, steroid hormone biosynthesis and pyrimidine metabolism showed age dependence, with metabolites of pantothenate, testosterone glucuronide and 8-hydroxy-7-methylguanine showing different levels with age. Herein, we evaluated the continuous change trends of the represented significant metabolites involved in gender and age-dependent pathways, giving an overview of relative levels across the life span in males (boys) and females (girls). The annotations of these metabolites were provided in Fig.S3

Metabolites of 5-hydroxyindoleacetic acid showed higher levels in females (girls) than males (boys) across the life span (Fig 5a), indicating high activity of tryptophan metabolism in females (girls). Metabolites of phenylalanyl-valine showed higher levels in males (boys) than in females (girls). These differences in adults were larger than those in children. For viewing the change trend with aging, age series lines of the representative metabolites were plotted.

**Fig 5.**
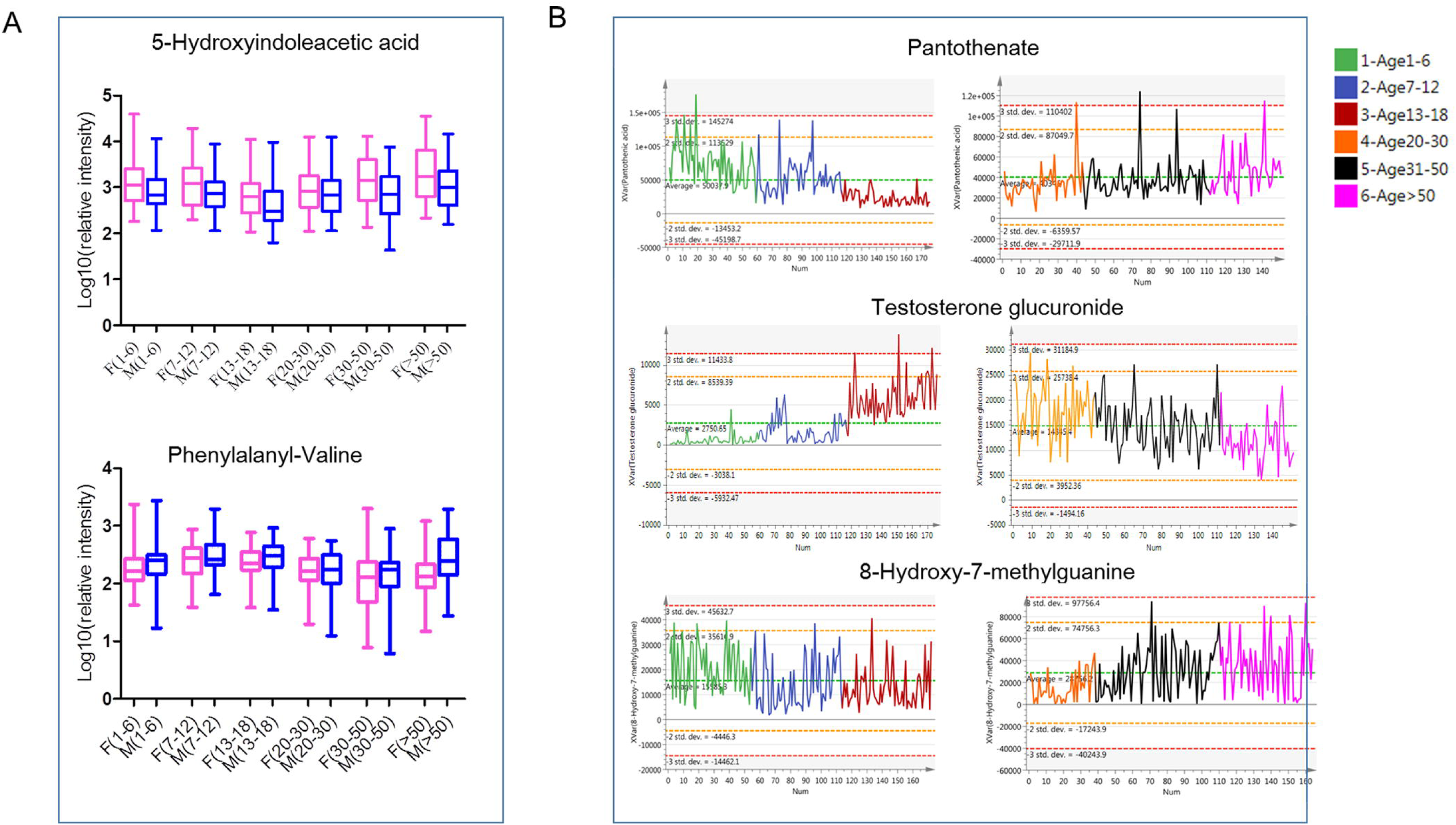
Key metabolite variation with gender and age. a. Relative intensity of gender-differential metabolites, 5-hydroxyindoleacetic acid and phenylalanyl-valine in male (boy) and female (girl) during different age stage. b. Relative intensity change of age differential metabolites, pantothenate, testosterone glucuronide and 8-hydroxy-7-methylguanine with age in males.

As shown in Fig 5b and Fig S4, pantothenate, also called vitamin B5, is a vitamin required to sustain life. Pantothenate is needed to form CoA and is thus critical in the metabolism and synthesis of carbohydrates, proteins, and fats. Pantothenate was the highest in the early life stage, contributing to nutritional demands for rapid development.

Steroids are characteristic metabolites and showed the highest level in the youth life stage. Testosterone glucuronide is the downstream metabolites of testosterone. Herein, we detected some testosterone glucuronide secreted in the urine. The level of testosterone glucuronide was very low in children and increased with puberty and youth. In the adult population, a high level of testosterone glucuronide was also observed in the youth and then decreased with aging in the old. Steroid level changes were tightly associated with sexual development during the life span.

8-Hydroxy-7-methylguanine is an endogenous methylated nucleoside. Methylated nucleoside in urine could be derived from covalently bound adducts in DNA, which are formed by exposure to various carcinogens. The level of 8-hydroxy-7-methylguanine remained fairly constant from early life to youth. However, it increased significantly during the midlife stage and reached the highest level during the old stage.

## Discussion

In the present study, we tested urine samples from a large population of children and adults to characterize metabolic features in different age stages. Urine metabolism characteristics from early life to old age were profiled, which is the first systematic comparison study based on such a large and healthy population.

### Gender-dependent metabolism status in children and adults

In the present study, we characterized the urine metabolomes of males and females in children and adults. Consistent with previous studies (4,6,7), amino acid metabolism, including tryptophan metabolism and arginine and proline metabolism, showed gender differences in both children and adults. In females/girls, tryptophan metabolites, indoleacrylic acid, indolylacryloylglycine and 5-hydroxyindoleacetic acid showed higher levels compared with males/boys. 5-Hydroxyindoleacetic acid is a breakdown product of serotonin that is excreted in the urine and is mainly involved in serotonin degradation and serotonin receptor signaling regulation. The higher level in females could be partly explained by greater precursor, such as tryptophan, availability (12). Studies on experimental animals have revealed the effects of gonadal hormones on the indoleamine system, and in the rat brain, 5-hydroxyindoleacetic acid is higher in females than in males (13). These results were consistent with our present urine results.

In addition, children and adults showed different gender-dependent metabolic statuses. In the child population, purine metabolites, steroid metabolites and amino acid metabolites showed specific gender dependence. For example, uric acid, an oxidation product of purine, showed a significantly higher level in boys compared with girls but showed no gender variations in adults. Renal excretion of uric acid in children differs from that in adults. It was reported that the younger the child, the greater the excretion of uric acid (14). Altered urine uric acid level is an indispensable marker in detecting rare inborn errors of metabolism (15). The level of uric acid in urine was higher in boys than girls, probably contributing to the higher prevalence of hyperuricemia in boys than in girls (16).

Compared to children, we identified more specific gender-dependent metabolites in adults. Most of the specific gender-dependent metabolites in adults showed higher levels in males, including fatty acids, acylcarnitines, steroid hormones and dipeptides. Higher levels of acylcarnitines in males were found in our previous study, indicating a higher activity of fatty acid oxidation and energy production in males compared to females (7). In addition, a higher level of steroid hormone biosynthesis was found in males, consistent with previous studies (17).

### Age-dependent metabolism status in children and adults

For children and adults, we found many common age-dependent pathways between boys (males) and girls (females), indicating common metabolism status variations with age in humans. For instance, pantothenate and CoA biosynthesis, fatty acid biosynthesis and tryptophan metabolism were found to be changed with age in both children and adults. These pathways correspond to energy demand changes with aging (4). Interestingly, tryptophan metabolism was only found to be age dependent in boys and males, while no significant changes were found in girls and females. Metabolites of tryptophanol and 5-methoxytryptophan were reported to be associated with increased cellular anti-inflammatory and blood circulation properties (18), probably reflecting age-associated metabolism activity differences between genders. In addition, we found that the pathway of steroid hormone biosynthesis was changed with age in adults and children. As a whole, steroid hormone biosynthesis showed high activity during the adolescent and youth stages, corresponding to sexual development (19).

Additionally, children and adults showed different age-dependent metabolic statuses. Particularly in the children population, the fatty acid biosynthesis pathway was found to be age dependent. The results showed a positive correlation with increasing age. Fatty acids constitute a large energy source for the body. Increased fatty acid metabolism indicated high ATP generation with age in a children population (20). In adults, pyrimidine metabolism and caffeine metabolism were found to be age dependent. Pyrimidine metabolism was found to be positively correlated with aging, showing the highest level in the old adults. Deoxyuridine, a naturally occurring nucleoside, is considered to be an antimetabolite that is converted to deoxyuridine triphosphate during DNA synthesis. Disturbance of DNA synthesis may modulate the aging process and contribute to the high incidence of cancer with aging (21).

### Metabolomic characteristics during each age stage

#### Metabolomic characteristics during the pre- and primary school stages

During early life in the preschool and primary school stages, gender differences were relatively smaller than in other age stages, as shown in Fig 4. During the preschool stage, pathways of pantothenate and CoA biosynthesis, pyrimidine metabolism, vitamin B6 and alanine metabolism showed high activity in girls and boys. These active pathways were associated with energy and nutrient supply. These metabolic characteristics correspond to the physiological characteristics during this life stage, high metabolism activity for rapid growth and development demands (4). Research on children aged from 2-7 years old suggested that pantothenate and CoA biosynthesis, pyrimidine metabolism and vitamin B6 metabolism were significantly related to autism spectrum disorder (ASD) (22). These results highlight the importance for these pathways to maintain homeostasis during preschool.

During primary school stage, the most active metabolism pathway is tryptophan metabolism, lipid metabolism, nicotinate and nicotinamide metabolism and histidine metabolism. These metabolic features were corresponding to the main physiological characteristic-visual development, blood circulation increasing and rapid metabolism activity during this stage. Although gender differences were small during the primary school stage, metabolites with the highest level in boys and girls showed specific features. In boys, tryptophan metabolites, such as 5-methoxytryptophan, were found to show the highest level in boys. 5-Methoxytryptophan is an endogenous tryptophan metabolite with anti-inflammatory properties (18). In addition, 5-methoxytryptophan was reported to be involved in the cyclic metabolism of the retina (23), ventricular remodeling and maintaining liver function (24, 25). In girls, urocanic acid, a breakdown (deamination) product of histidine, showed the highest level. Urocanic acid is one of the essential components of human skin (26). It could accumulate in the epidermis and may be both a UV protectant and an immunoregulator. A higher level of these metabolites in girls may contribute to skin development during this age stage. It was reported that skin disorders are more common among girls than boys aged 6 to 17 years (27), which could be affected by the immunomodulatory effects of urocanic acid (28).

#### Metabolomic characteristics during adolescence and youth

During adolescence and the youth stage, gender differences become more significant, partly due to changes in hormone and endocrine levels. The main metabolic feature during these stages was fatty acid oxidation and biosynthesis, androgen and estrogen metabolism and steroidogenesis. These metabolic characteristics correspond to pubertal development, neurodevelopmental changes and heightened stress sensitivity during adolescence and youth stages. Cortisol, androstenol, testosterone, and their glucuronide metabolites showed higher levels during this period. Cortisol is the main glucocorticoid secreted by the adrenal cortex and is involved in the stress response. Synergies between cortisol reactivity and testosterone were reported to influence antisocial behavior in young adolescence. The youth with high diurnal testosterone, combined with high or moderate cortisol reactivity, were significantly higher on antisocial behavior and attention behavior problems (29).

In addition to the common metabolic features during adolescence and the youth stage, boys and girls also showed specific metabolic characteristics. During adolescence, spermidine biosynthesis was higher in boys. One of the involved metabolites was 5-methylthioadenosine, a byproduct of polyamine synthesis in DNA turnover cycles that increases with inflammation to modulate cellular stress. It has been shown to influence the regulation of gene expression, proliferation, differentiation, and apoptosis (30). Higher serum levels of 5-methylthioadenosine have been reported in youth boys when compared to girls (31). A direct association between 5-methylthioadenosine and high metabolic risk was found in boys, possibly driven by proinflammatory pathways (32). While in adolescent girls, fatty acid oxidation and biosynthesis showed high activity. Acylcarnitines showed the highest level, indicating high activity of carnitine acetyltransferase in mitochondria (33). These results indicated the preferred metabolic fuel-from fatty acid oxidation in adolescent girls (33).

In the youth males aged 20 to 30 years, linoleic acid metabolites showed higher levels in addition to steroid metabolism. The involved metabolite eicosapentaenoic acid serves as the precursor for prostaglandin-3. It could enhance the production of the cytoprotective prostanoid 15d-PGJ2 (34), which corresponds to the elevated prostaglandin production in youth males (35). In contrast to youth males, arachidonic acid metabolites showed higher levels in youth females. It was reported that in females, arachidonic acid metabolism could rescue anti-inflammatory and butyrate-producing microbiota, then upregulate GPR41 and GPR109A and control hypothalamic inflammation (36).

#### Metabolomic characteristics during the middle and old stages

During the middle age stage, males and females showed the most significant gender differences. In males, the main metabolic features were vitamin B6 and purine metabolism. 8-Hydroxy-7-methylguanine, a methylated purine base, showed higher levels in males. High methylated purine bases were found in tumor-bearing patients compared to healthy controls (37). Urine alkylated purines were partly derived from covalently bound adducts in DNA formed from exposure to carcinogenic alkylating agents (38). Purine disorders may be associated with serious, sometimes life-threatening consequences. In females, the metabolism pathway of steroidogenesis and caffeine metabolism showed high activity. Menopausal symptom are an unavoidable problem in females during this period, which could contribute to some metabolic disorders. Caffeine metabolism was reported to be associated with menopausal symptoms, particularly vasomotor symptoms (39).

For the stage aged above 50 years, the gender difference decreased. During this period, organ metabolism activity gradually slows down. Energy-supply metabolism pathways, such as fatty acids and amino acids, showed low levels. In contrast, steroidogenesis, caffeine and pyrimidine metabolism showed high levels. Higher levels of pyrimidine metabolites in the old stage have adverse effects on health (38). Additionally, several cognitive impairment-related metabolites, including acetylhistidine and steroid hormones, were found to be higher in the old population, partly contributing to the high incidence of cognitive impairment diseases at older ages (40).

## Conclusion

In conclusion, we provide a comprehensive view of human metabolism status across the life span. To our knowledge, this is the first systematic study to analyze metabolism characterization based on population across a considerable age range. This study showed that gender differences existed from early life stages, and these differences were much smaller than those in adults. Age is another recognized confounder. Metabolism characteristics for each age group could reflect the metabolism status during different life stages and possibly contribute to some age-dependent disease incidences. Our present study would be helpful to understand the age- and gender-dependent metabolism differences, which will be a critical component for the development of metabolomics-based systems biology as a population screening and precision medicine research.

Additionally, several limitations still exist and need to be further validated in the future. First, urine samples of children and adults were collected from two different hospitals. Although sampling operation is strictly controlled, batch effects resulting from center differences could not be completely eliminated. Considering batch effects, data from children and adults were first analyzed separately and then a cross comparison was performed, resulting in an age gap between second school and the youth stages. Second, in the children population, each stage could show specific features for their rapid development. However, due to sample size limitations, we provide an overview of the average metabolism status over a period of 6 years. Third, the influences of diets and circadian rhythm on urine metabolomics could not be completely eliminated, though all subjects were from the same region. For future validation analysis, these influences should be systematically evaluated and analyzed.

## Methods

### Study sample population

Our study enrolled 348 children aged 1 to 18 years from Beijing Children’s Hospital and 315 adults aged 20 to 70 years from 3rd Clinical Hospital, Jilin University (Table 1). According to physiological developmental characteristics, the enrolled population was divided into 6 age stages for males (boys) and females (girls): preschool (aged 1-6), primary school (aged 7-12), secondary school (aged 13-18), youth (aged 20-30), middle (aged 31-50) and old (aged >50). Participates were checked and examined by trained nurses according to standard operating procedures. All physical examination indexes were in the normal range. On the day of the examinations, urine samples were collected at approximately 7:00 and 9:00 a.m. on an empty stomach. This study was approved by the Ethics Committee of Peking Union Medical College. A doctor informed the eligible participants about the nature of the study and invited them to participate. All human subjects provided informed consent before participating in this study. The research was carried out according to The Code of Ethics of the World Medical Association (Declaration of Helsinki). and institutional review board of Peking Union Medical College has approved the study.

**Table 1.** Basic characteristics of the participants in this study.

| Age stage | Female | Male | Total |
| --- | --- | --- | --- |
| 1-6 | 54 | 59 | 113 |
| 7-12 | 59 | 57 | 116 |
| 13-18 | 59 | 60 | 119 |
| 20-30 | 38 | 43 | 81 |
| 31-50 | 72 | 68 | 140 |
| >50 | 54 | 40 | 94 |
| Total | 336 | 327 | 663 |

**Table 2.** Metabolic characteristics in males (boys) and females (girls)

|  | **Males (boys)** | **Females (girls)** |
| --- | --- | --- |
| **1-6** | Pantothenate and CoA biosynthesis<br>Alanine metabolism<br>Pyrimidine metabolism | Pantothenate and CoA biosynthesis<br>Alanine metabolism<br>Vitamin B6 metabolism |
| **7-12** | Tryptophan metabolism | Ether lipid metabolism<br>Nicotinate and nicotinamide metabolism<br>Histidine metabolism |
| **13-18** | Fatty Acid oxidation and biosynthesis<br>Spermidine and spermine biosynthesis<br>Riboflavin metabolism | Fatty acid oxidation and biosynthesis |
| **20-30** | Androgen and estrogen metabolism<br>Steroidogenesis<br>Linoleic acid metabolism | Androgen and estrogen metabolism<br>Steroidogenesis<br>Arachidonic acid metabolism |
| **30-50** | Vitamin B6 metabolism<br>Purine metabolism | Steroidogenesis<br>Caffeine metabolism |
| **>50** | Vitamin K metabolism<br>Steroidogenesis<br>Caffeine metabolism | Vitamin K metabolism<br>Pyrimidine metabolism |

### Urine sample preparation

Urine samples were processed according to our previous protocols. Briefly, proteins from urine samples (200 μL) were precipitated using acetonitrile (400 μL). The mixture was vortexed for 30 sec and centrifuged at 14,000 g for 10 min. The supernatant was dried under vacuum and then reconstituted with 200 μL 2% acetonitrile/water. Quality control (QC) samples were prepared by mixing aliquots of 50 representative samples across different groups to be analyzed. All samples were randomly injected into the LC-MS system within a single analysis (within 12 days). QC sample was injected every ten or twelve samples.

### Urine metabolite LC-MS measurements

HRLC-MS was selected for urinary metabolite detection due to its high sensitivity and reproducibility. Urine metabolite separation and analysis was conducted using a Waters ACQUITY H-class LC system coupled with a LTQ-Orbitrap Velos Pro mass spectrometer (Thermo Fisher Scientific, MA). The following 18-min gradient on a Waters HSS C18 column (3.0 × 100 mm, 1.7 μm) at a flow rate of 0.5 mL/min was used: 0-1 min, 2% solvent B (mobile phase A: 0.1% formic acid in H_2_O; mobile phase B: acetonitrile); 1-3 min, 2-55% solvent B; 3-8 min, 55-100% solvent B; 8-13 min, 100% solvent B; 13-13.1 min, 100-2% solvent B; 13.1-18 min, 2% solvent B. The column temperature was set at 45°C.

All samples were untargeted scanned from 100 to 1000 m/z at a resolution of 60 K at the positive ESI mode. The automatic gain control (AGC) target was 1 × 10^6,^ and the maximum injection time (IT) was 100 ms. The extracted MS features were divided into several targeted lists and imported to MS2 method for targeted data dependent analysis. MS/MS fragment acquisition was acquired at a resolution of 15 K with an AGC target of 5 × 10^5^. Collision energy was optimized as 20, 40 or 60 for each targeted list with higher-energy collisional dissociation (HCD) fragmentation. The injection order of urine samples was randomized to reduce any experimental bias. The QC sample was injected regularly to monitor system stability.

### Statistical analysis

Raw data files were processed by Progenesis QI (Waters, Milford, MA) software based on a previously published identification strategy, which included sample alignment, peak picking, peak grouping, deconvolution and final information export (8). The exported data were further preprocessed by MetaboAnalyst 3.0 (http://www.metaboanalyst.ca), which included missing value estimation, log transformation and Pareto scaling. Variables that were missed in 50% or greater of the samples were removed from further statistical analysis.

Nonparametric tests (Wilcoxon rank-sum tests and Kruskall-Wallis tests) were used to evaluate the significance of variables related to gender and age using MetaboAnalyst 3.0. Benjamini-Hochberg correction was applied throughout to account for multiple test comparisons. Cutoff of FDR 0.05 was applied. Pattern recognition analysis (principal component analysis [PCA]; partial least squares discrimination analysis [PLS-DA]; and orthogonal partial least square analysis [OPLS-DA]) was carried out using SIMCA 14.0 software (Umetrics, Sweden) to visualize group classification and select significant features. 100 permutation tests were used to validate the OPLS-DA and PLS-DA model to avoid over-fitting of the model. Significantly differential metabolites were chosen according to the following criteria: (i) adjusted p <0.05; (ii) fold change between two groups >2; and (iii) the variable importance plot (VIP) value obtained from OPLS-DA was above 1. Heat map virtualization and metabolic pathway enrichment analysis was performed by MetaboAnalyst 3.0, and the enrichment results were visualized using R based on previous methods (9).

### Feature annotation and metabolite identification

MS1 features were divided into several targeted lists and imported to MS method for targeted data dependent analysis. The MS/MS spectra were further imported to Progenesis QI for database searching (HMDB: http://www.hmdb.ca/) and MS/MS spectra matching using “MetFrag” algorithm (10). Detailed compound identification information (.csv file) included compound ID, adducts, formula, score, MS/MS score, mass error (in ppm), isotope similarity, theoretical isotope distribution, web link, and m/z values. Confirmation of the differential compounds was performed by the parameters, including Score, Fragmentation score, and Isotope similarity given by Progenesis QI. Score ranging from 0 to 60, is used to quantify the reliability of each identity. According to the score results of the reference standards, the threshold was set at 35.0. Isotope similarity is calculated by comparison of the measured isotope distribution of a precursor ion with the theoretical. The compound identification is more reliable the higher the values obtained. For metabolites with biological significance, the annotations were further manually validated by standards MS2 spectra or MS2 spectra database (Masbank: http://www.massbank.jp/), combined with fragmentation explanation.

## Abbreviations

LC-HRMS: liquid chromatography coupled with high resolution mass spectrometry
NMR: nuclear magnetic resonance
GC-TOF-MS: gas chromatography-time-of-flight mass spectrometry
TCA: tricarboxylic acid
CoA: coenzyme
QC: quality control
PCA: principal component analysis
PLS-DA: partial least squares discrimination analysis
OPLS-DA: orthogonal partial least square discrimination analysis
VIP: variable importance plot
HMDB: human metabolome database

## Authors’ contributions

XYL, XYT interpret the data and wrote the manuscript, QHS, NR and FJ collected the samples and designed the experiments, WS, WQS and CYH conceived and designed the study, helped to interpret the data and revised the manuscript. JL and XYT performed the sample preparation, ZGG and HDS helped to perform data analysis, YL and JM helped to collect the children samples. WS is the guarantor of this work and, as such, had full access to all the data in the study and takes responsibility for the integrity of the data and the accuracy of the data analysis.

## Acknowledgments

This work was supported by the National Key Research and Development Program of China (No. 2016 YFC 1306300, 2018YFC0910202), National Natural Science Foundation of China (No. 30970650, 31200614, 31400669, 81371515, 81170665, 81560121, 81572082), Beijing Natural Science Foundation (No. 7173264, 7172076), Beijing cooperative construction project (No.110651103), Beijing Science Program for the Top Young (No.2015000021223TD04), Beijing Normal University (No.11100704), Peking Union Medical College Hospital (No.2016 −2.27), CAMS Innovation Fund for Medical Sciences (2017-I2M-1-009), Capital’s Funds for Health Improvement and Research (2016-2-2096), Beijing Municipal Administration of Hospitals’ Youth Program (QML20171205), Beijing Children’s Hospital Young Investigator Program Fund (BCHYIPB-2016-01), and Biologic Medicine Information Center of China, National Scientific Data Sharing Platform for Population and Health. The funding bodies played no role in the design of the study and collection, analysis, and interpretation of data and in writing the manuscript.

## Availability of data and material

The datasets generated and/or analysed during the current study are available in the iProx repository: https://iprox.org/page/SSV024.html;url=1562292290181InPK.Password: U8b5.

## Supporting Information

### Supporting Figures

**FigS1.**
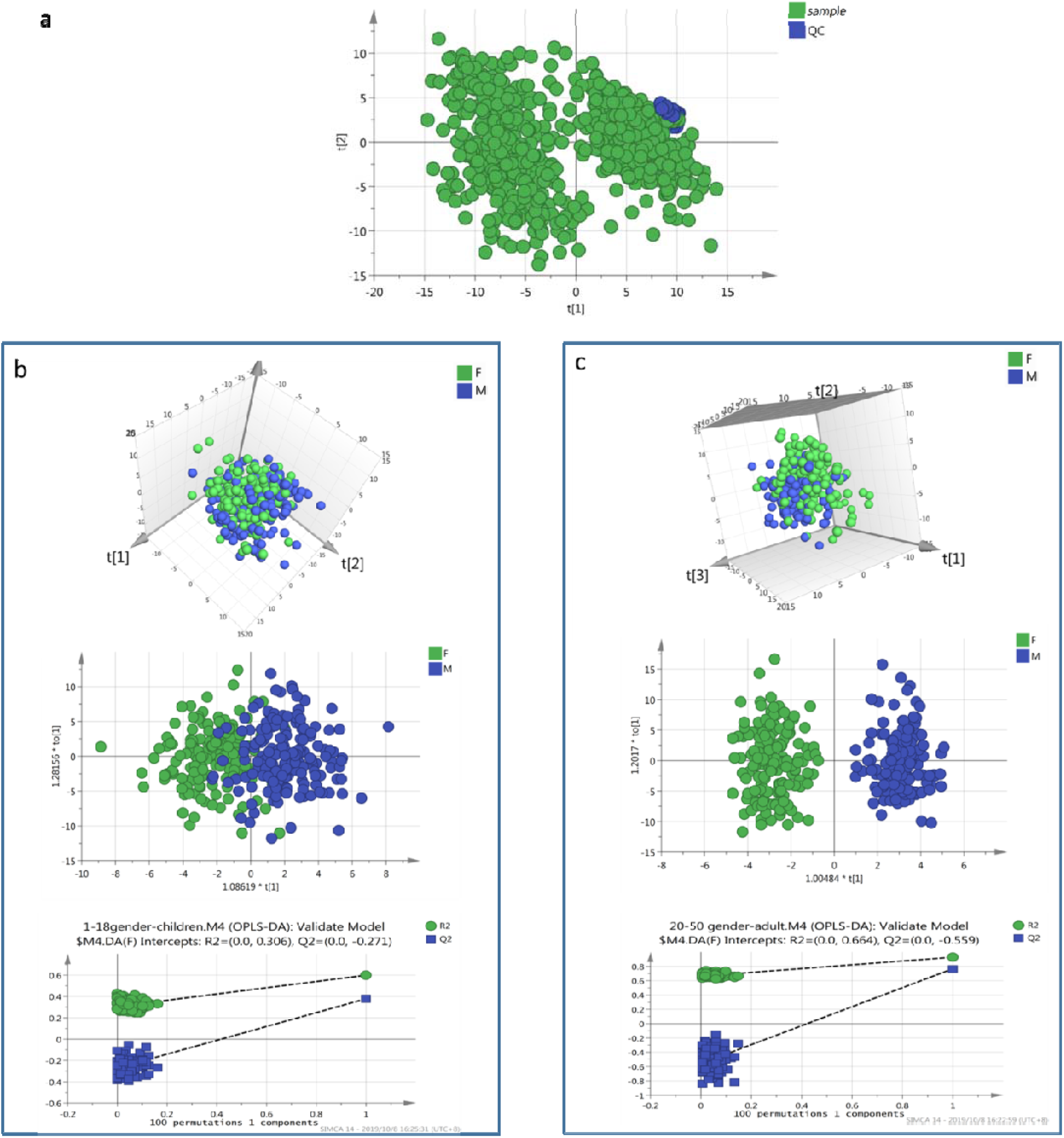
a. Score plot of all samples. Good cluster of QC samples indicated good stability of the analytical platform. Separation of adults and children samples resulted from biological or sampling center differences. b. PCA and OPLS-DA score plot of urine metabolomics between male and females in children. c. PCA and OPLS-DA score plot of urine metabolomics between male and females in adults. 100 permutation test was used to validate the OPLS-DA model. The permutation plots were given in “b” and “c”.

**Fig S2.**
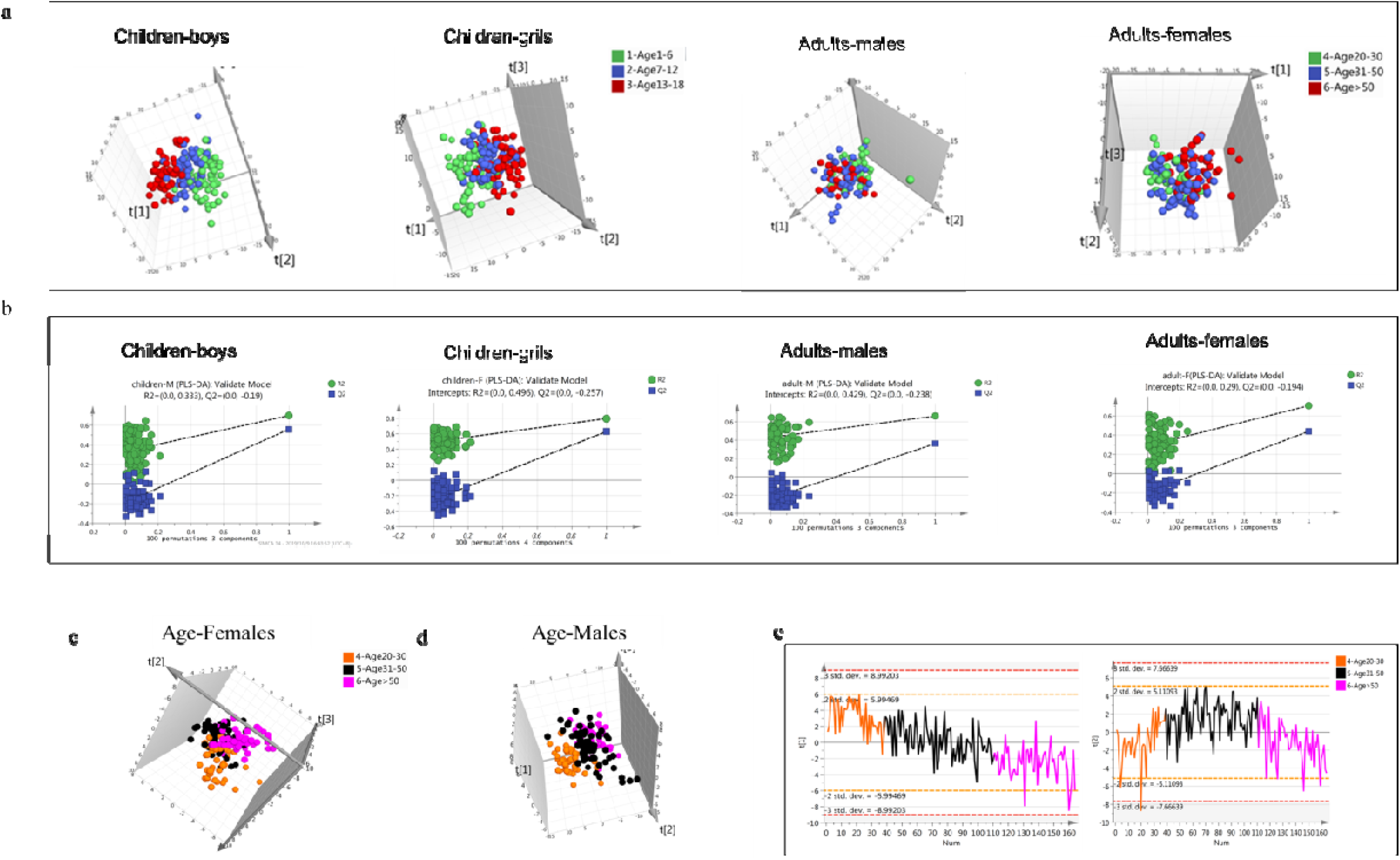
Separation of urine metabolites with different age stages in children and adults. A. PCA score plot of urine metabolomics in different age group for children-boys, children-girls, adults-males and adults-females. b. Permutation test plots of PLS-DA model based on different age group for children-boys, children-girls, adults-males and adults-females. c. PLS-DA score plot of urine metabolomics of different age in males. d. PLS-DA score plot of urine metabolomics of different age in females. e. Change trend of the first and second components of PLS-DA in adults. The first component explain the most variance of the model, thus contributing most to class separation.

**Fig S3.**
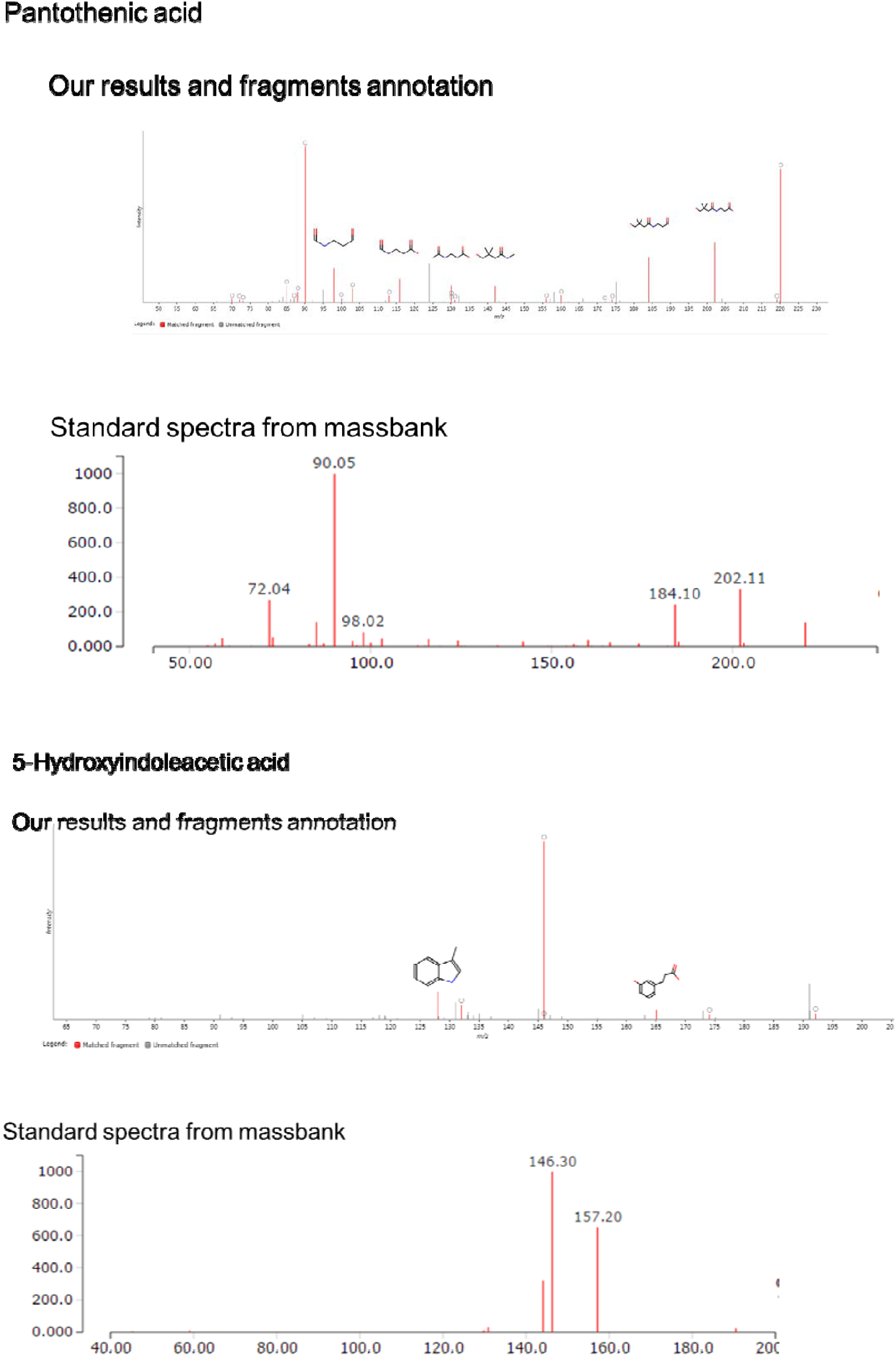
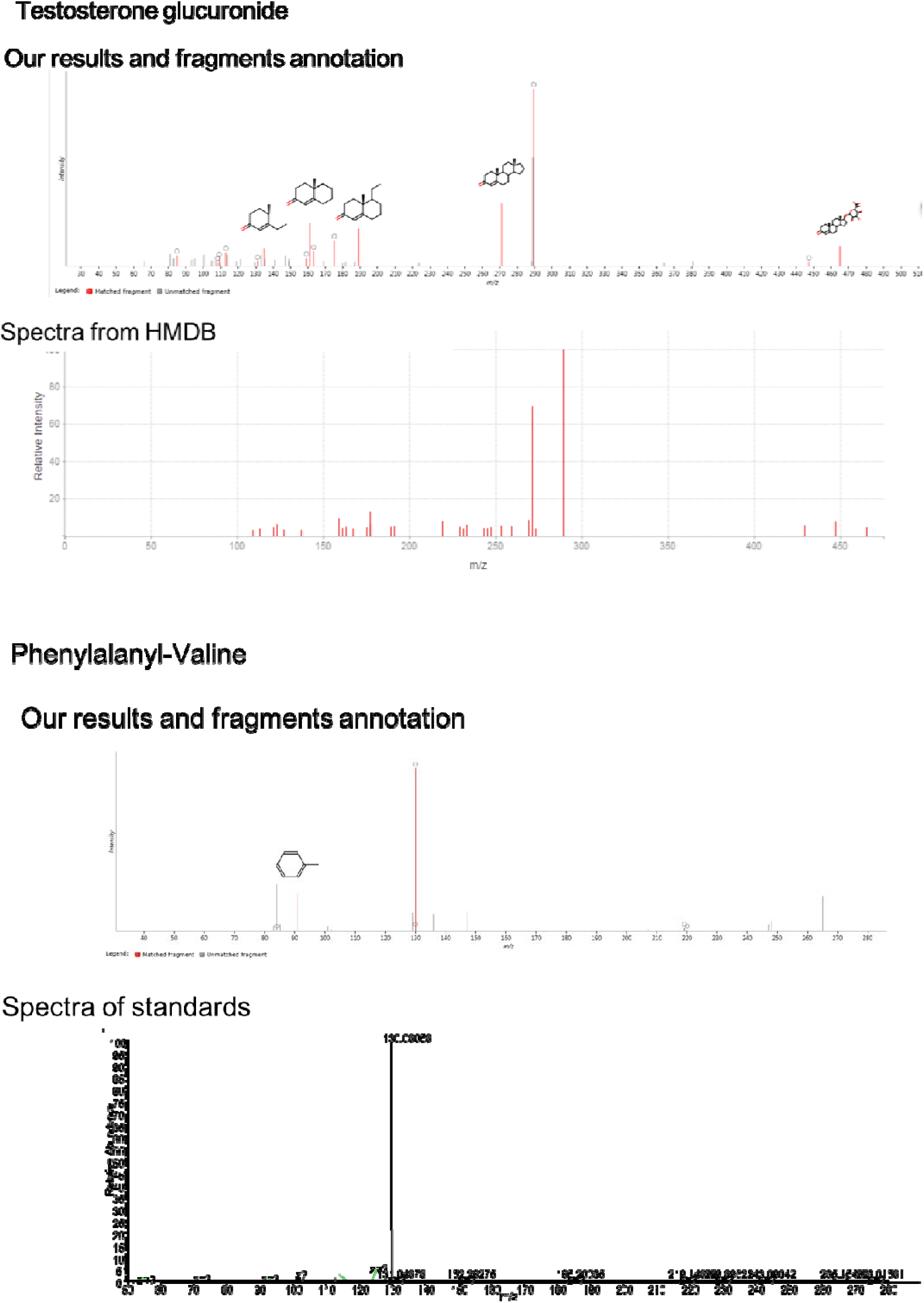
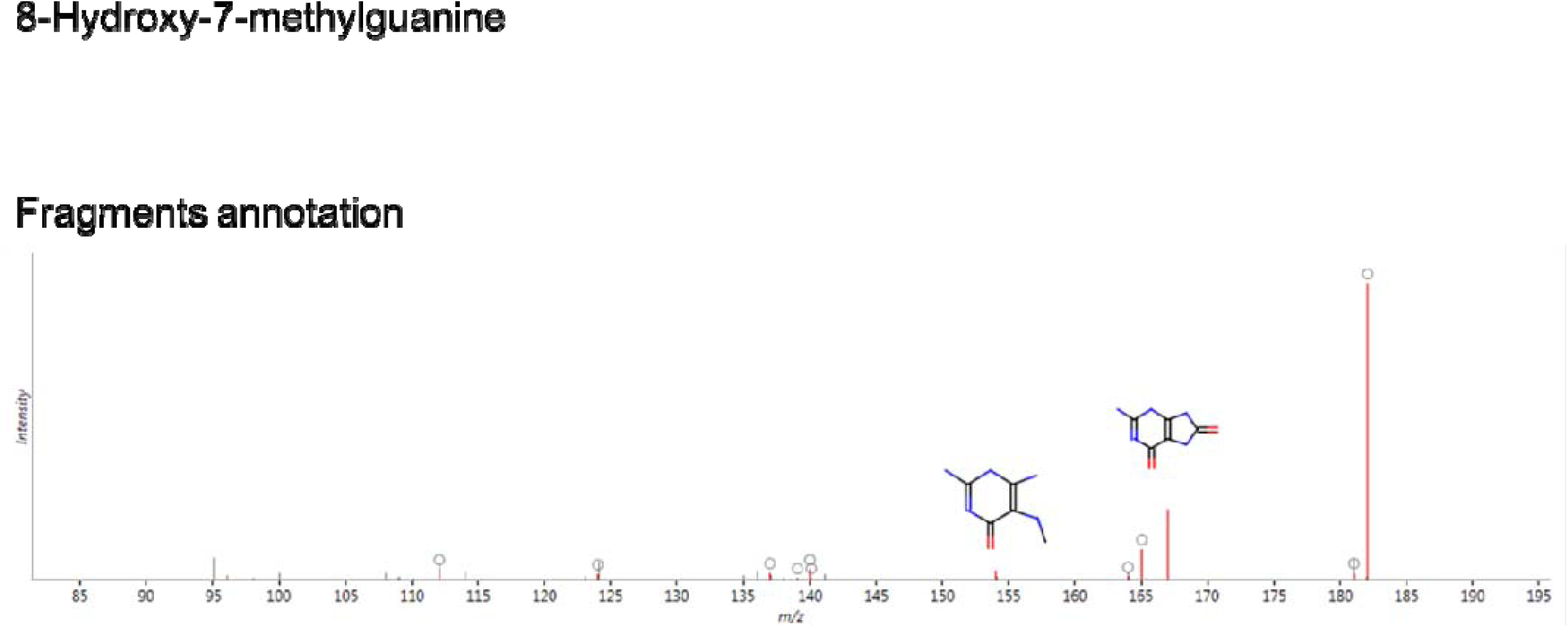
Metabolites annotation using database and standards spectra or fragments annotation

**Fig S4.**
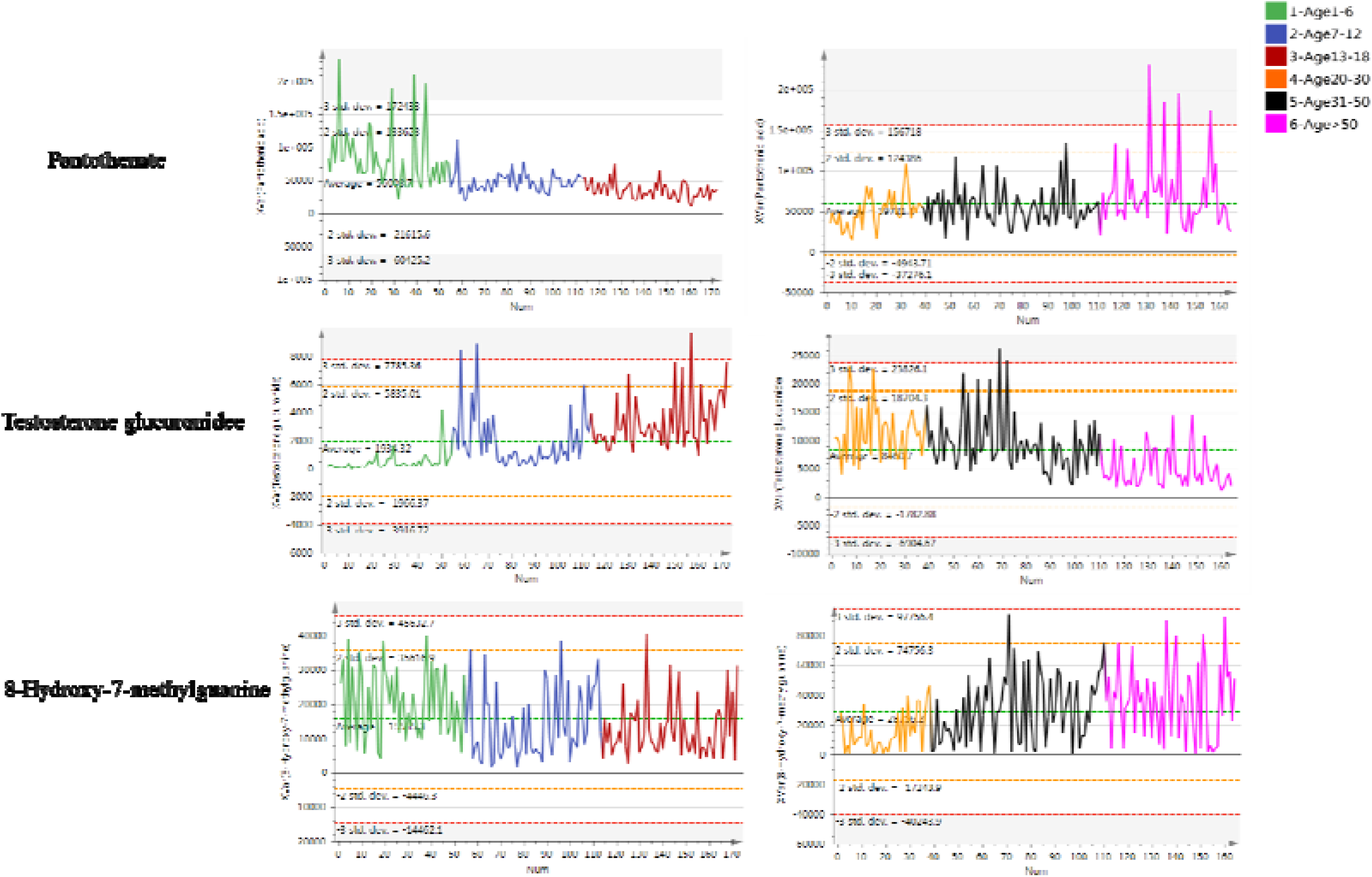
Metabolites variation with age in females (girls).

### Supporting Tables

**Table S1.**
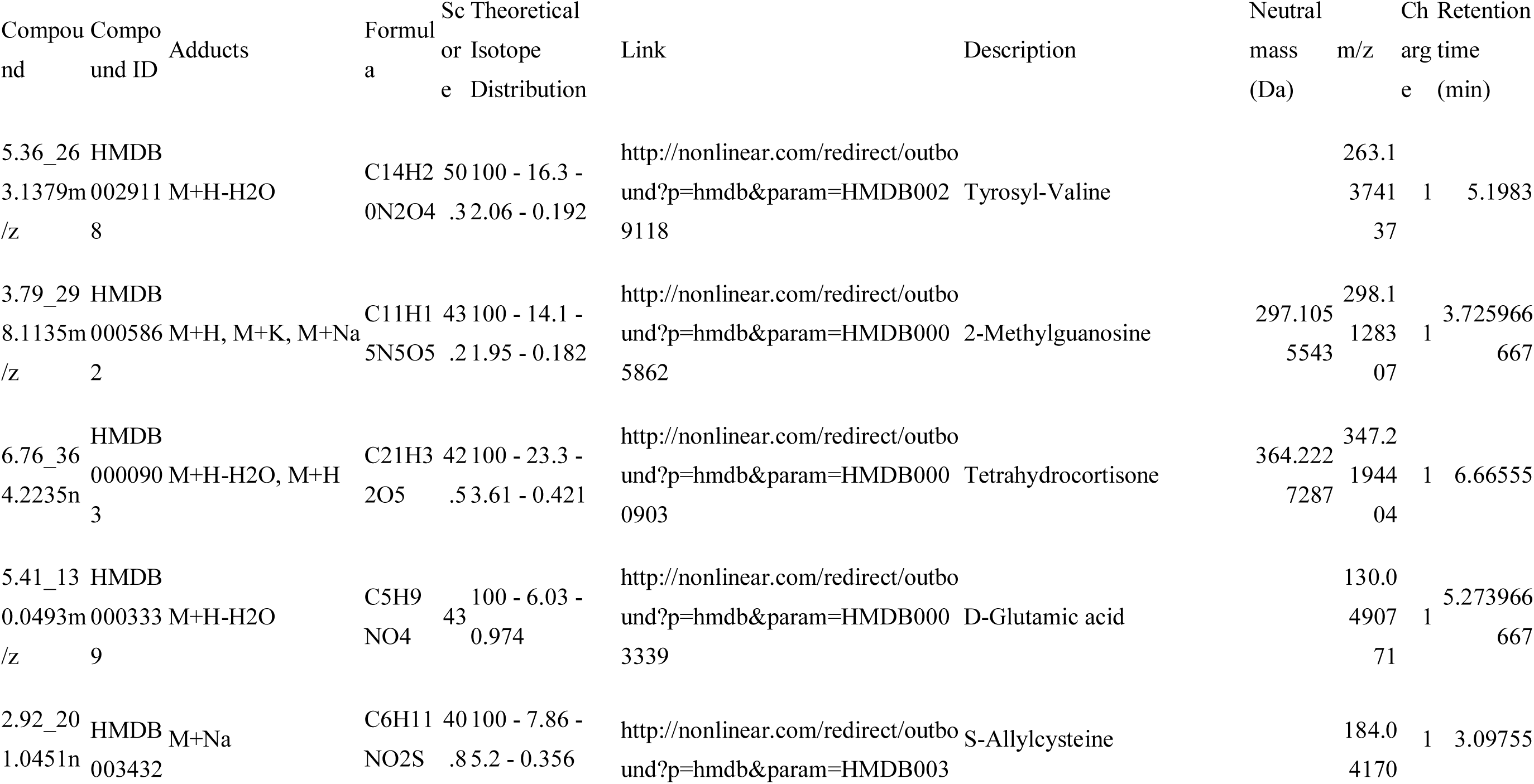
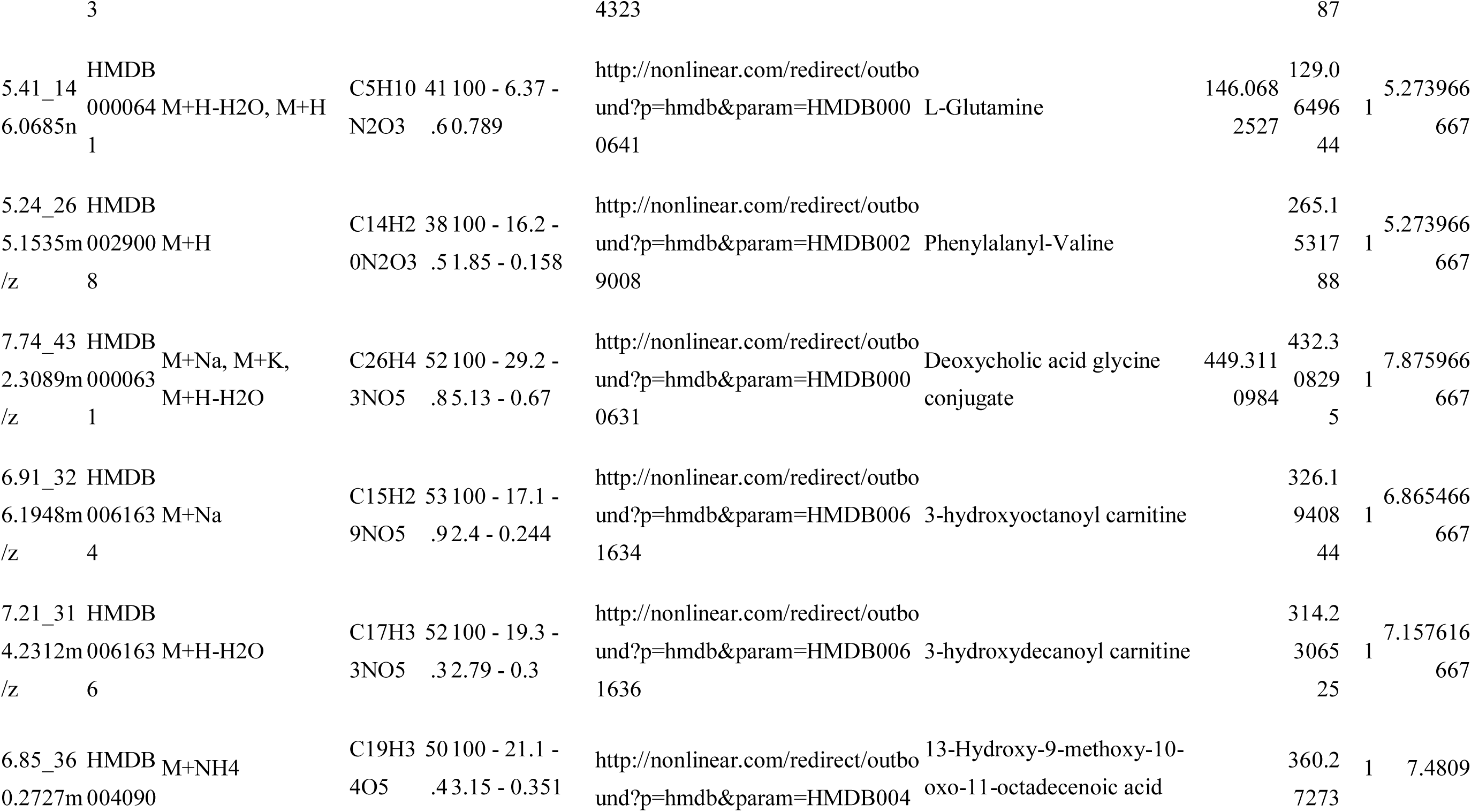
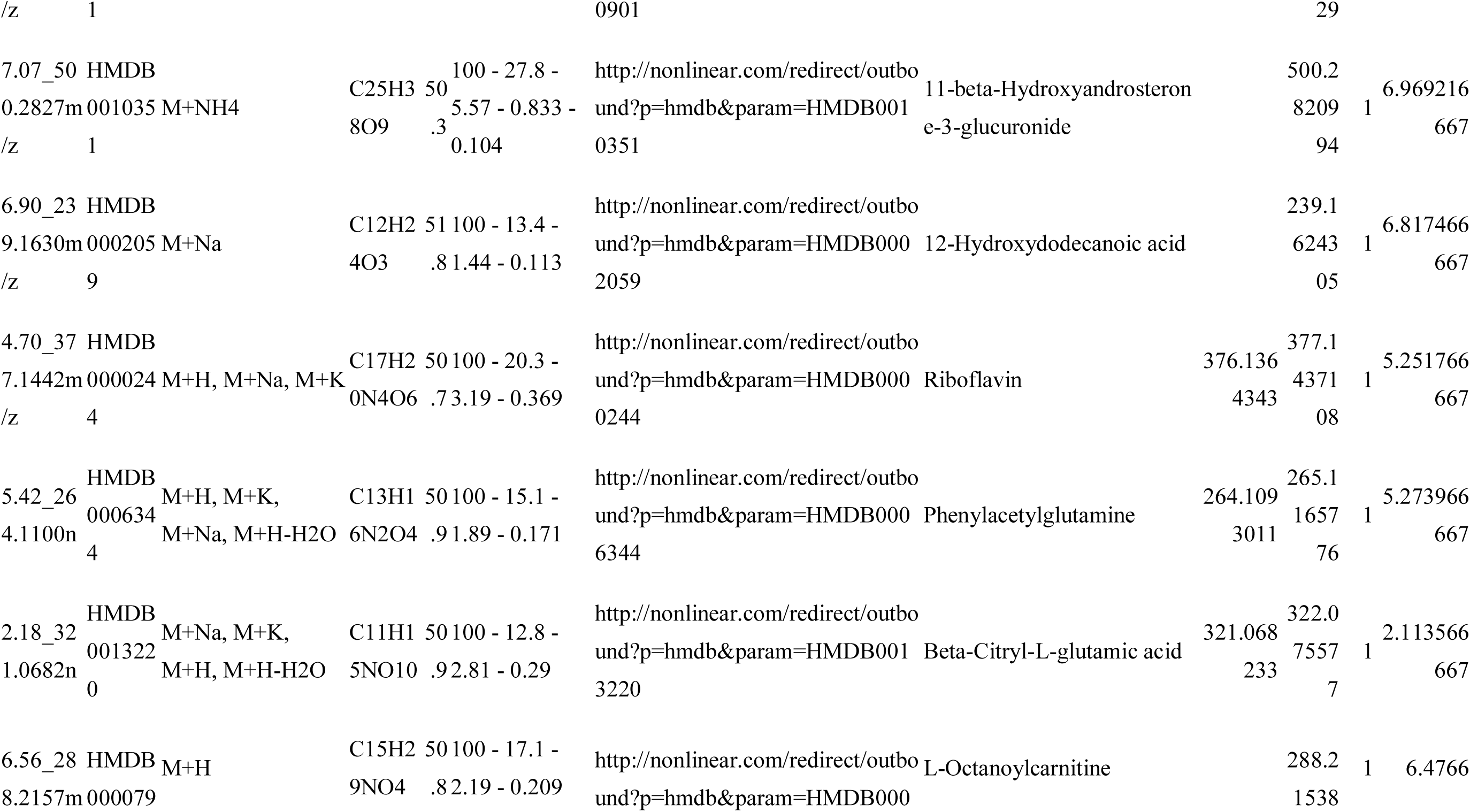
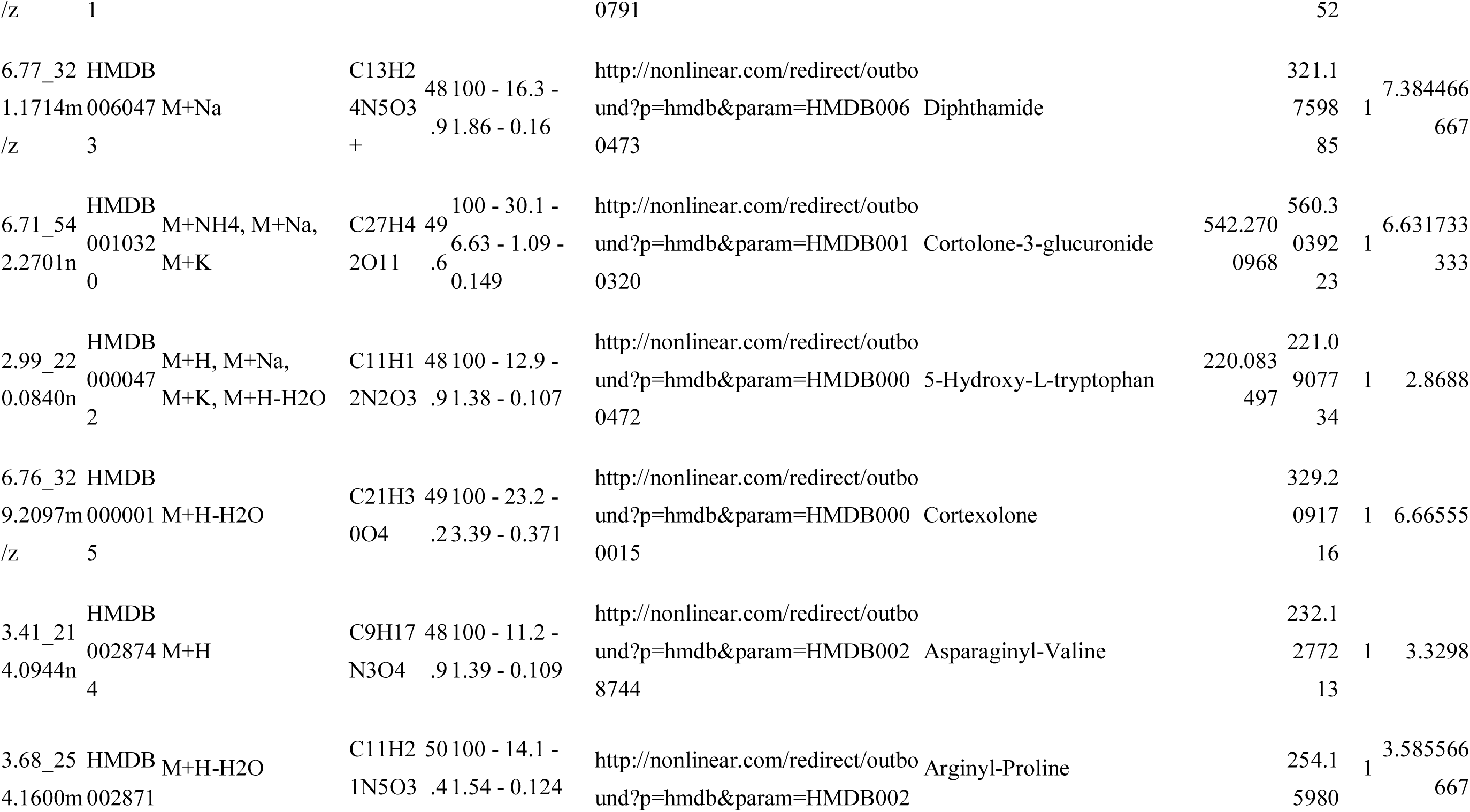
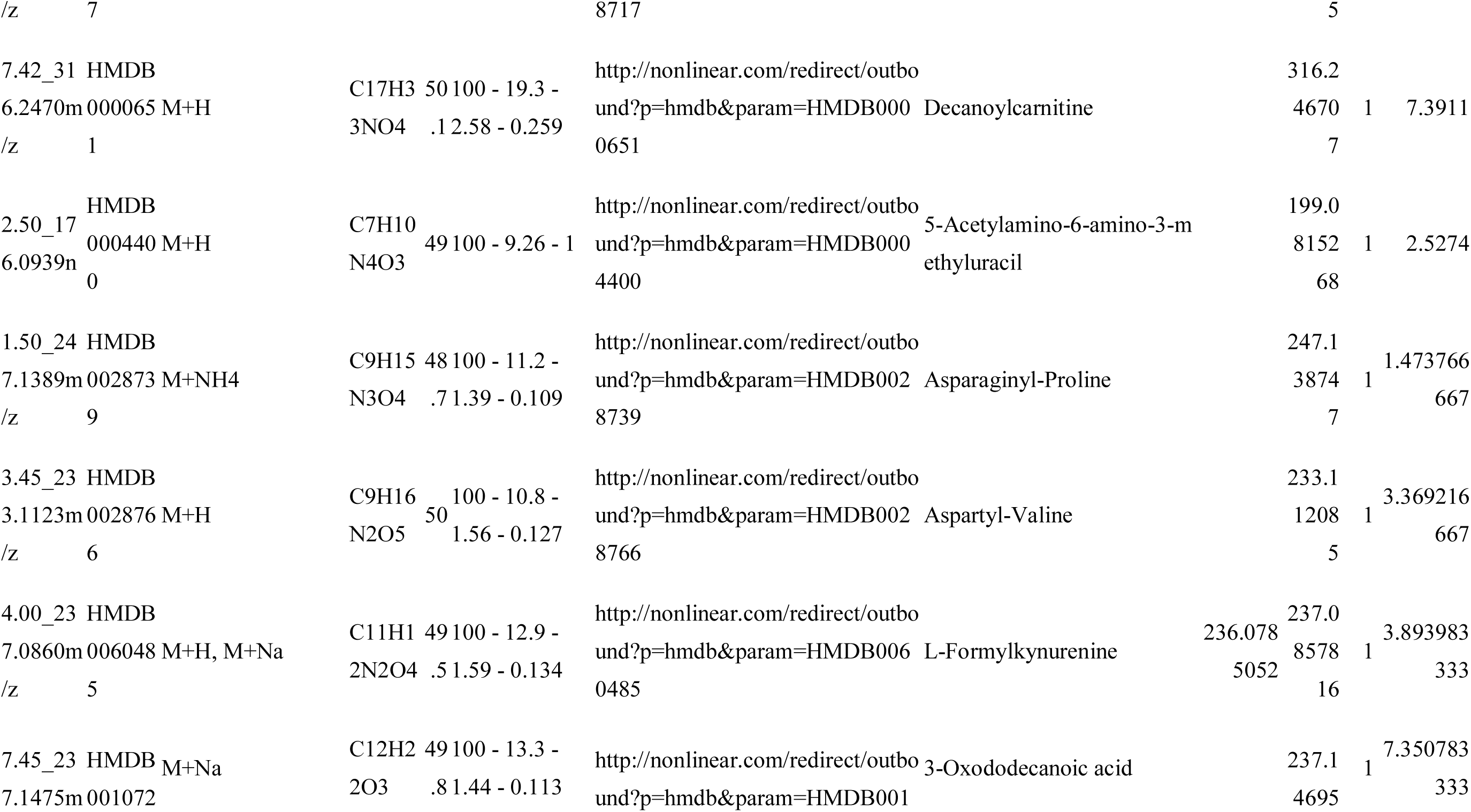
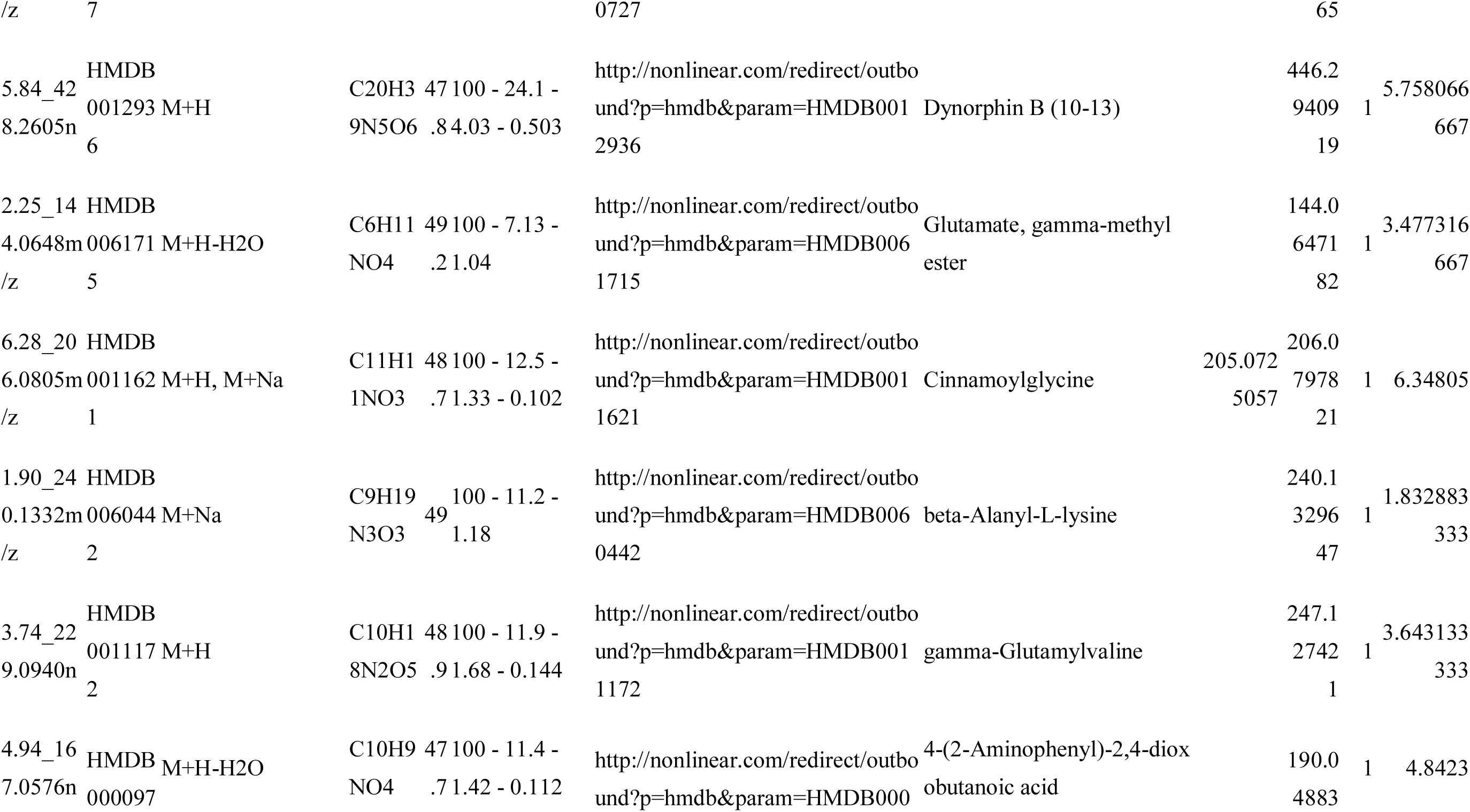
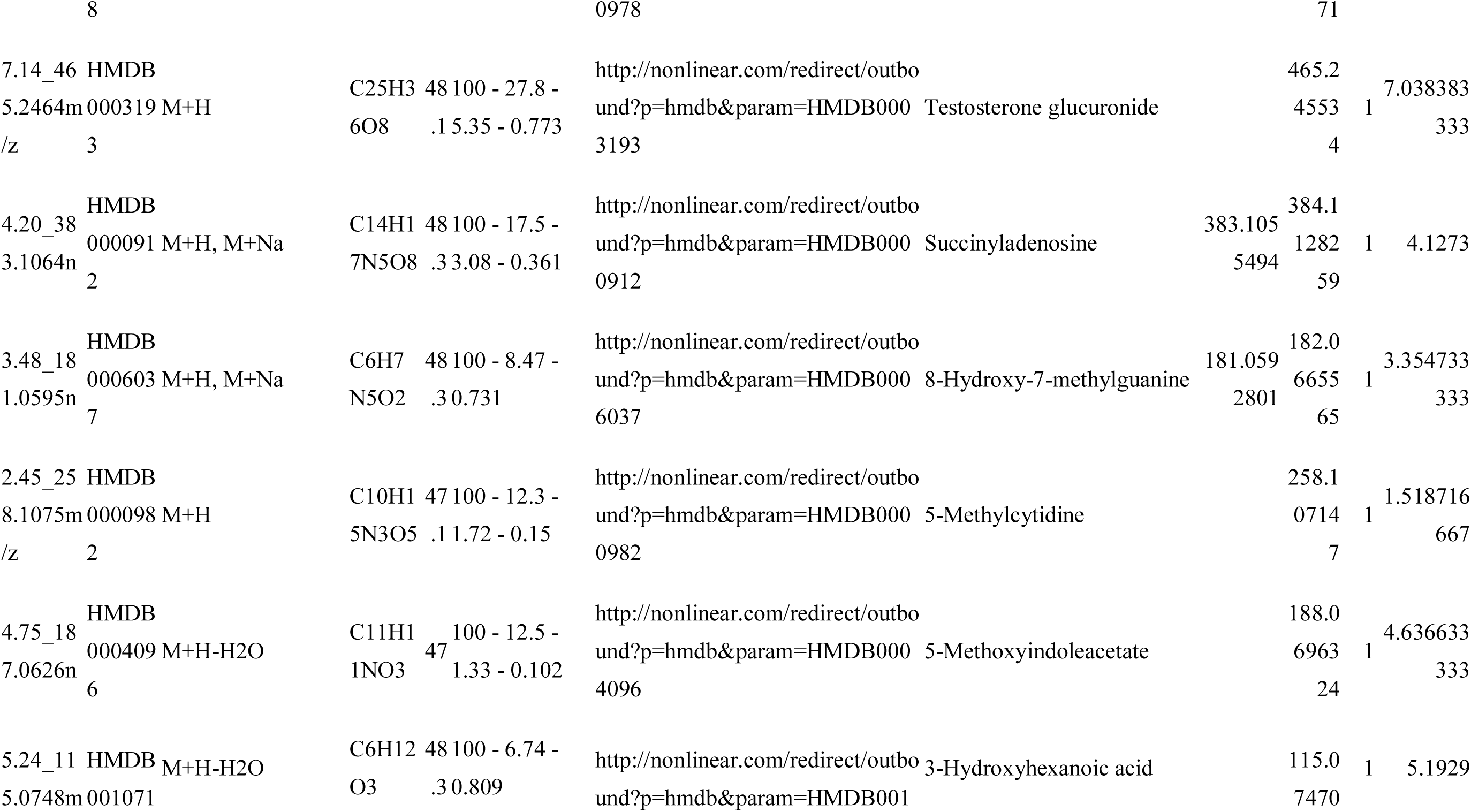
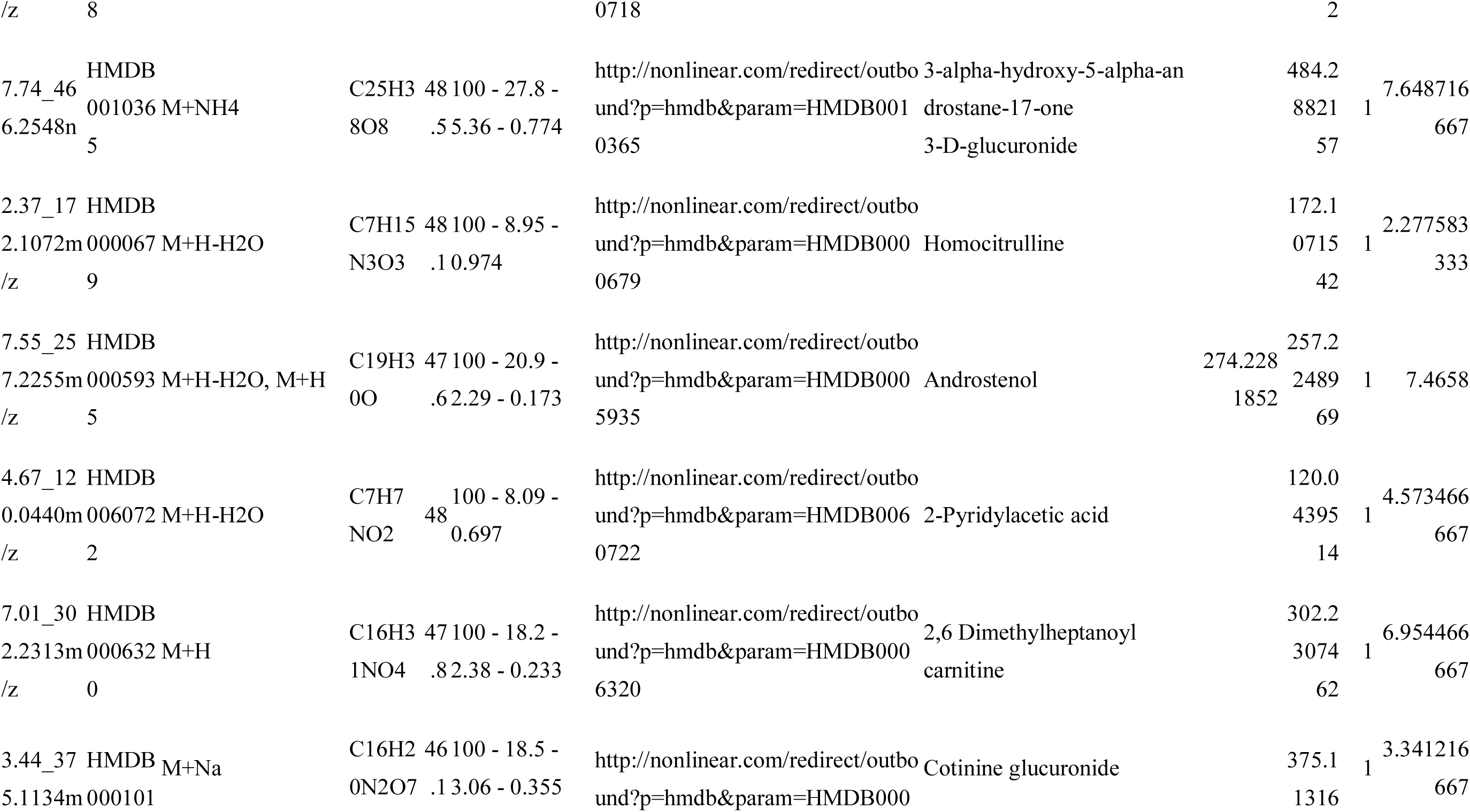
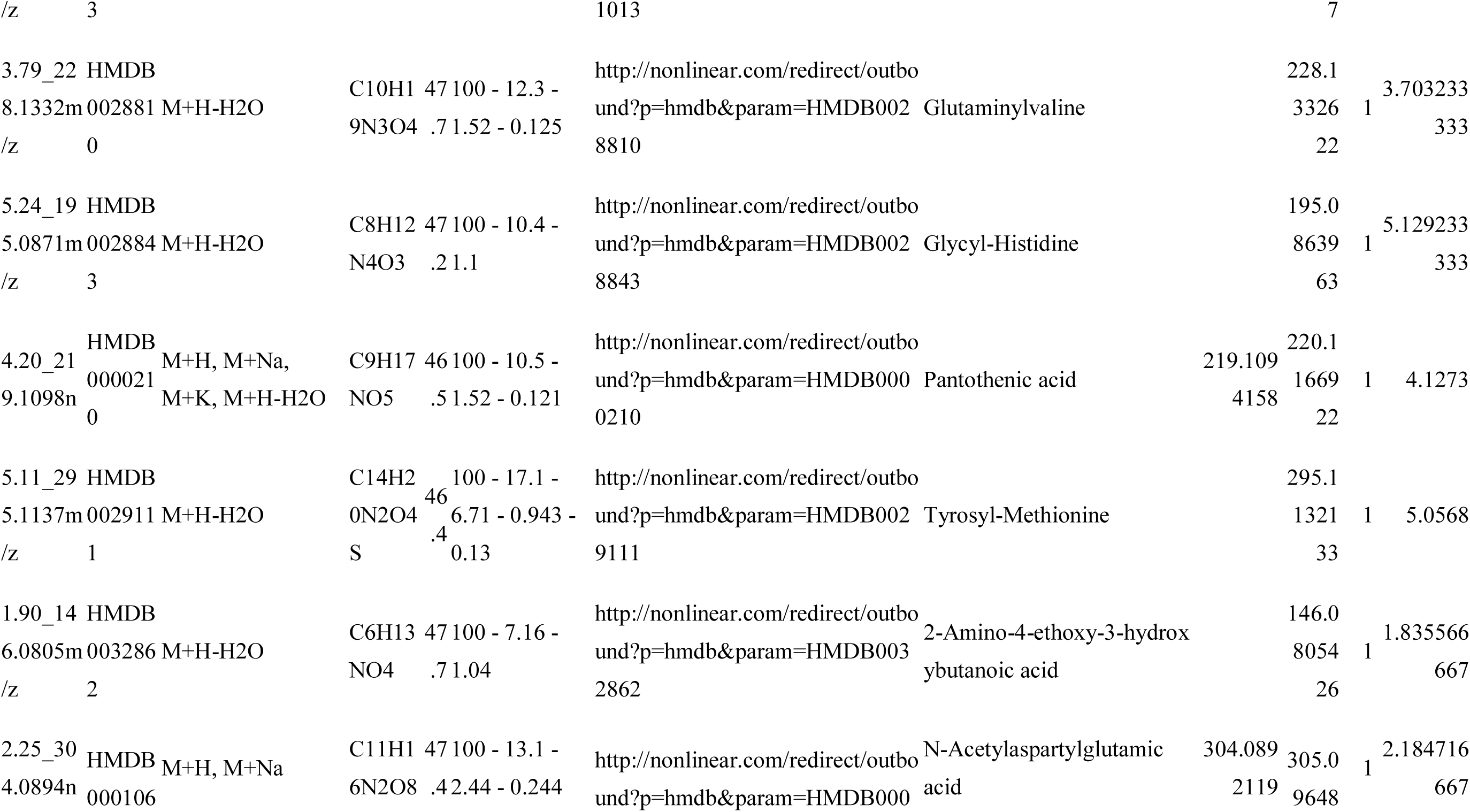
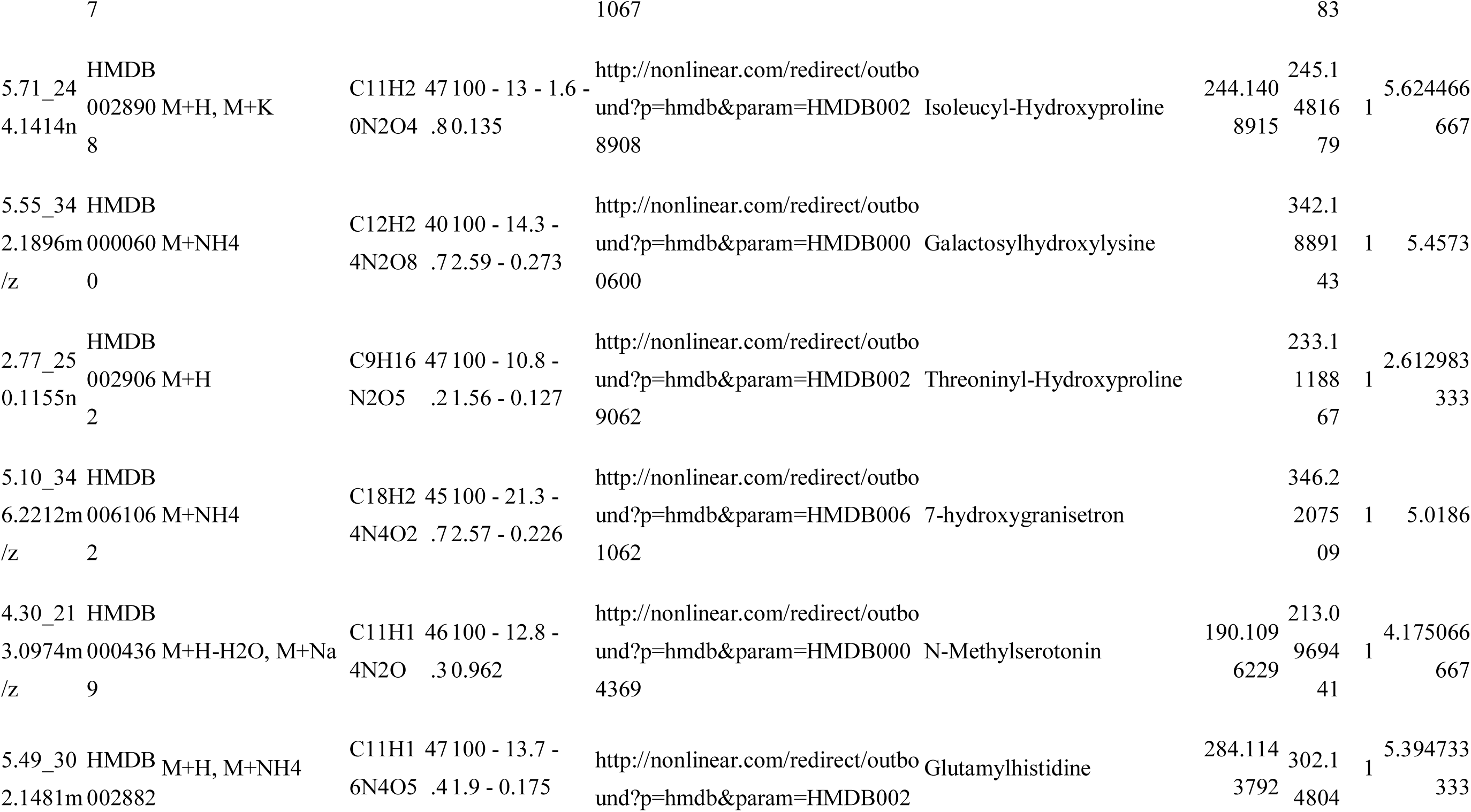
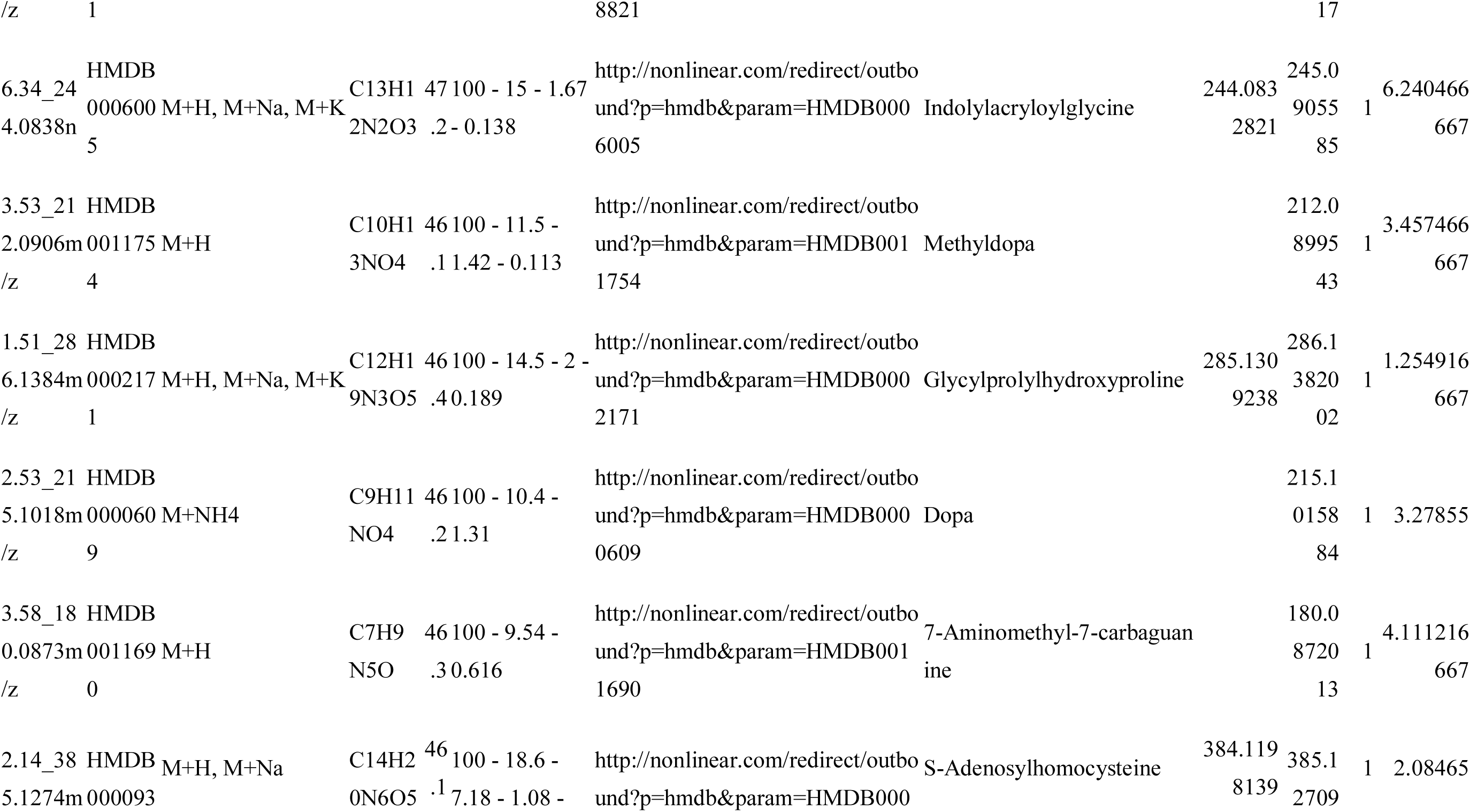
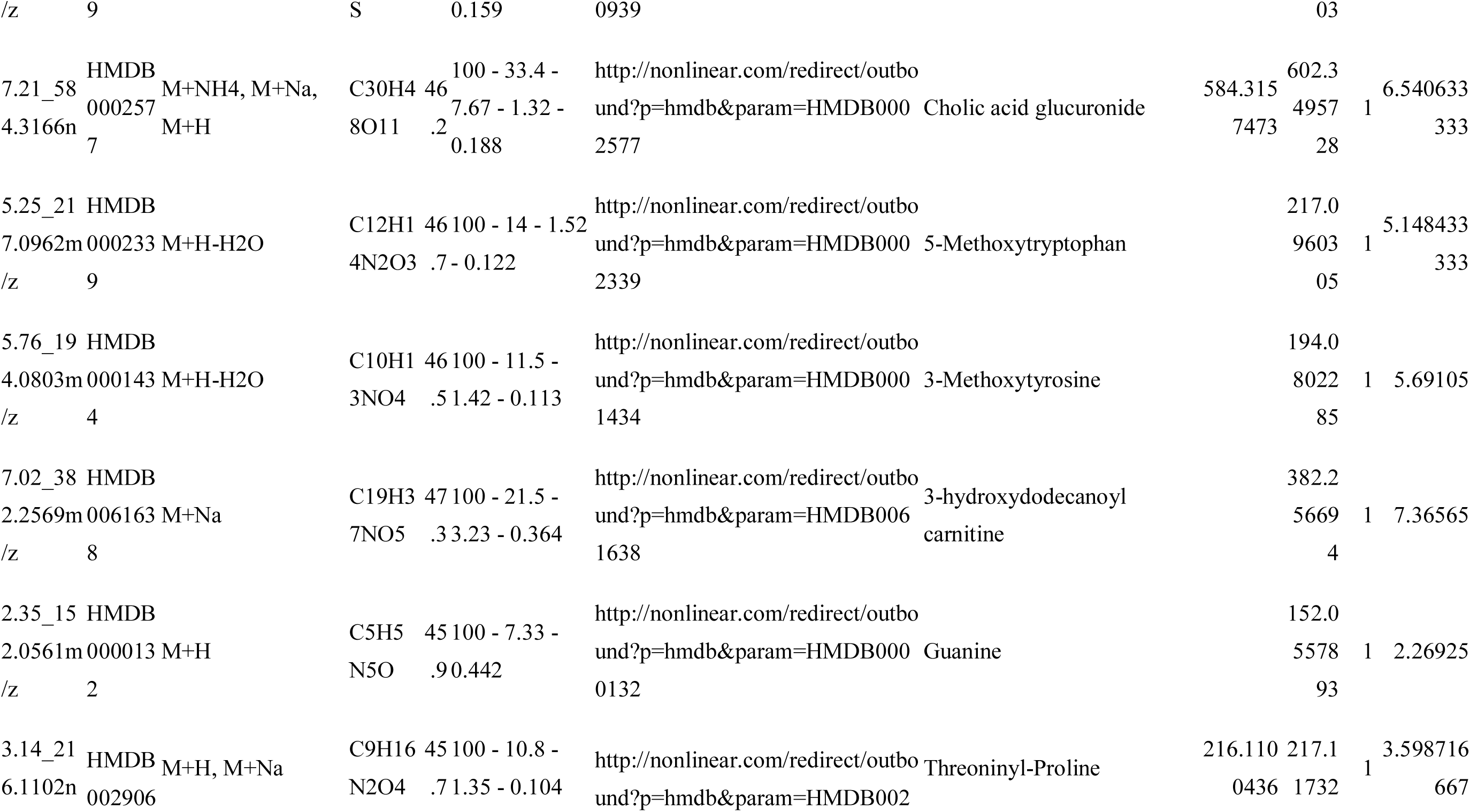
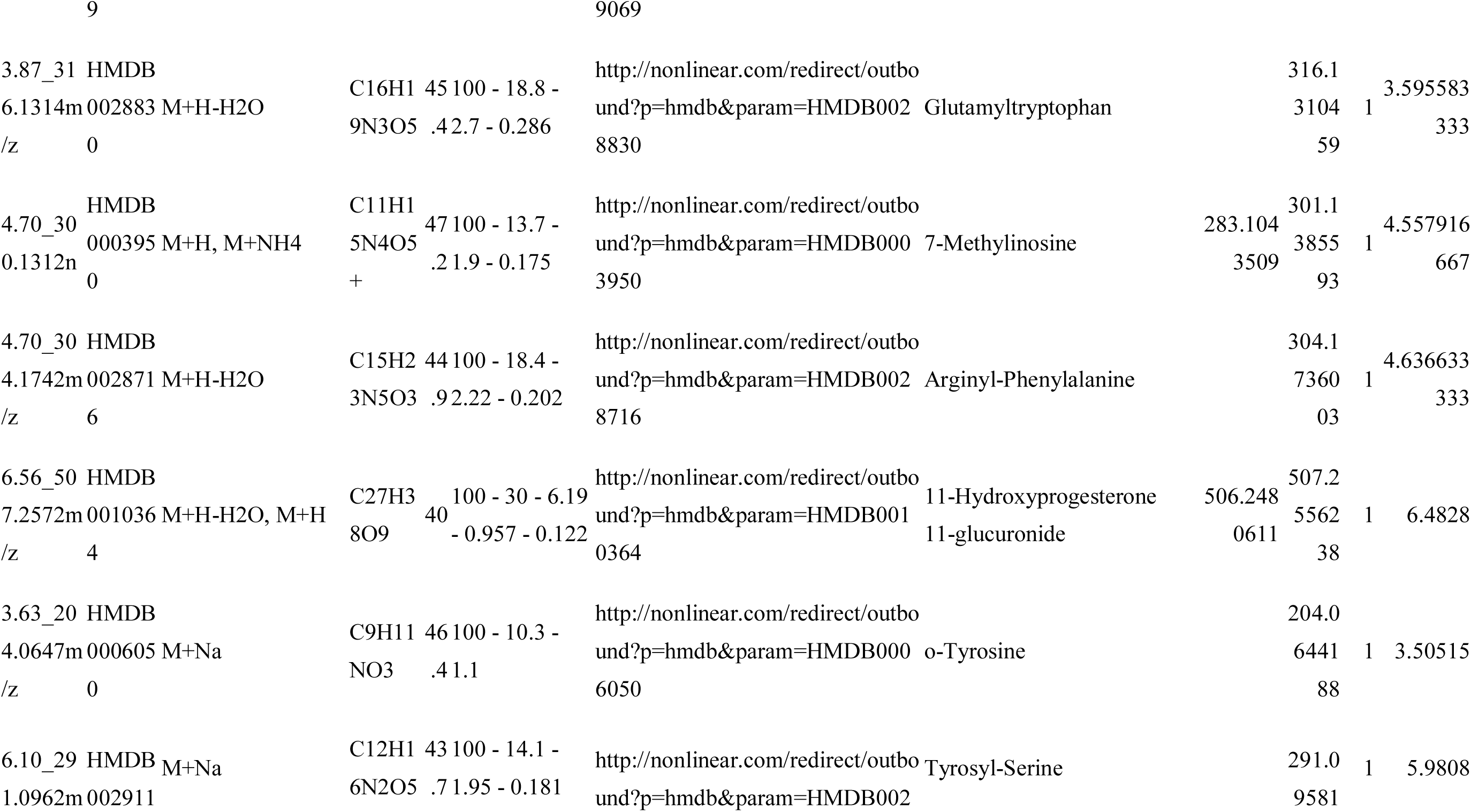
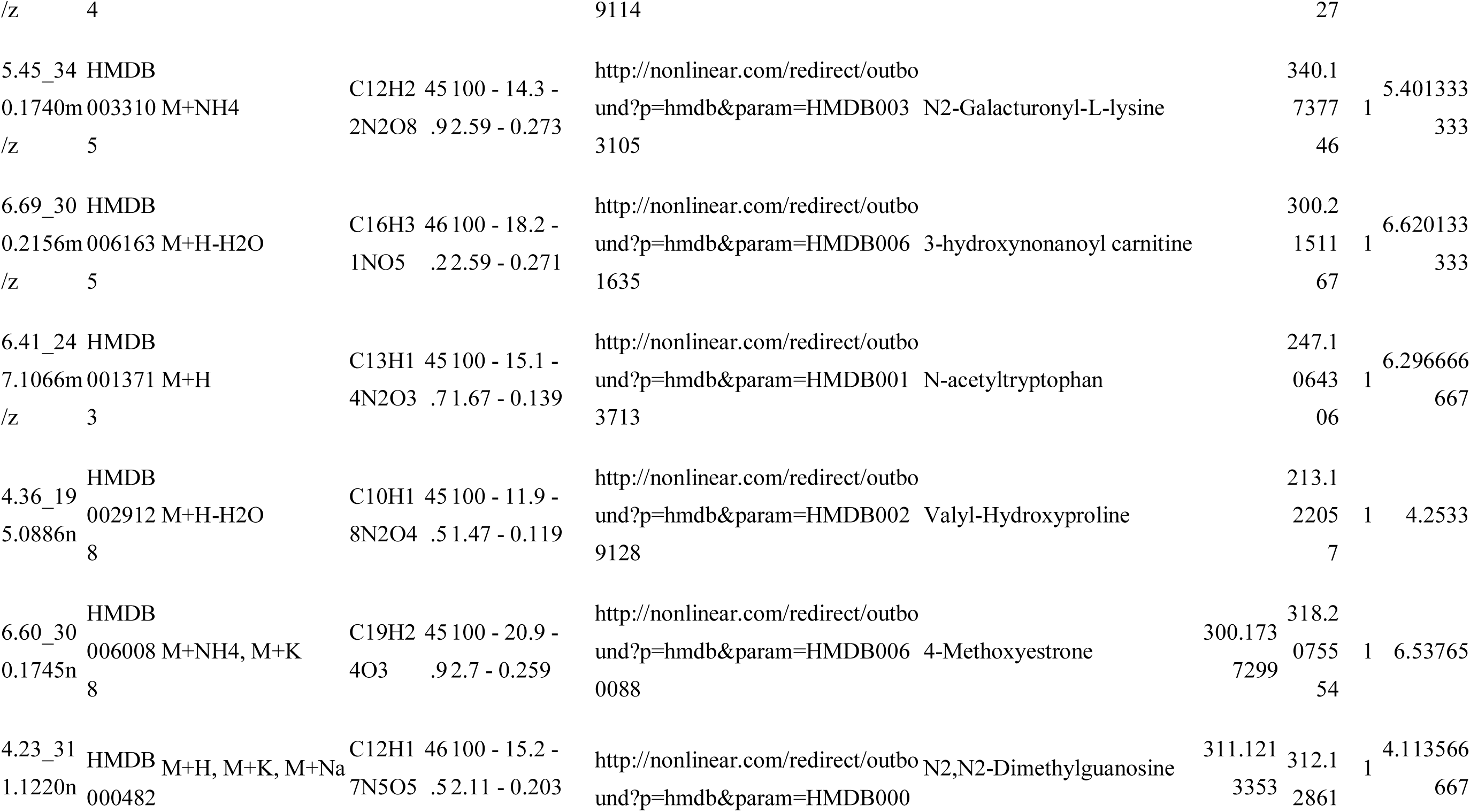
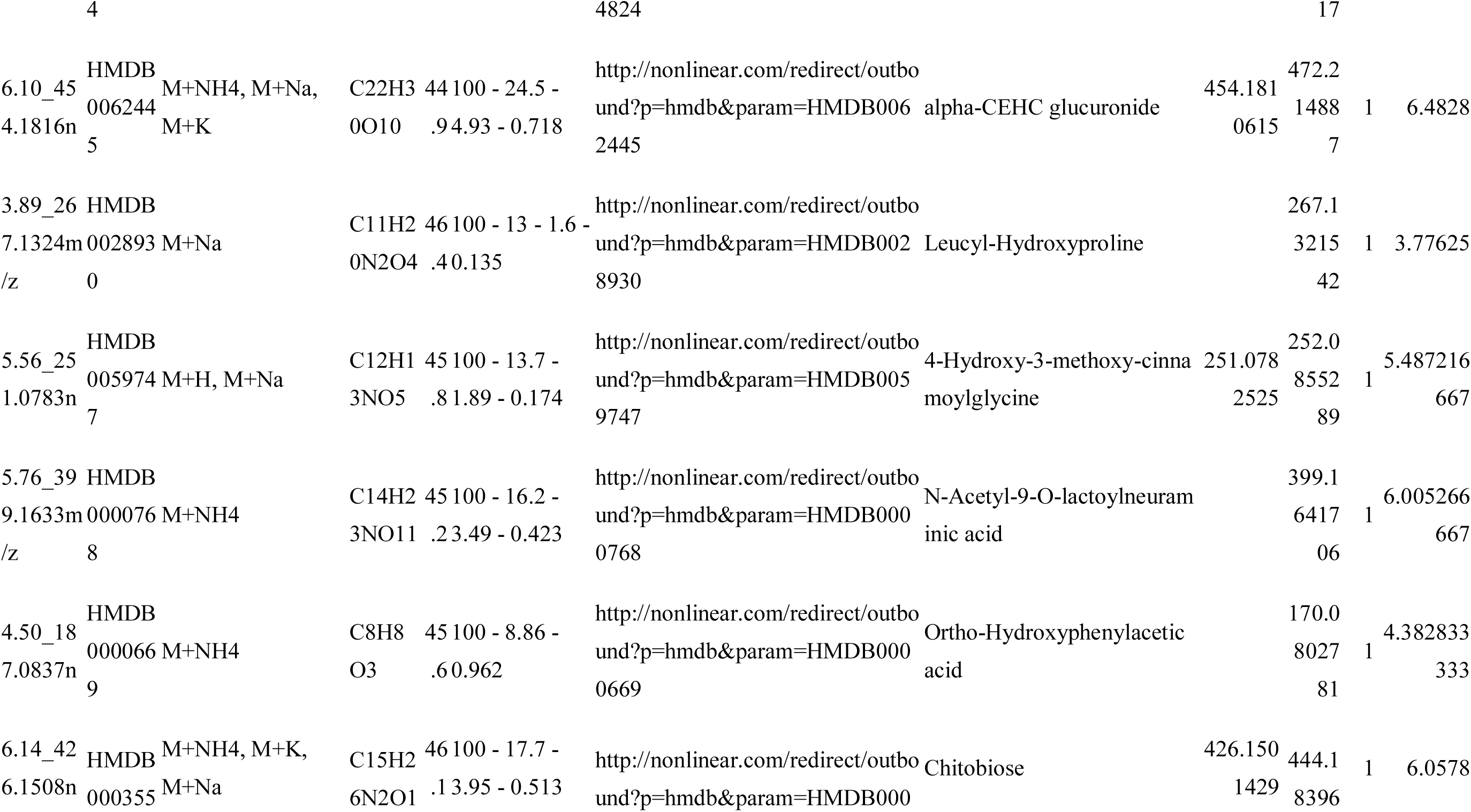
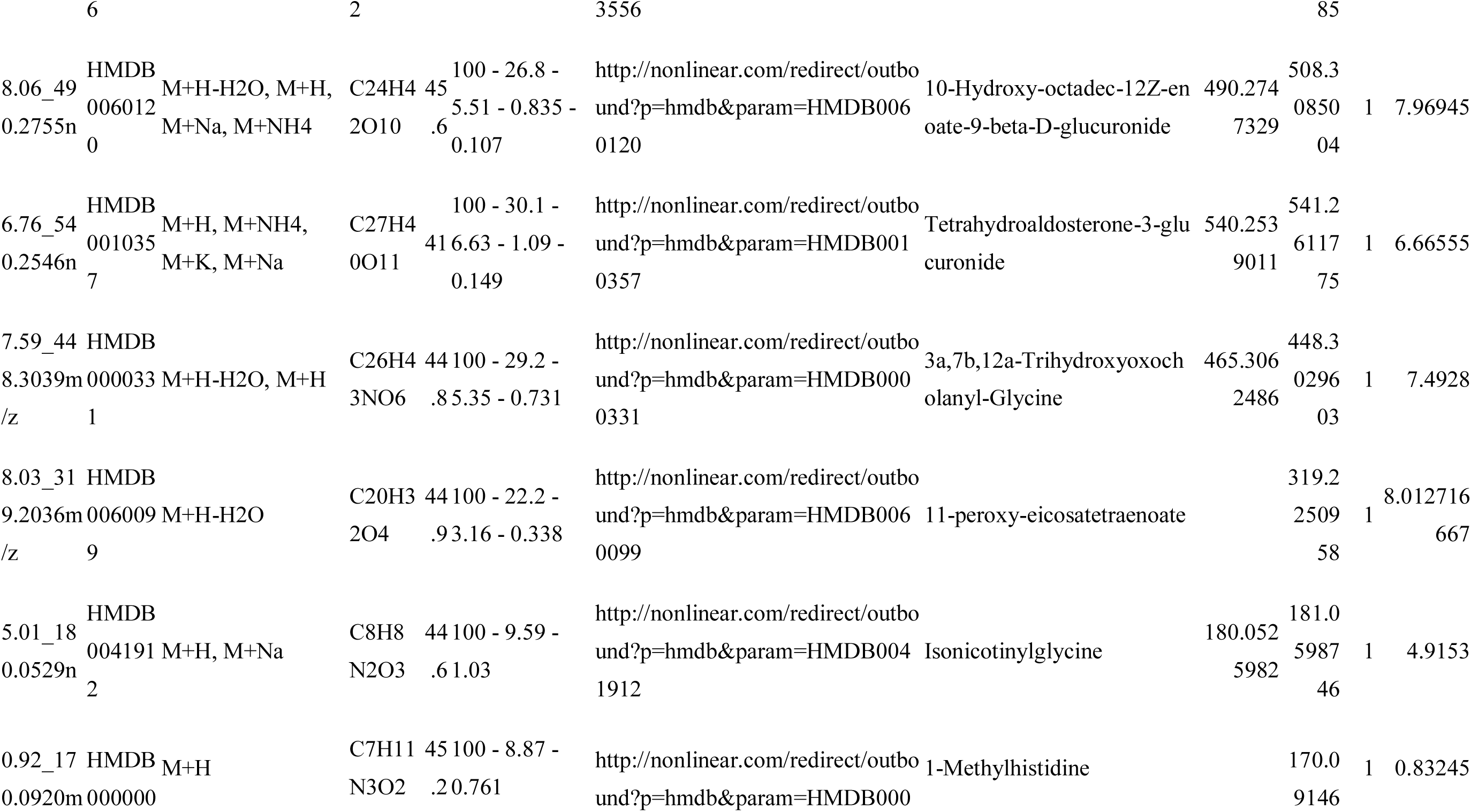
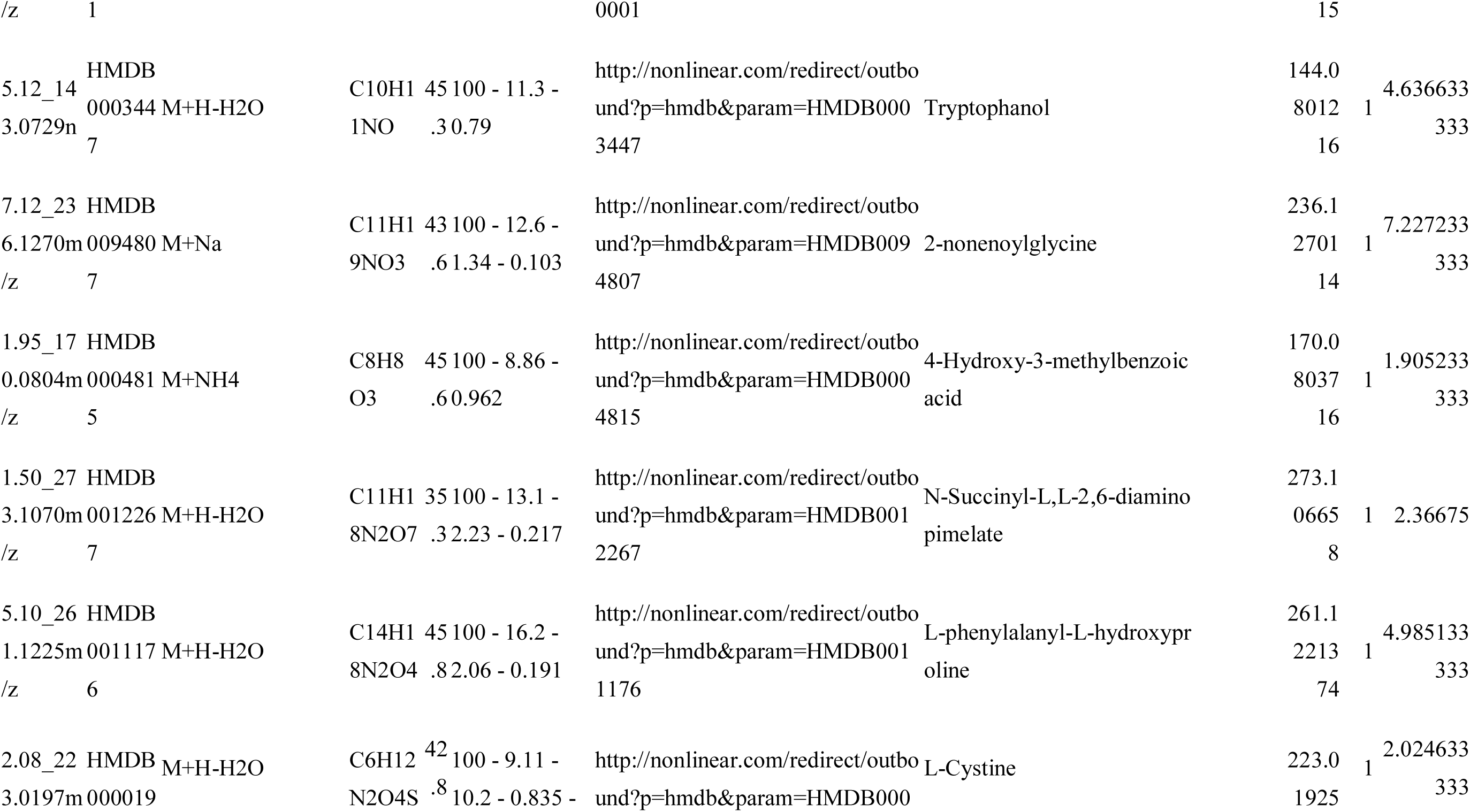
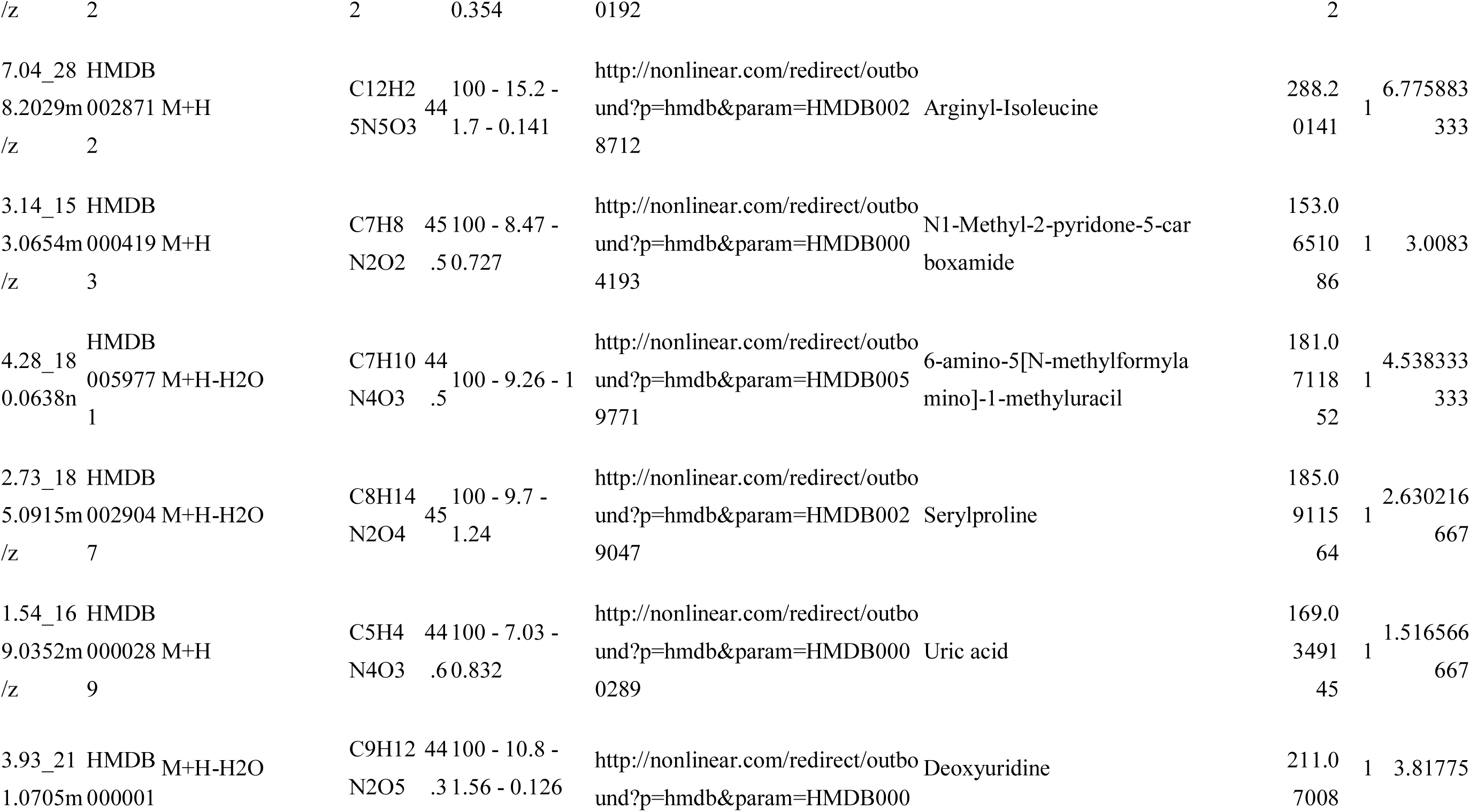
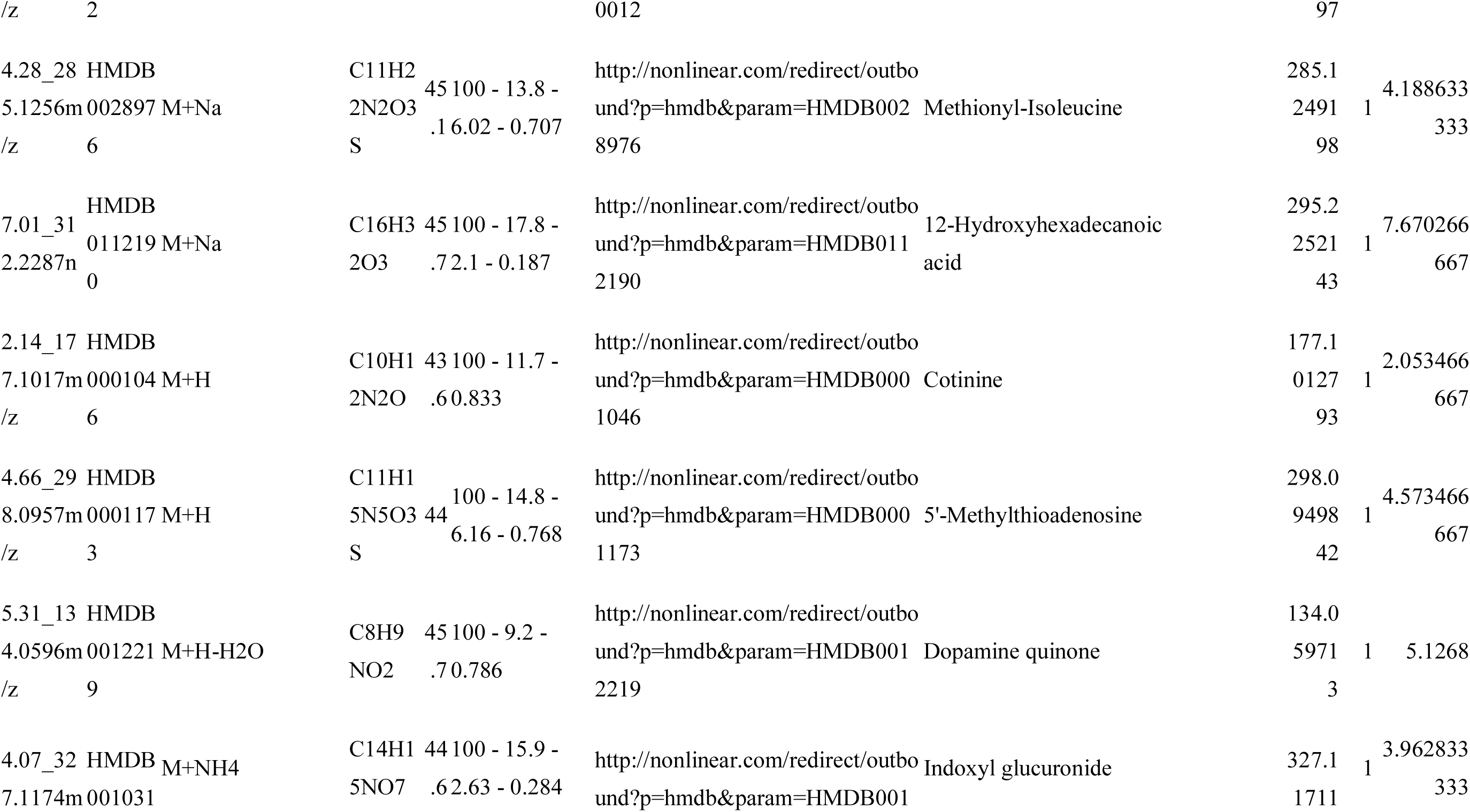
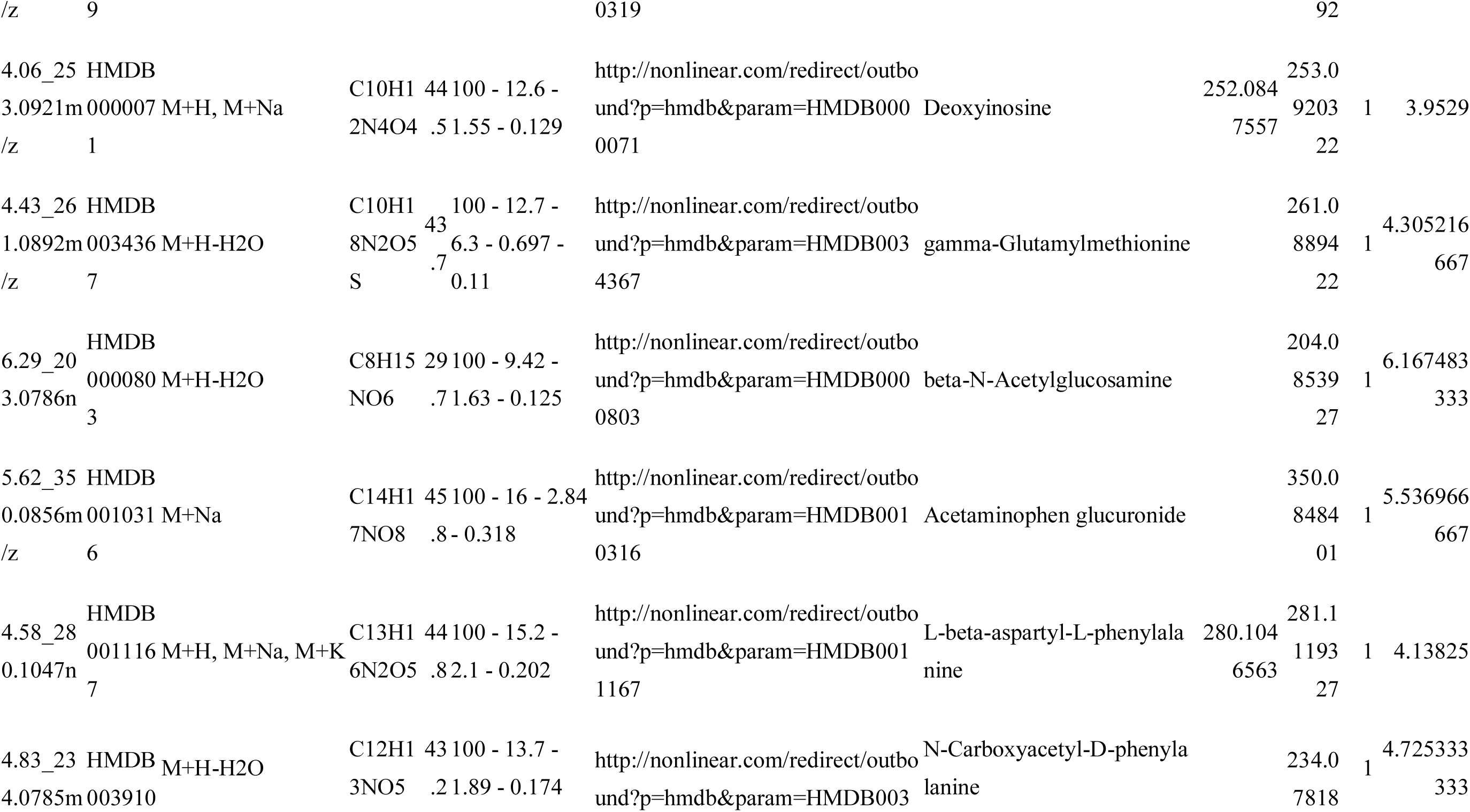
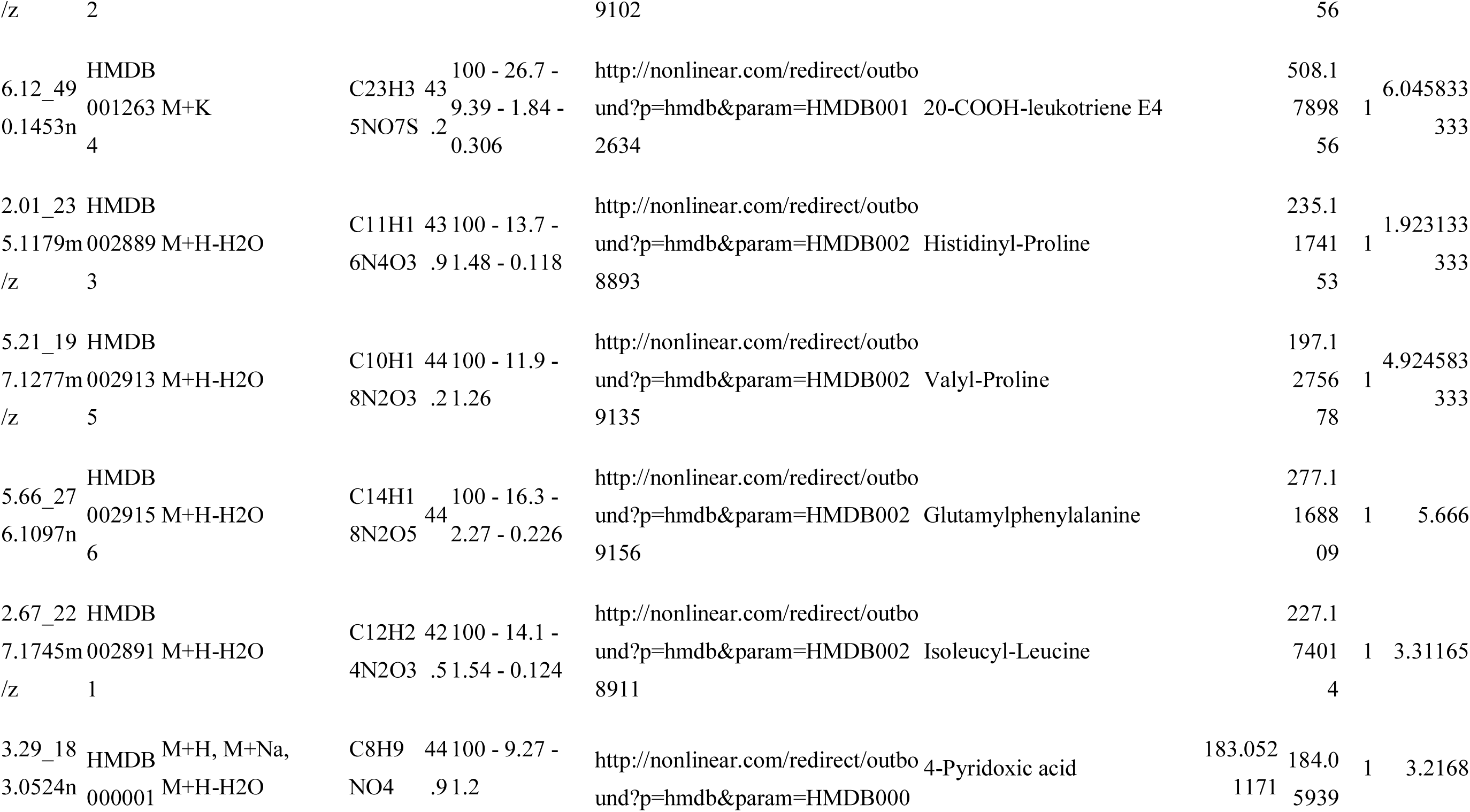
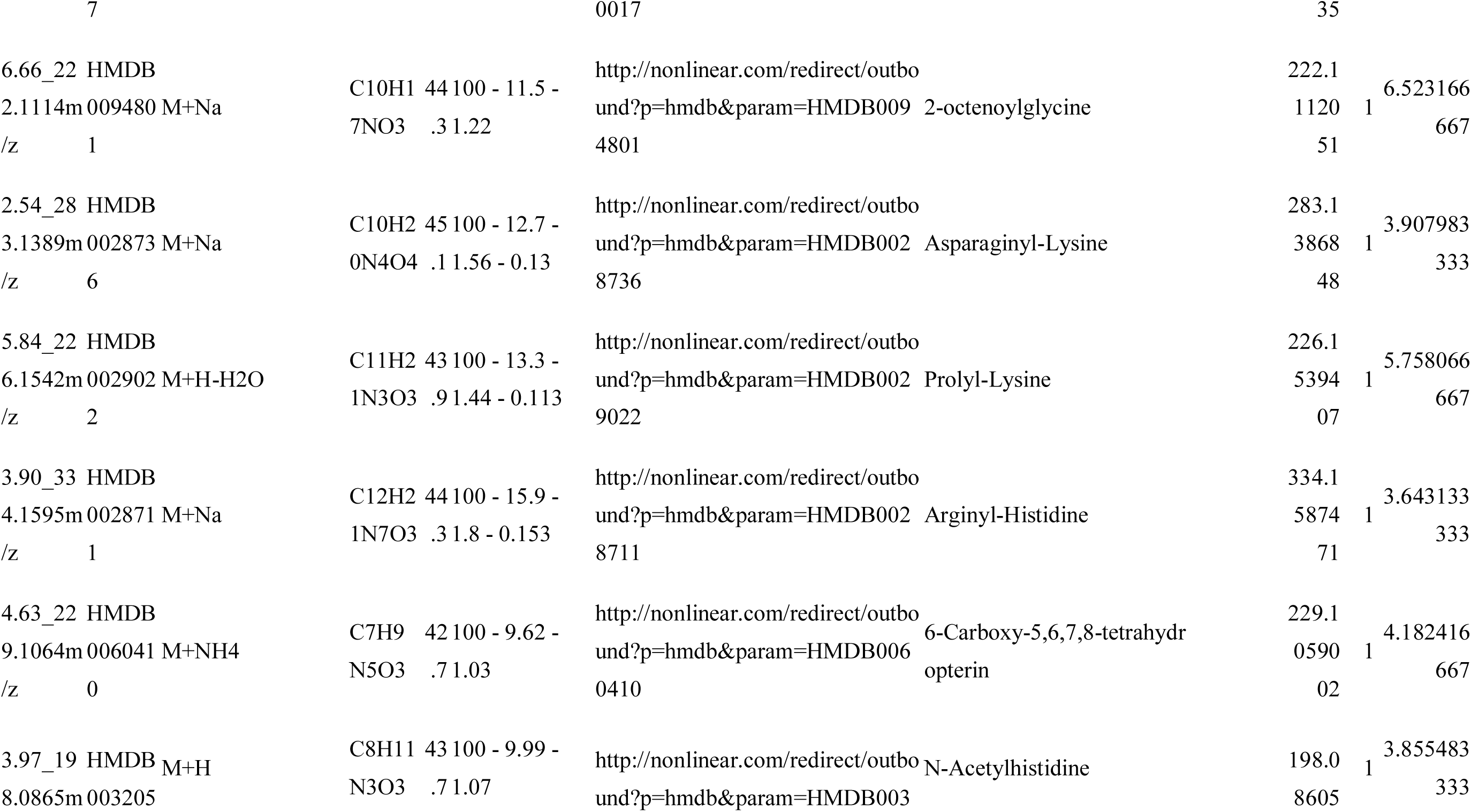
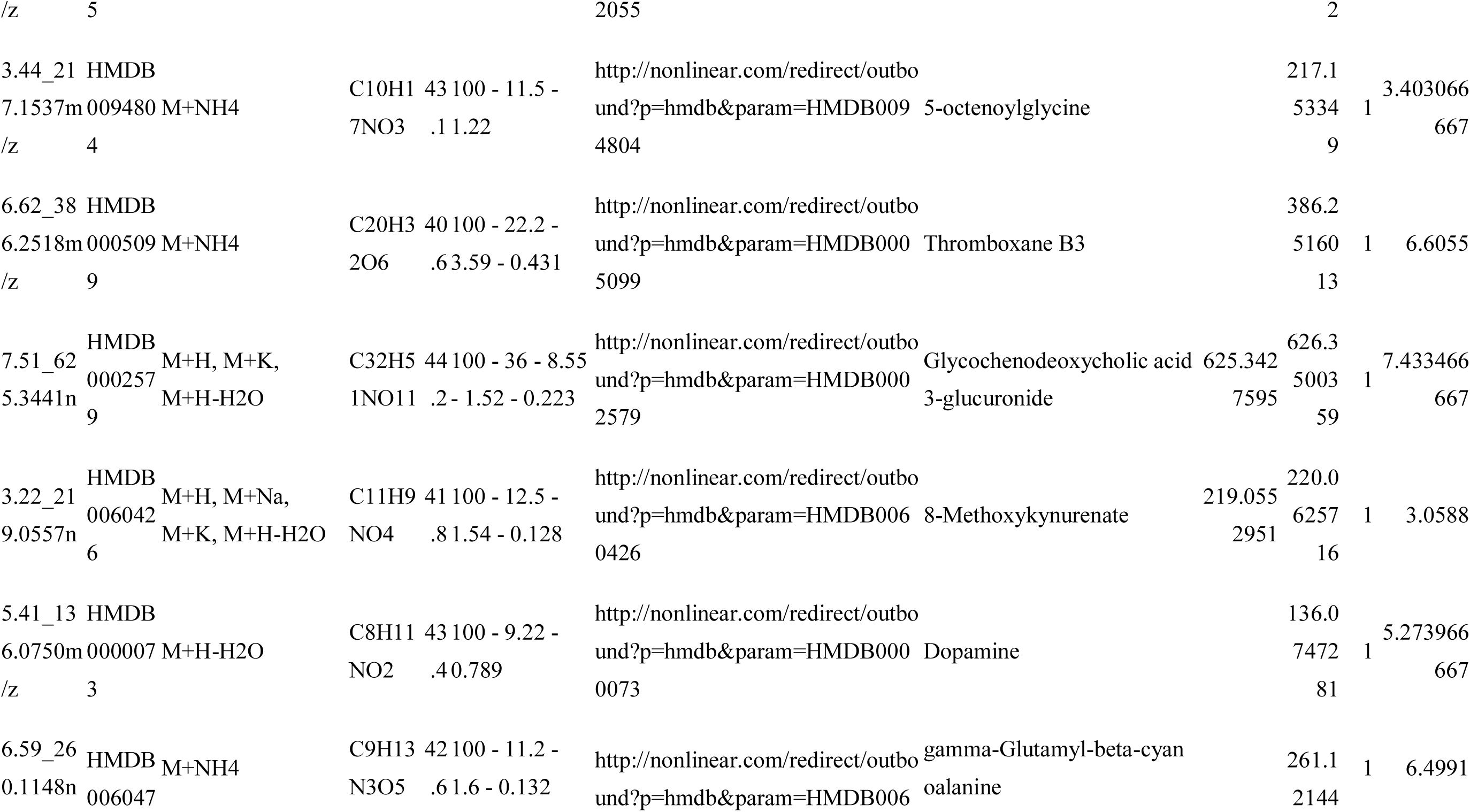
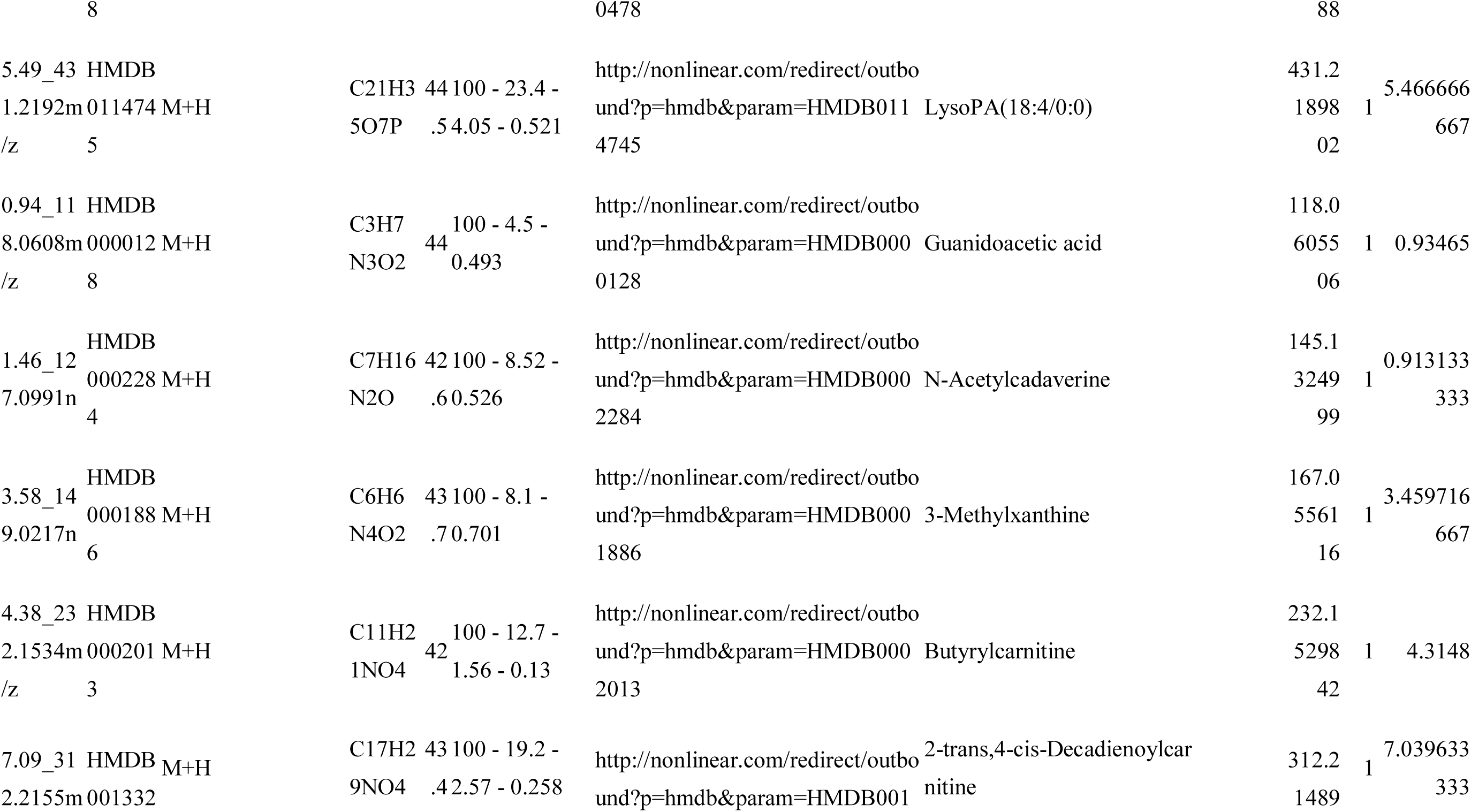
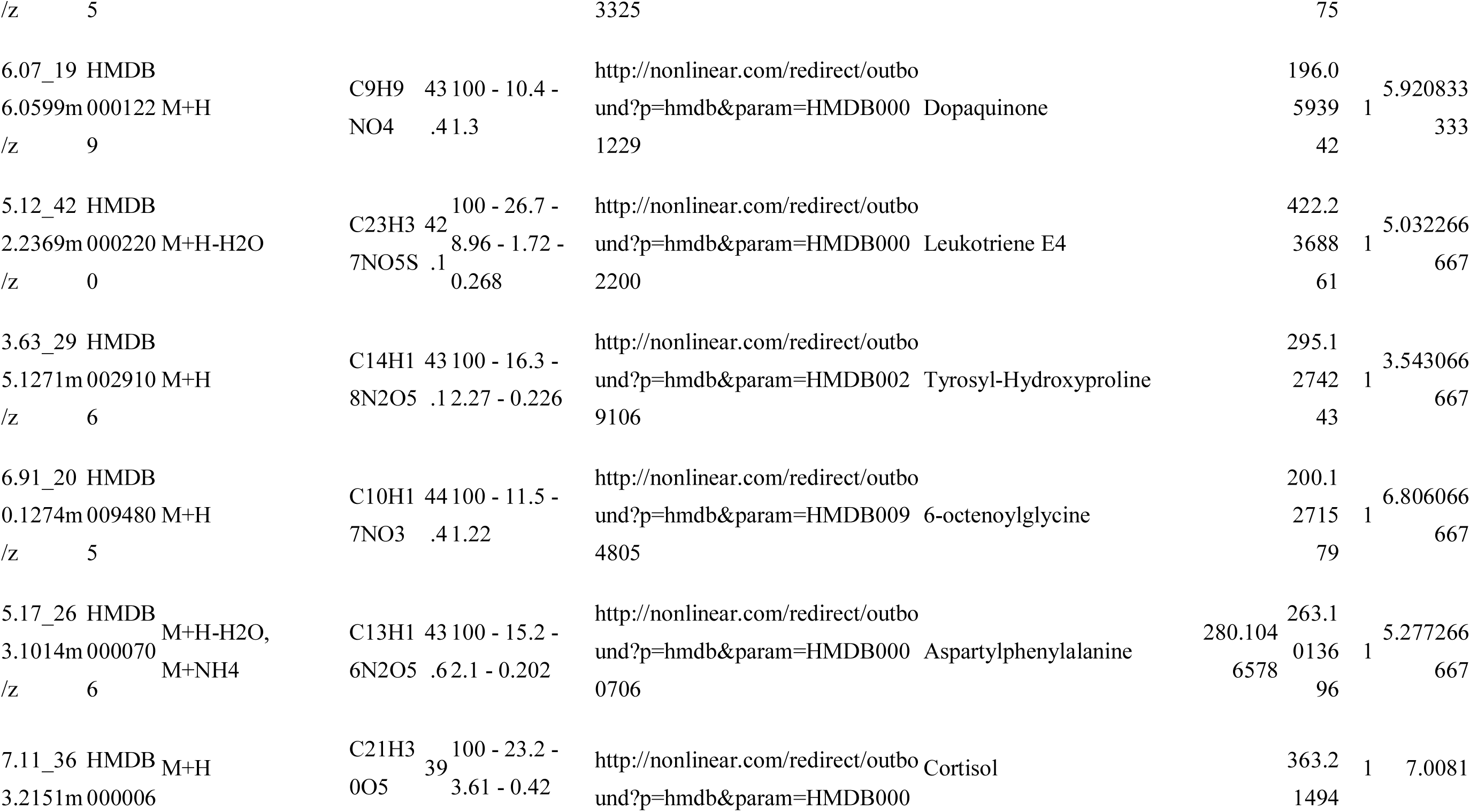
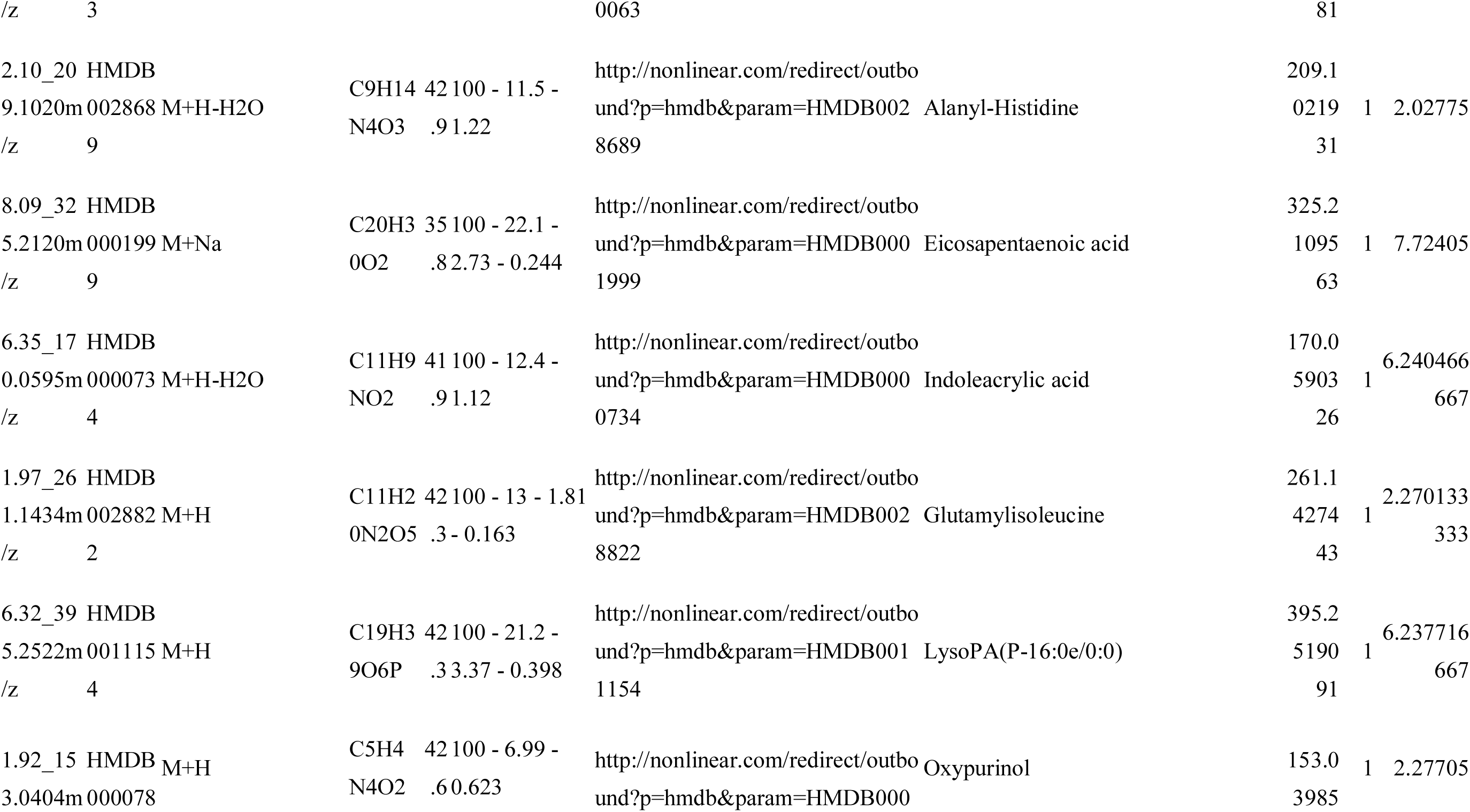
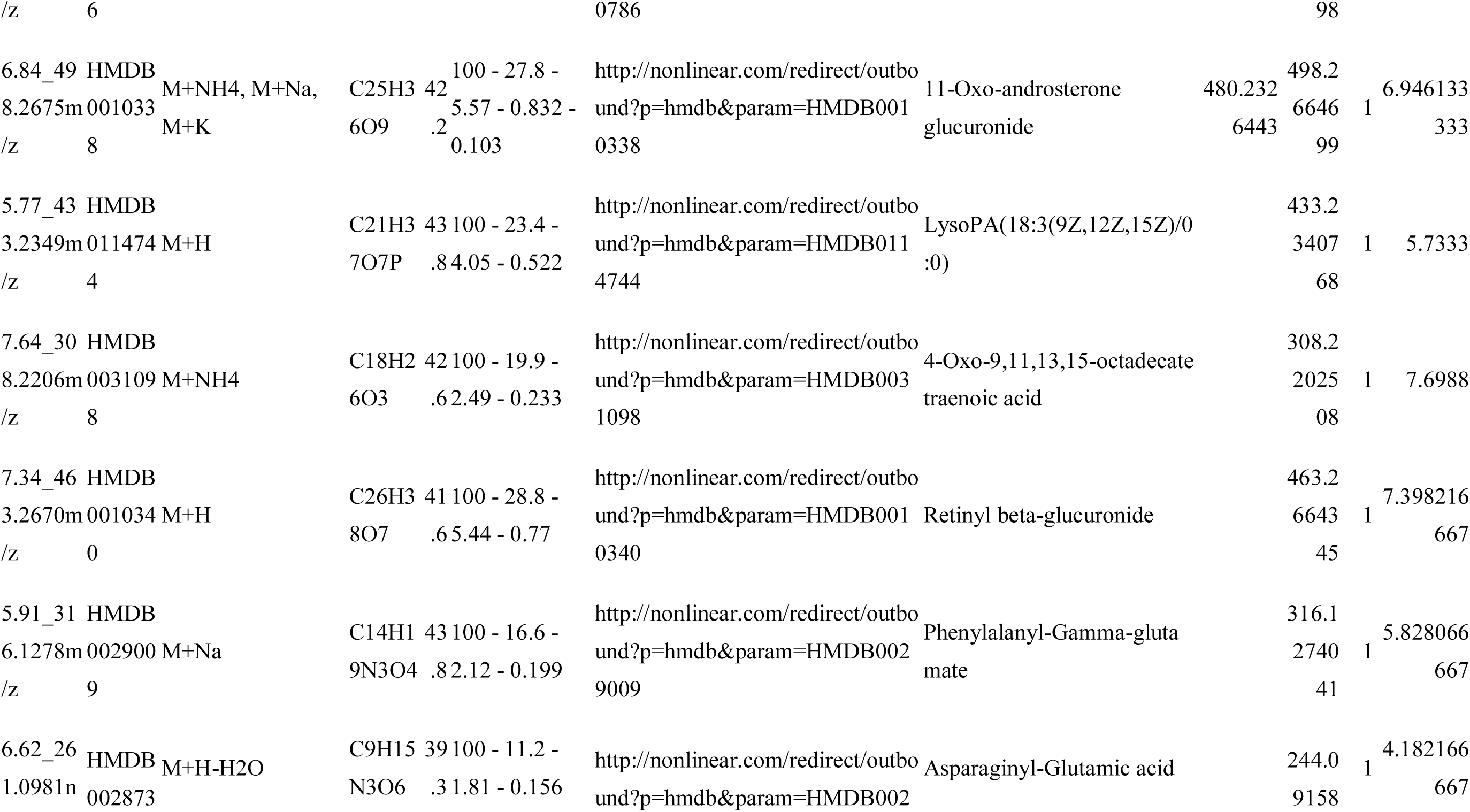
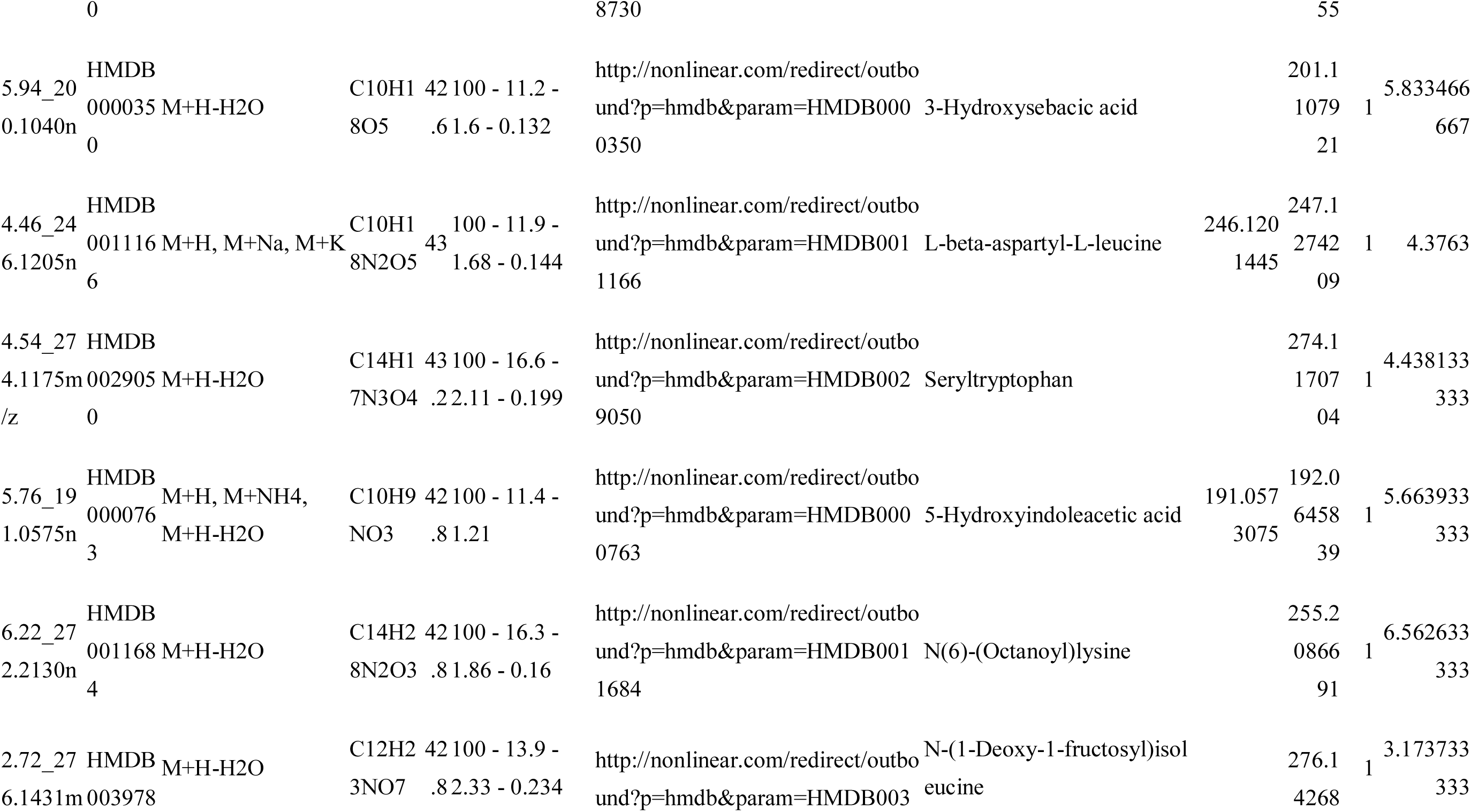
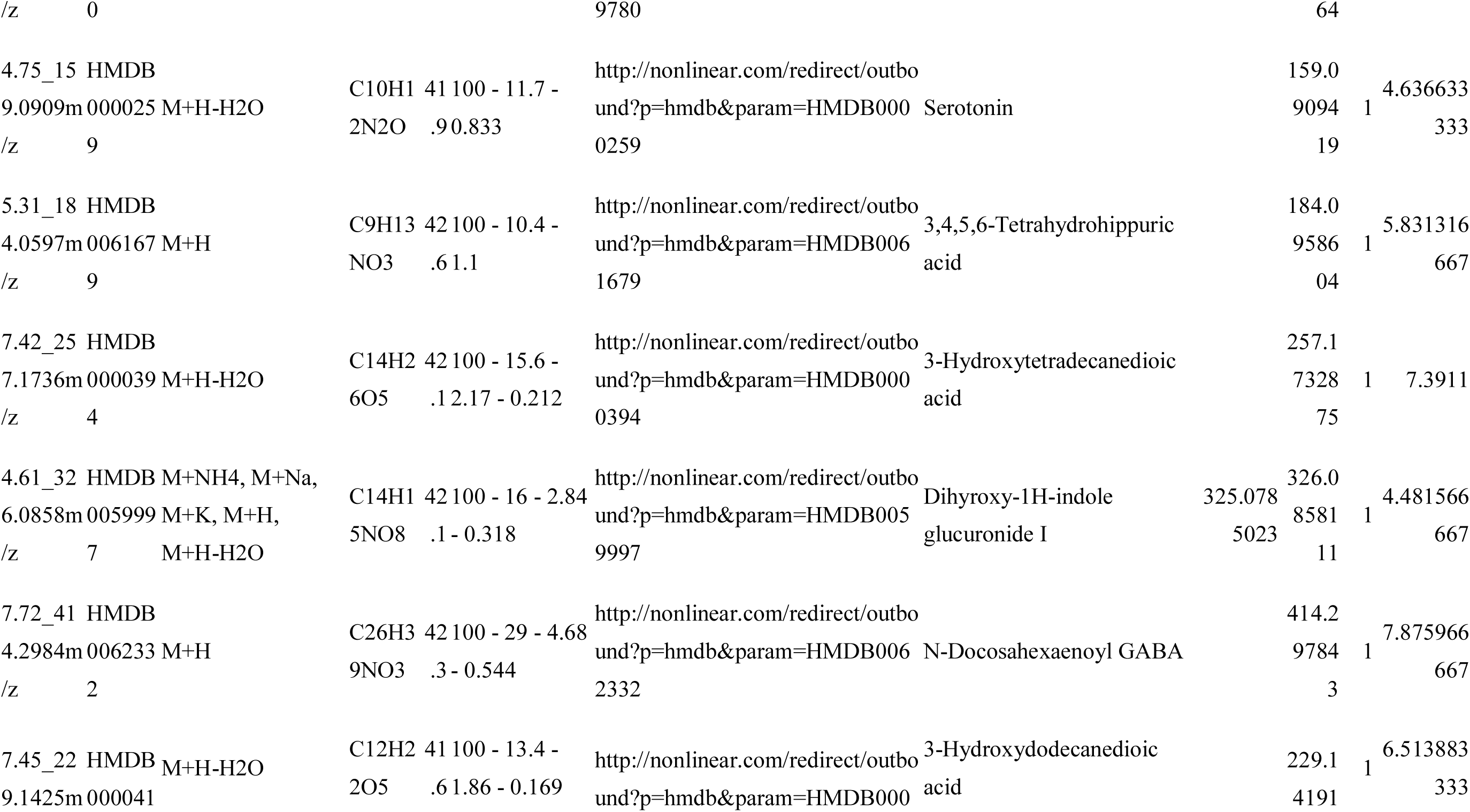
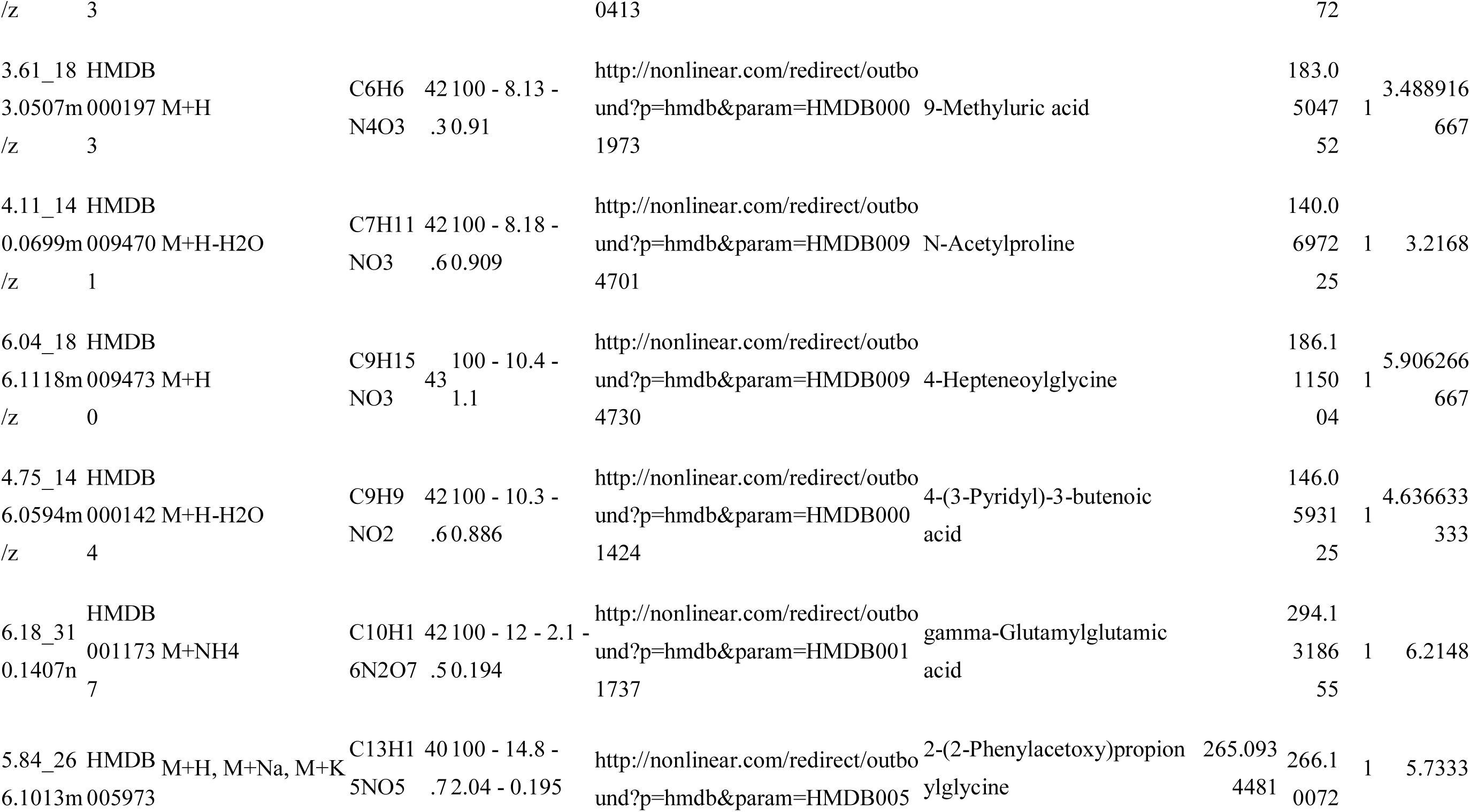
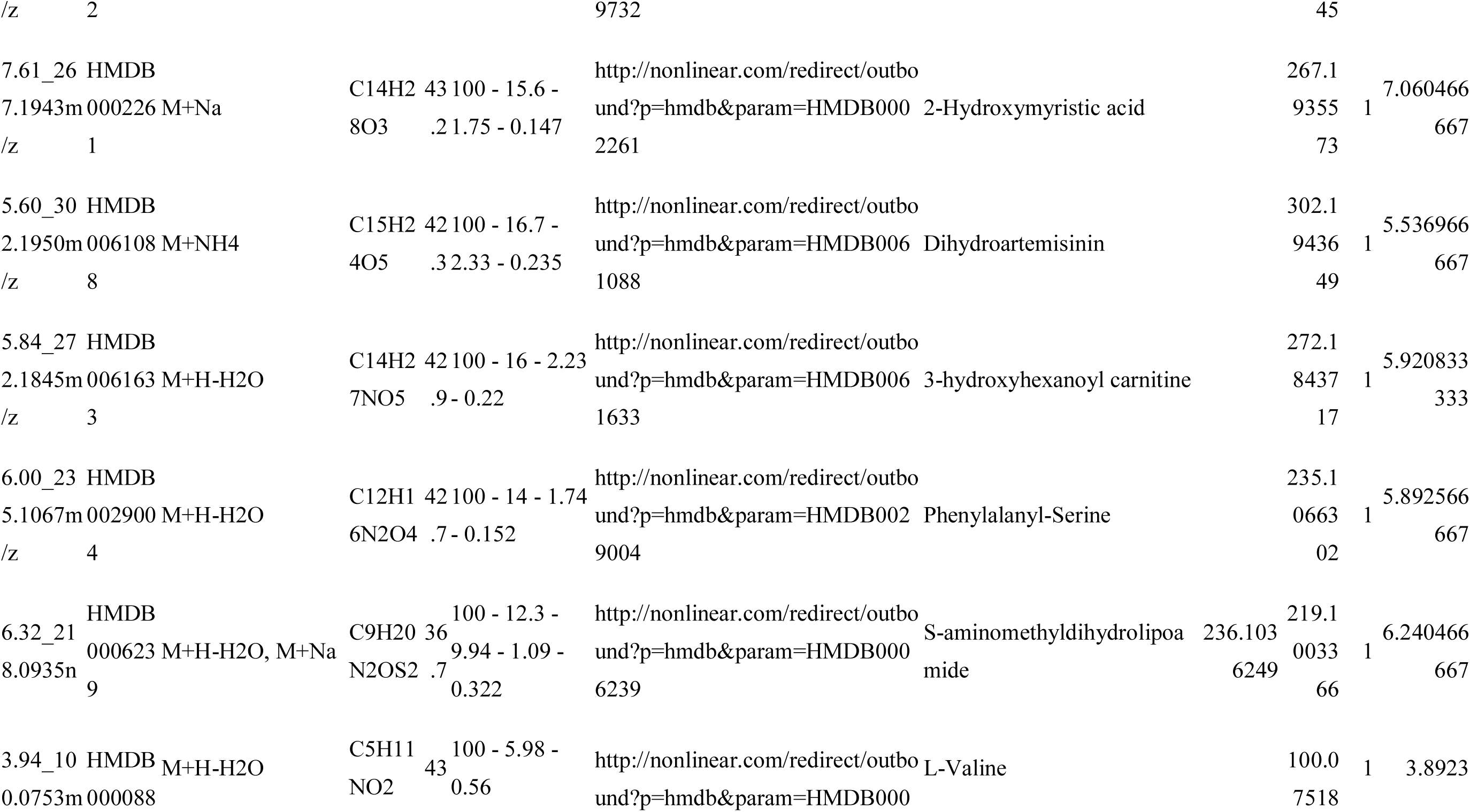
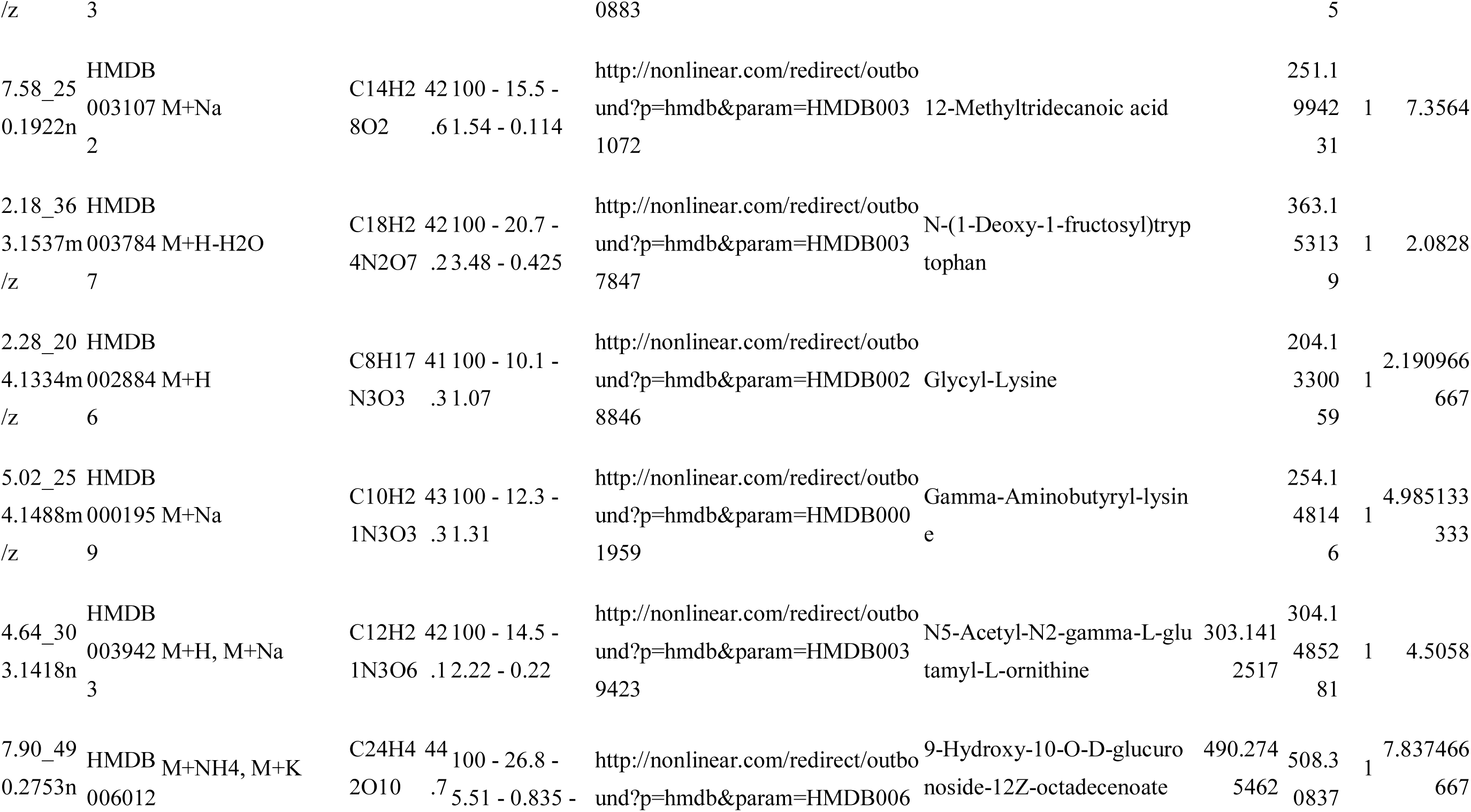
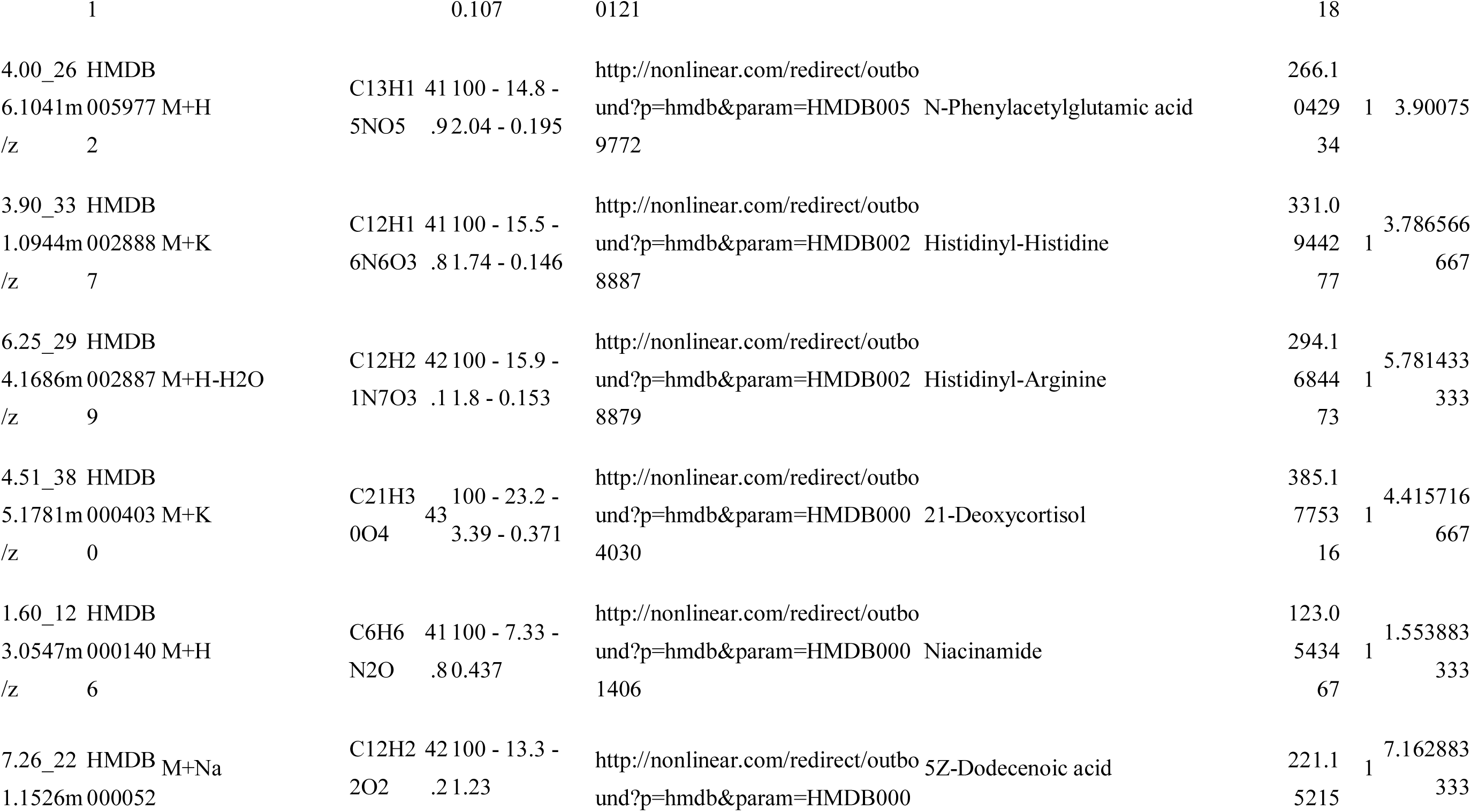
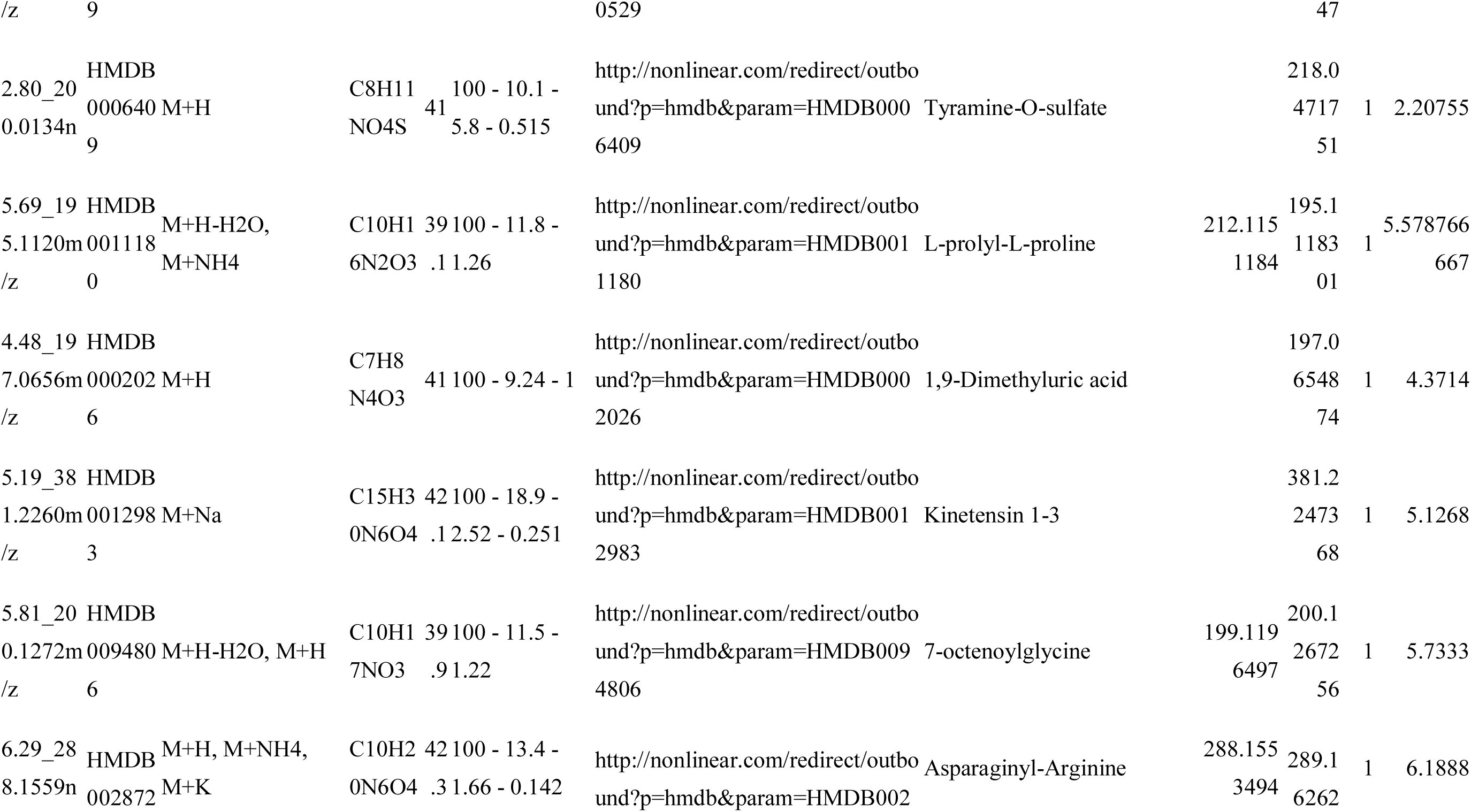
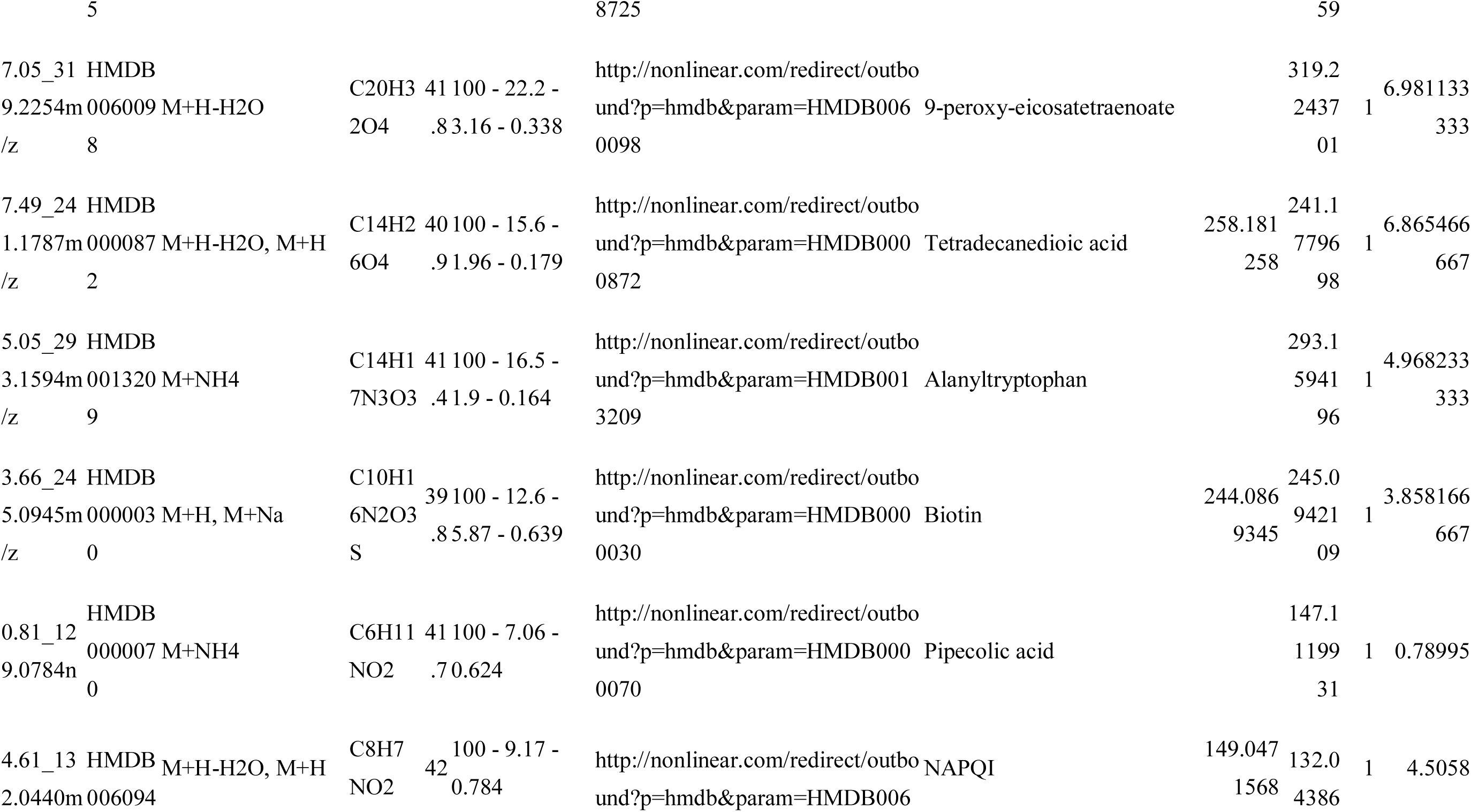
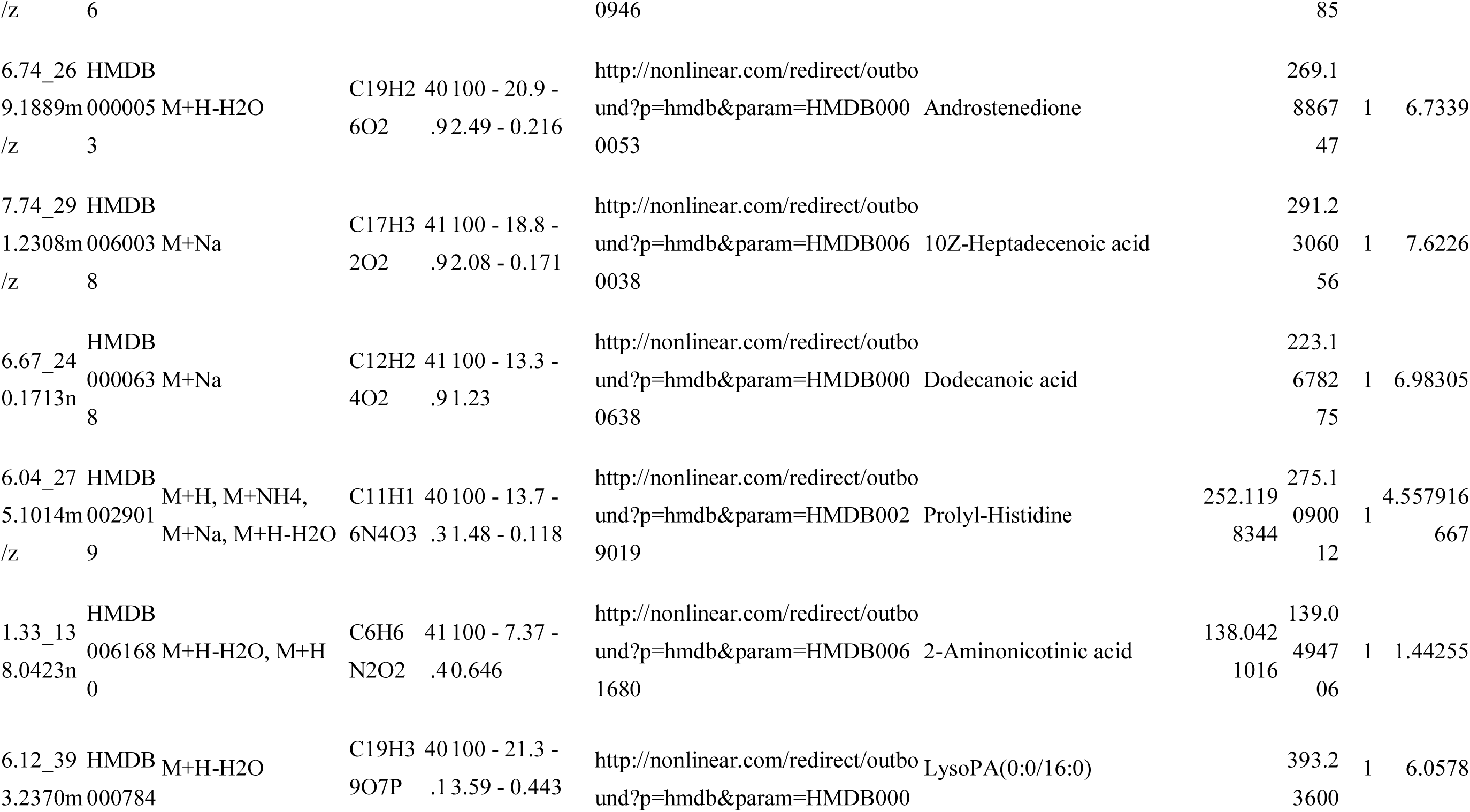
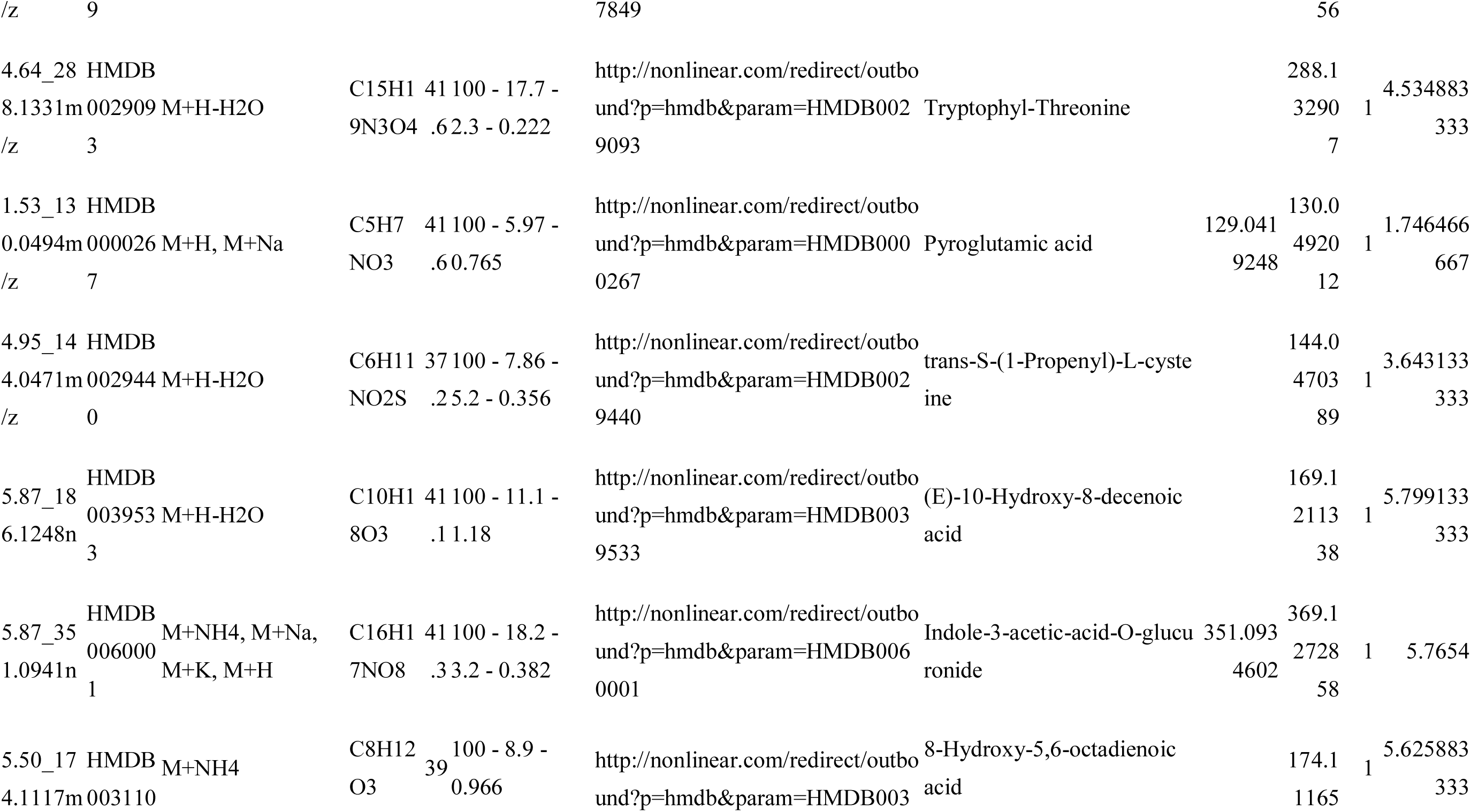
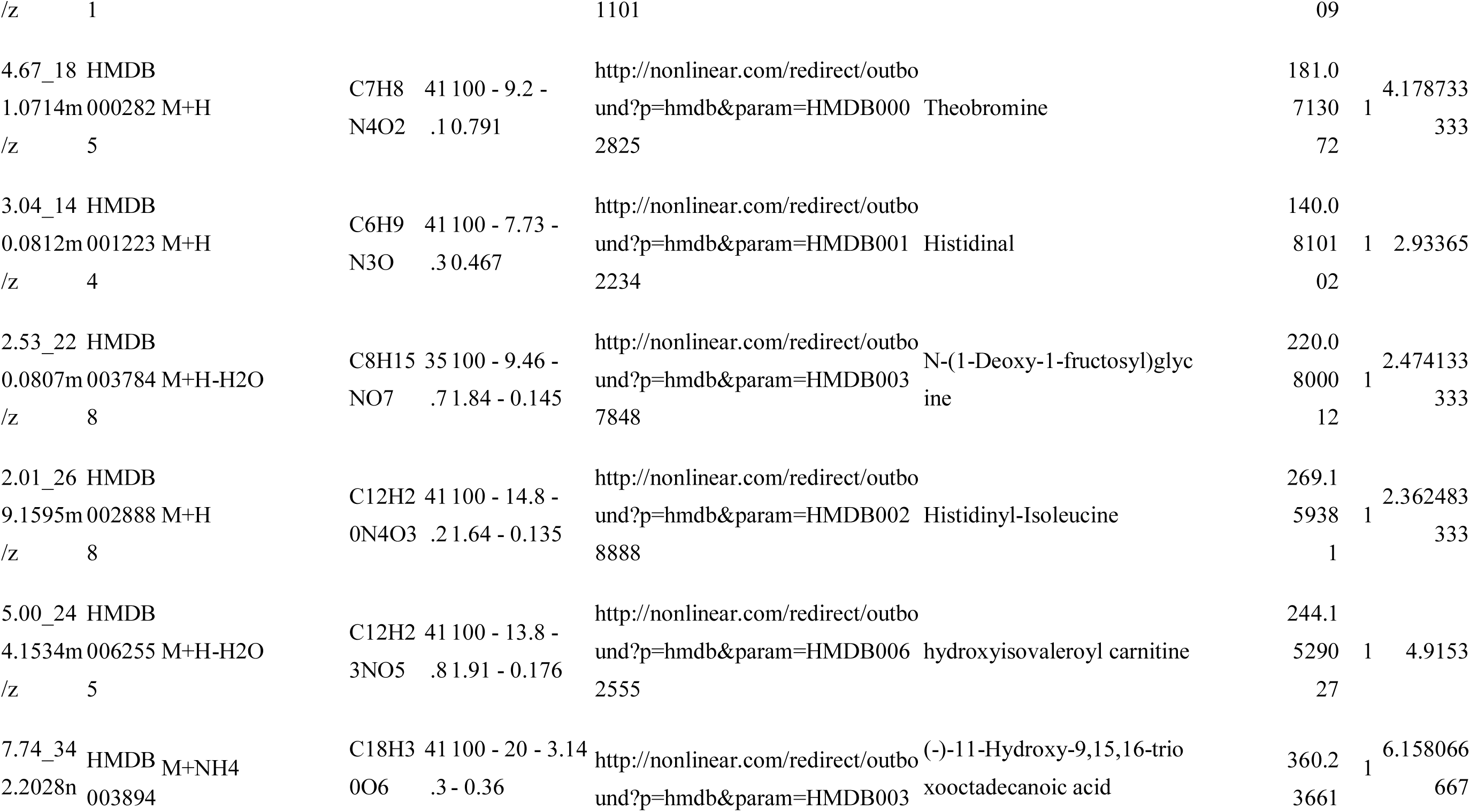
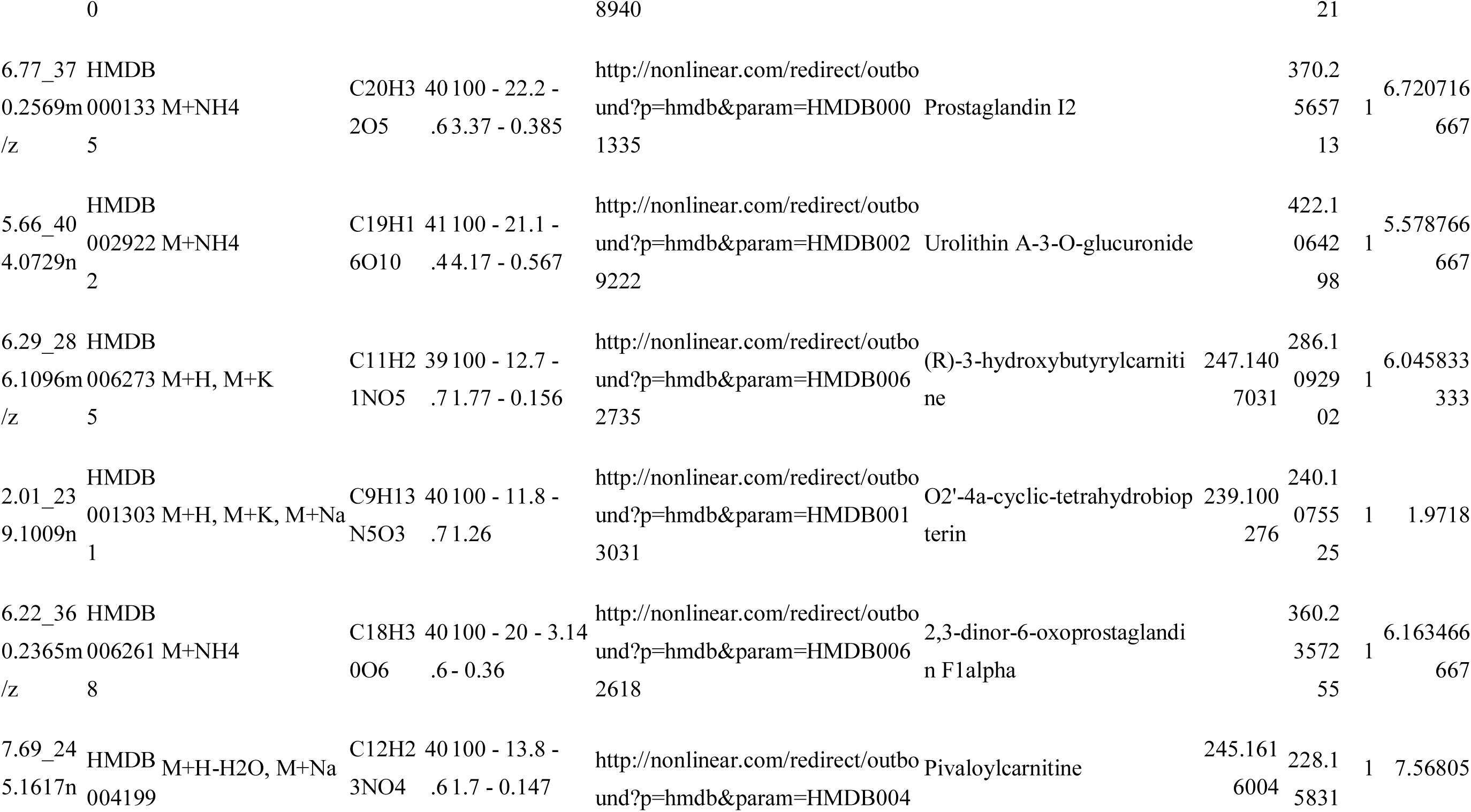
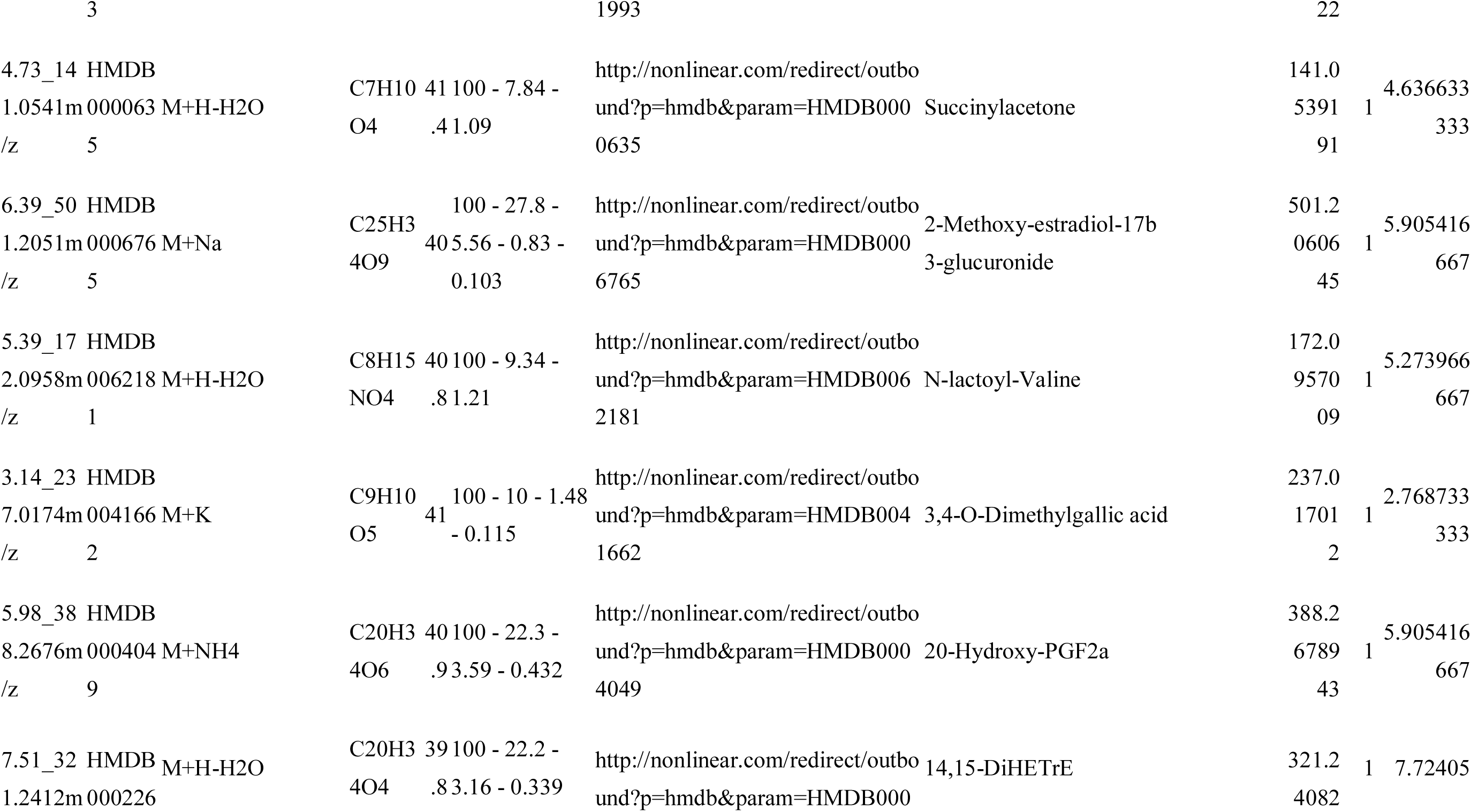
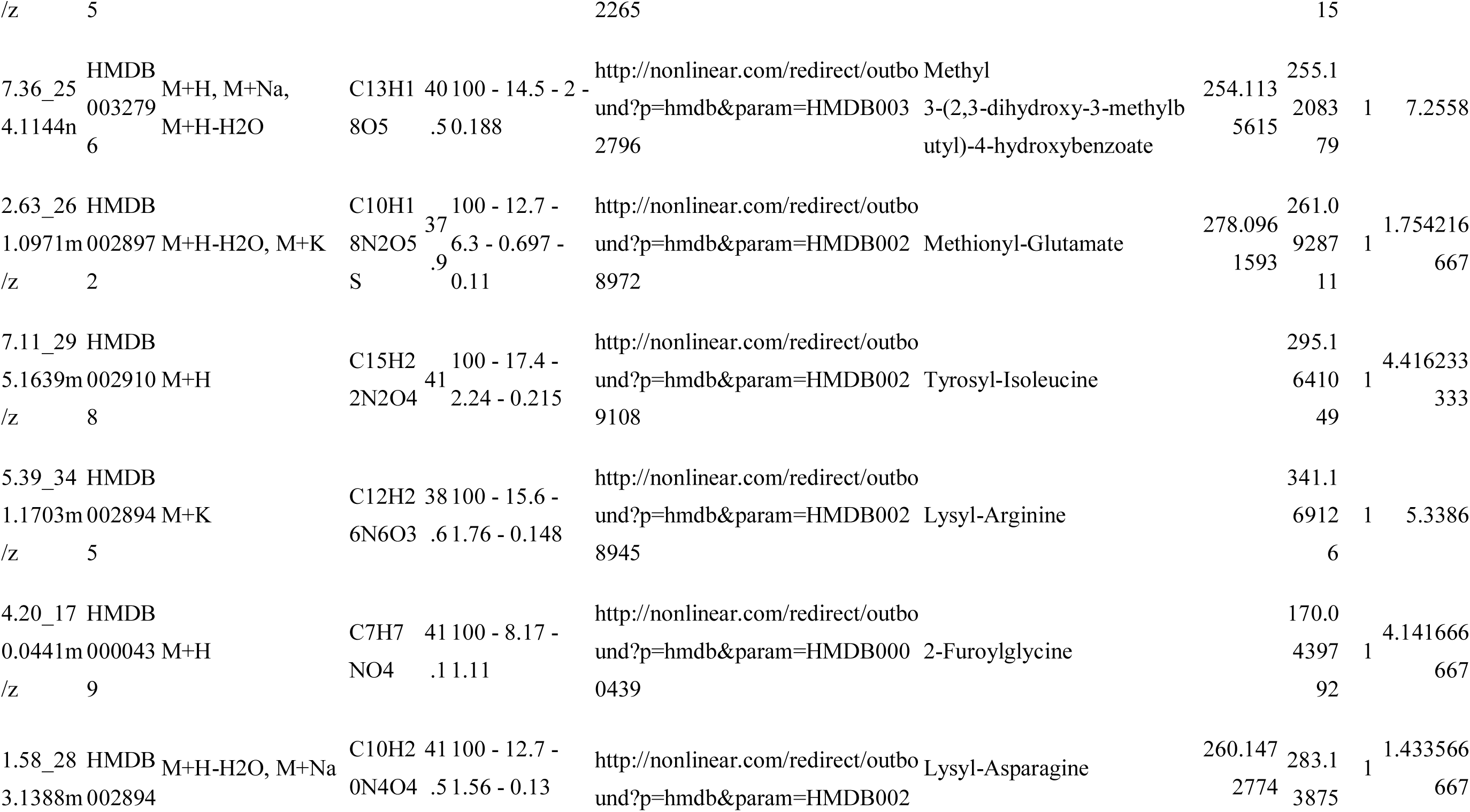
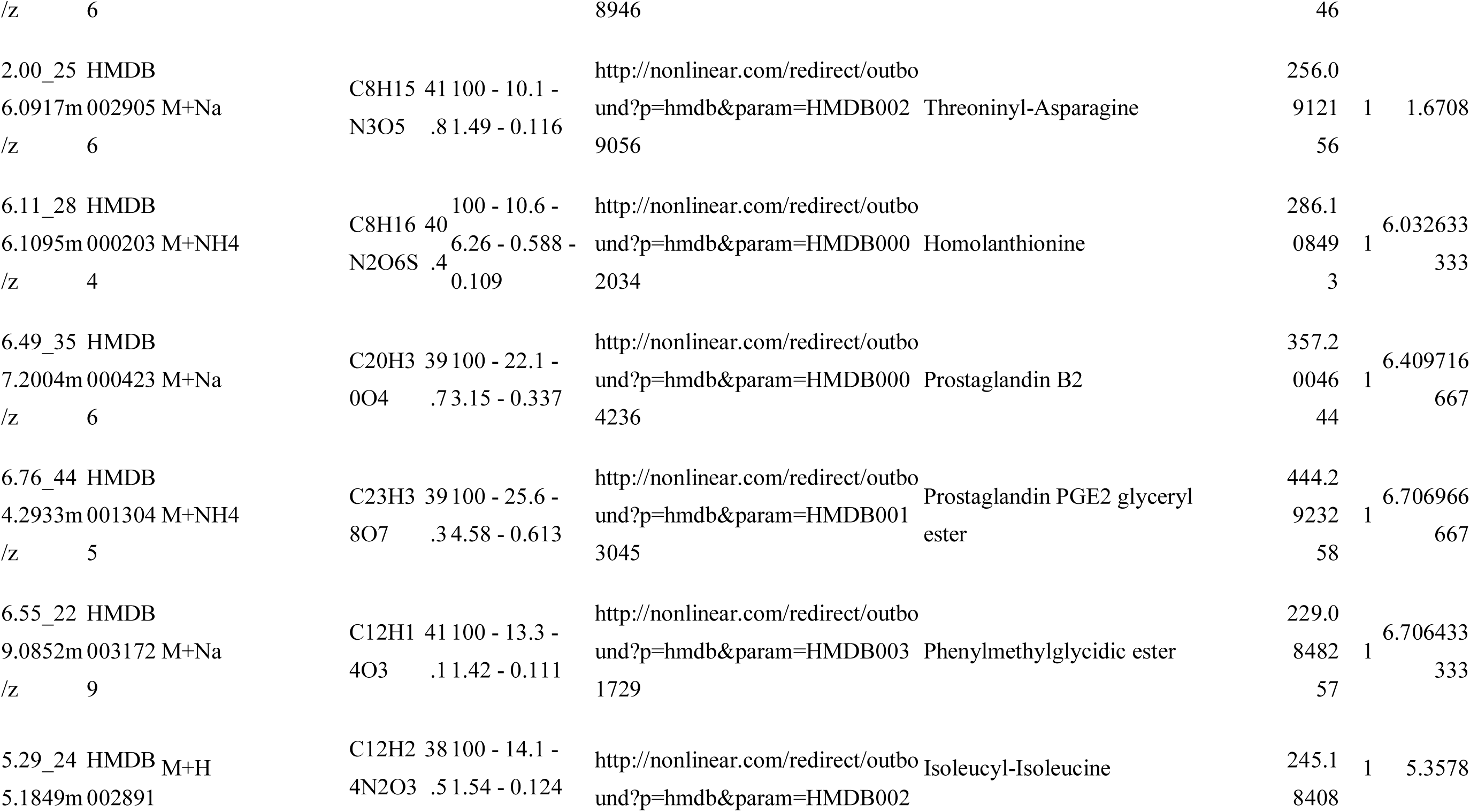
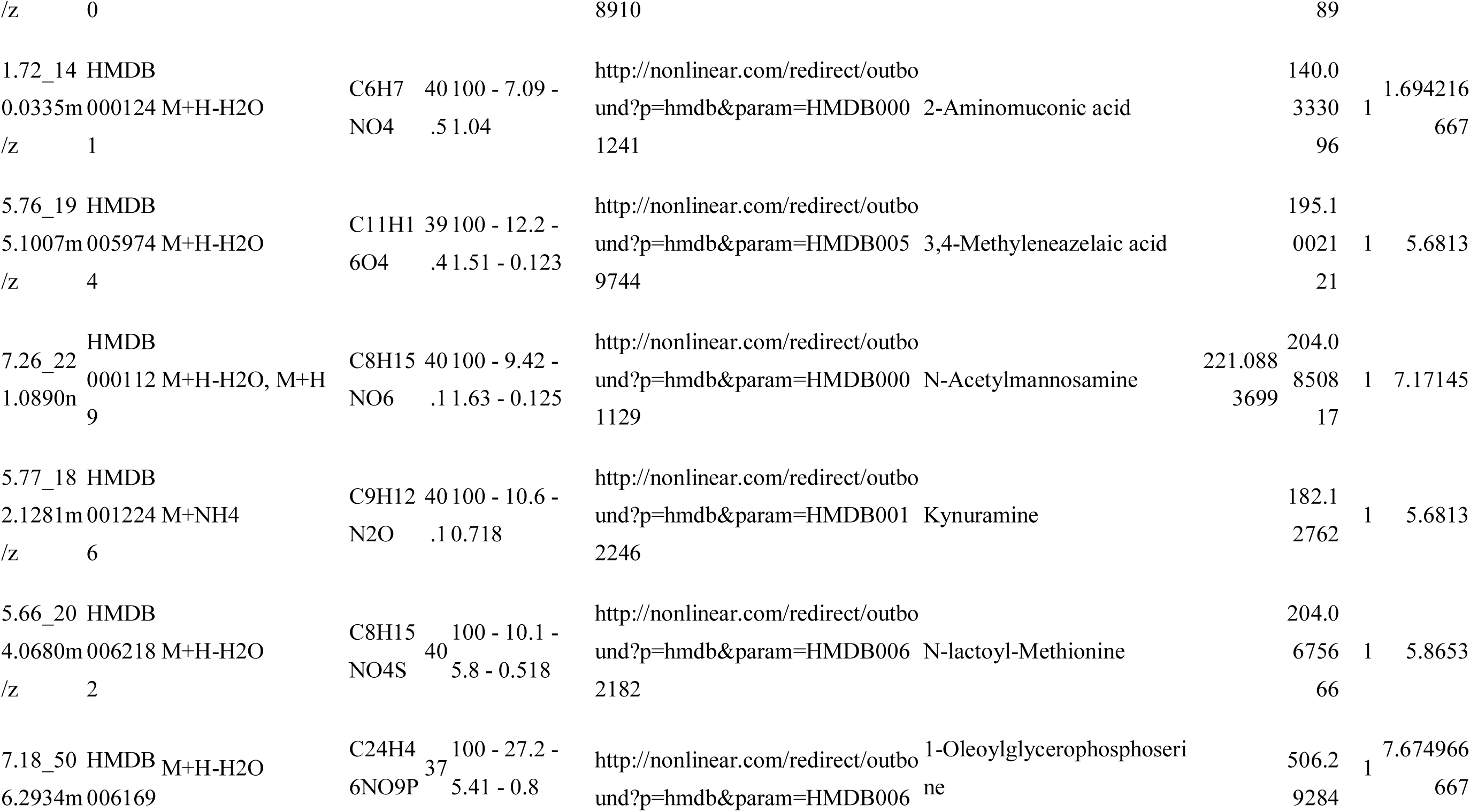
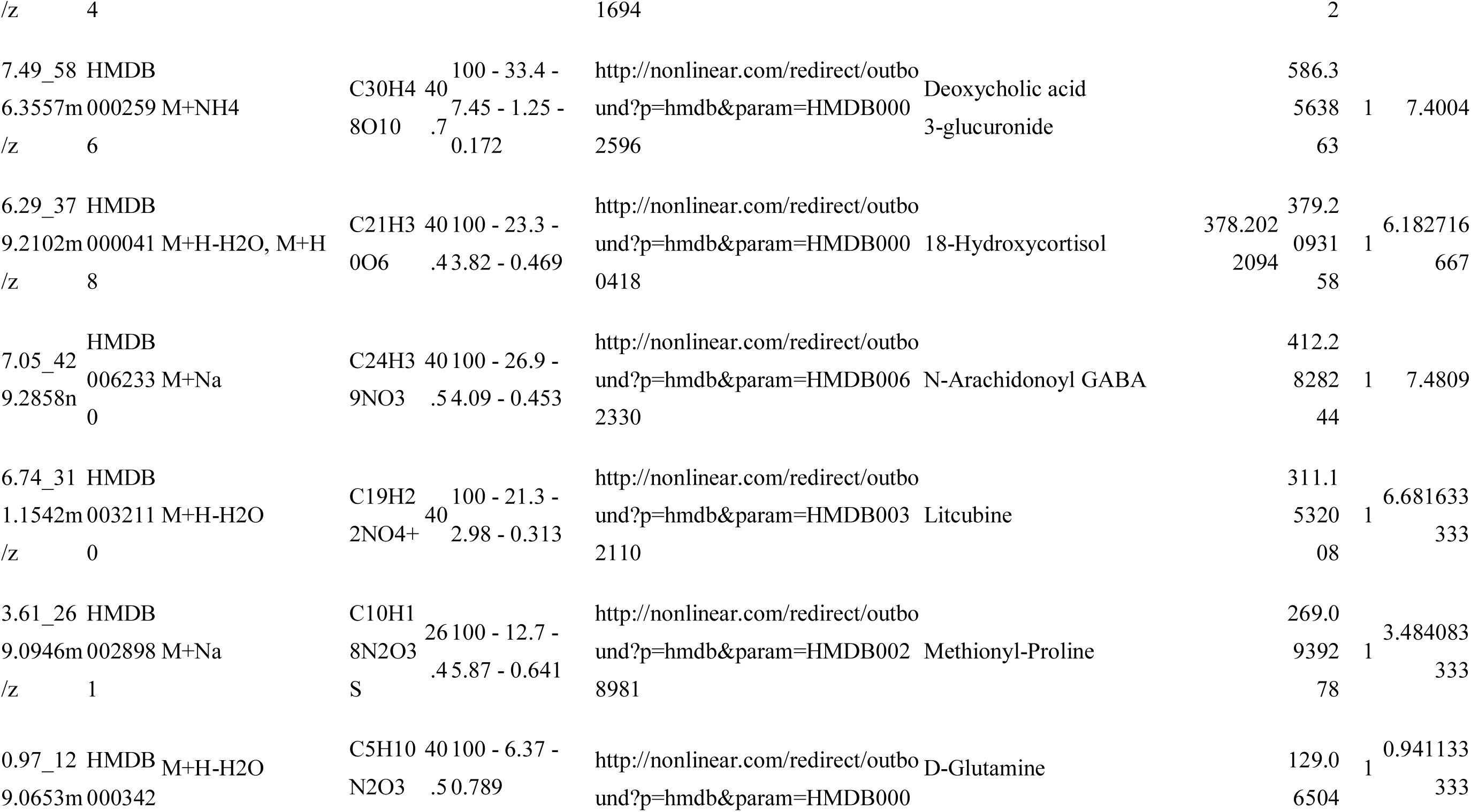
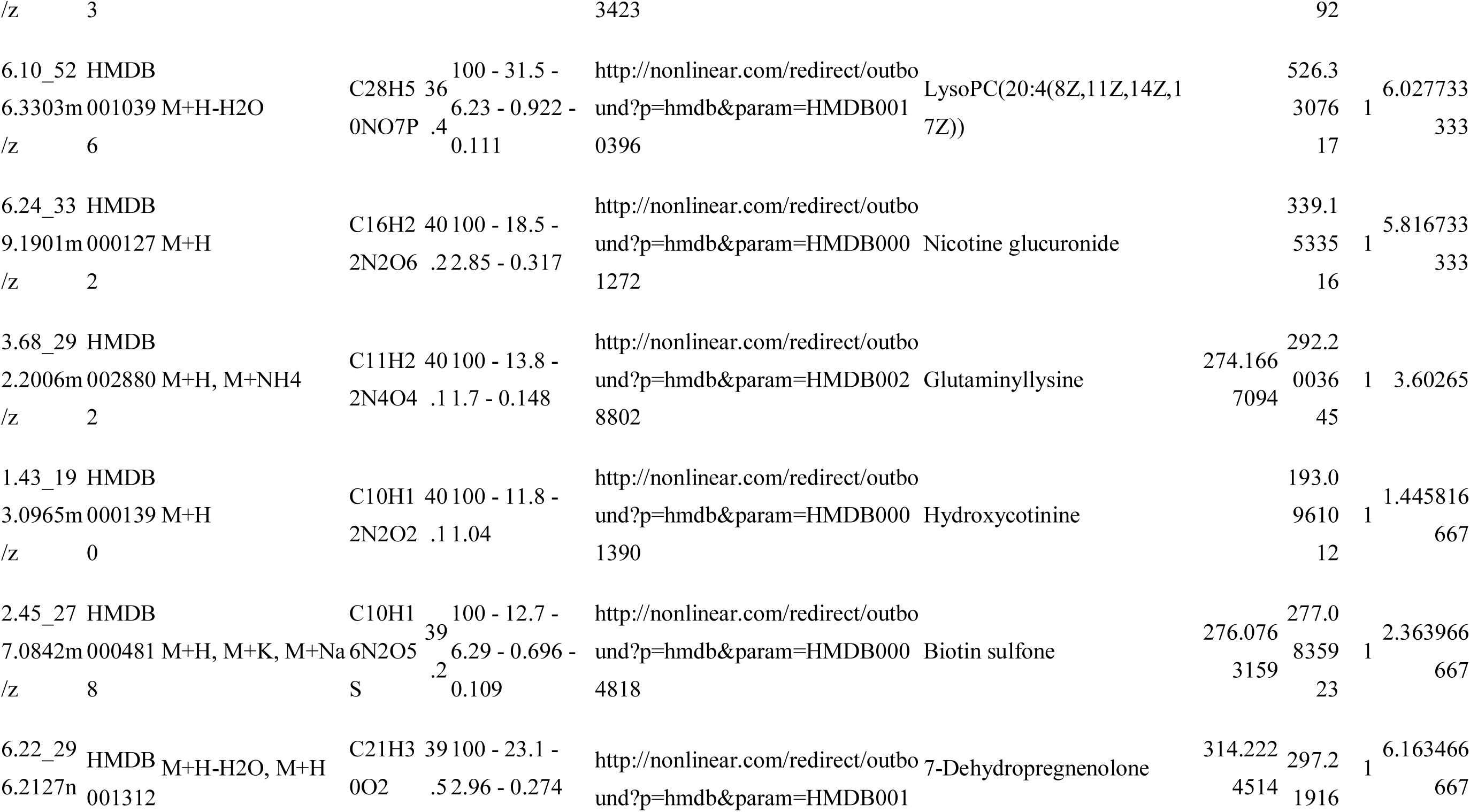
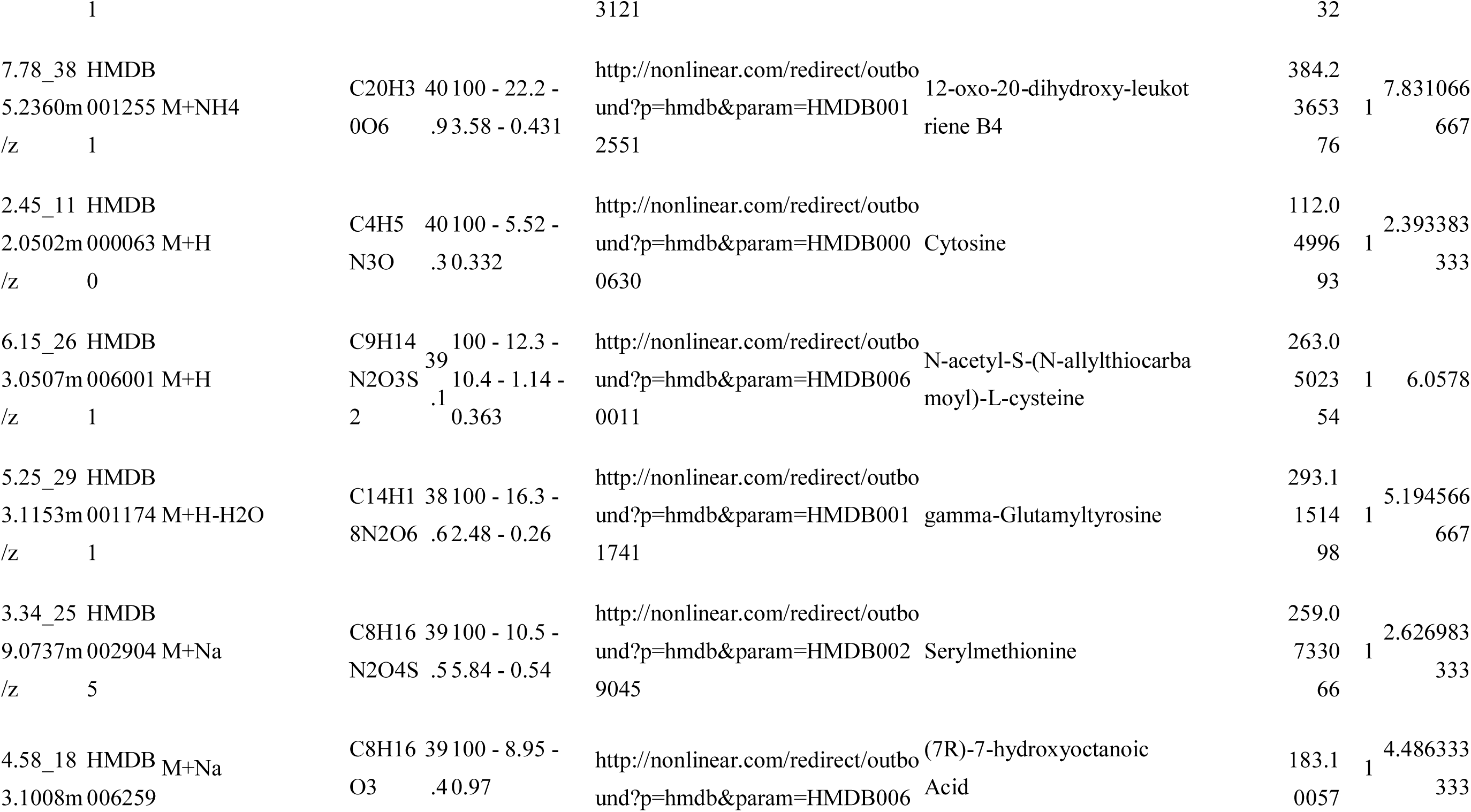
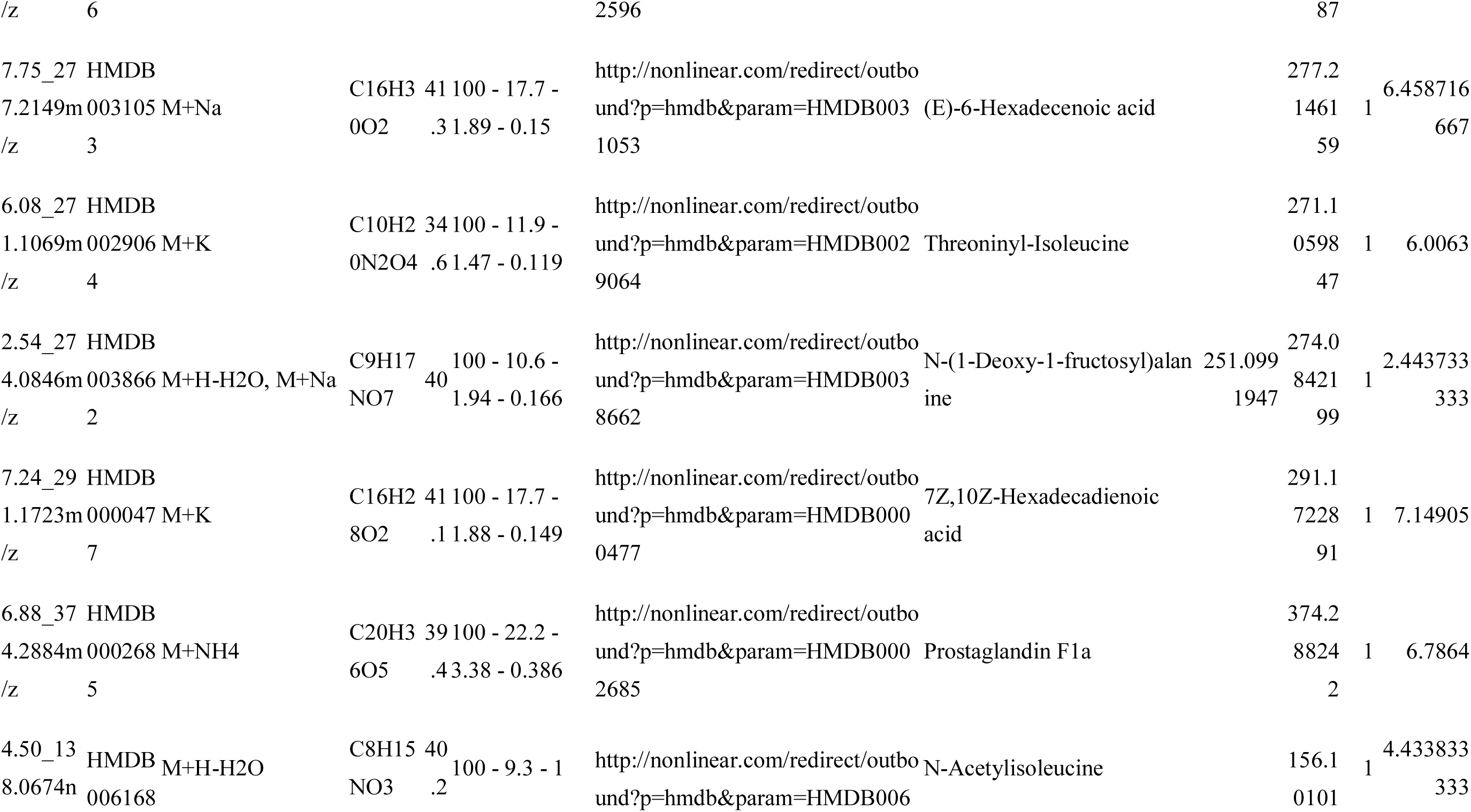
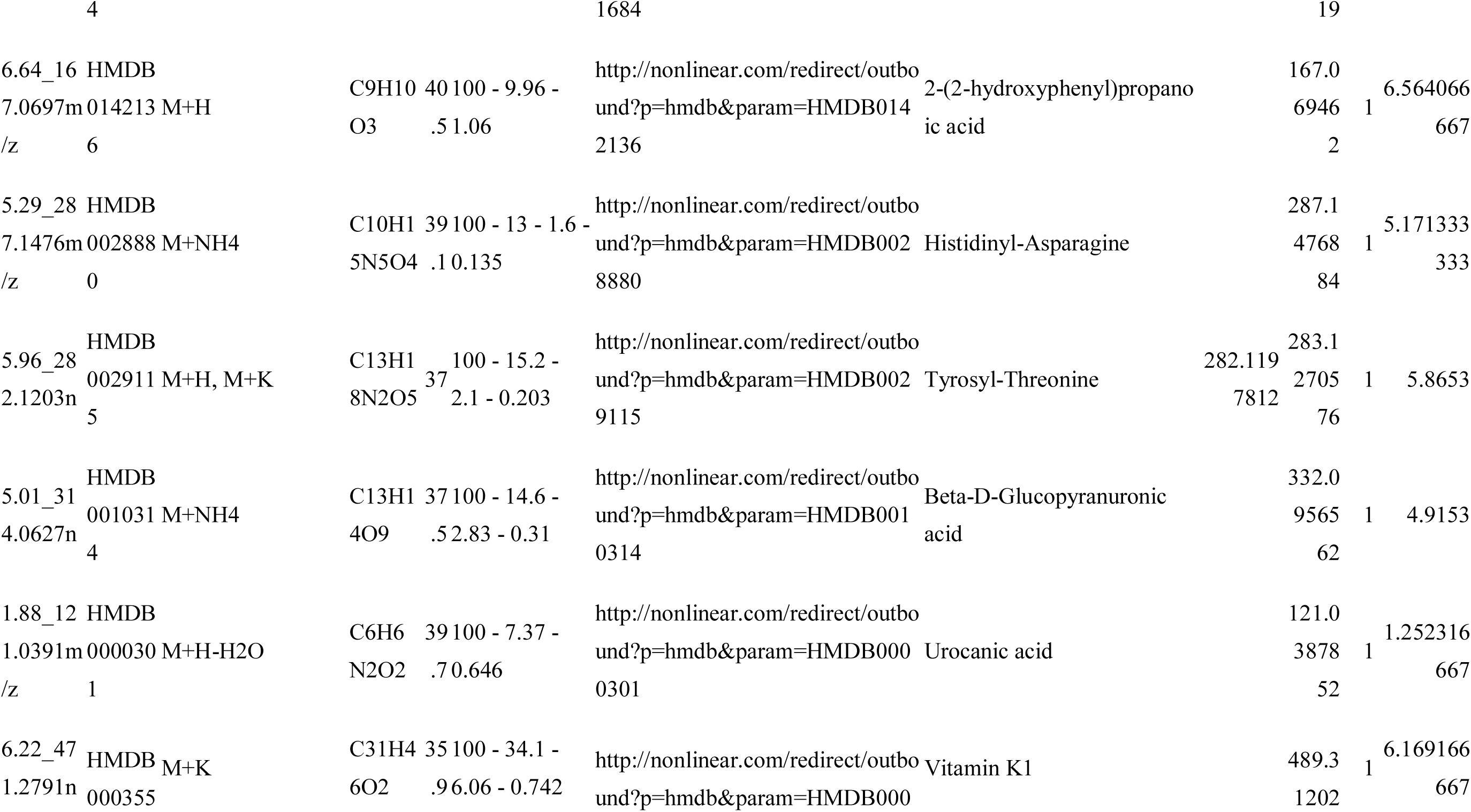
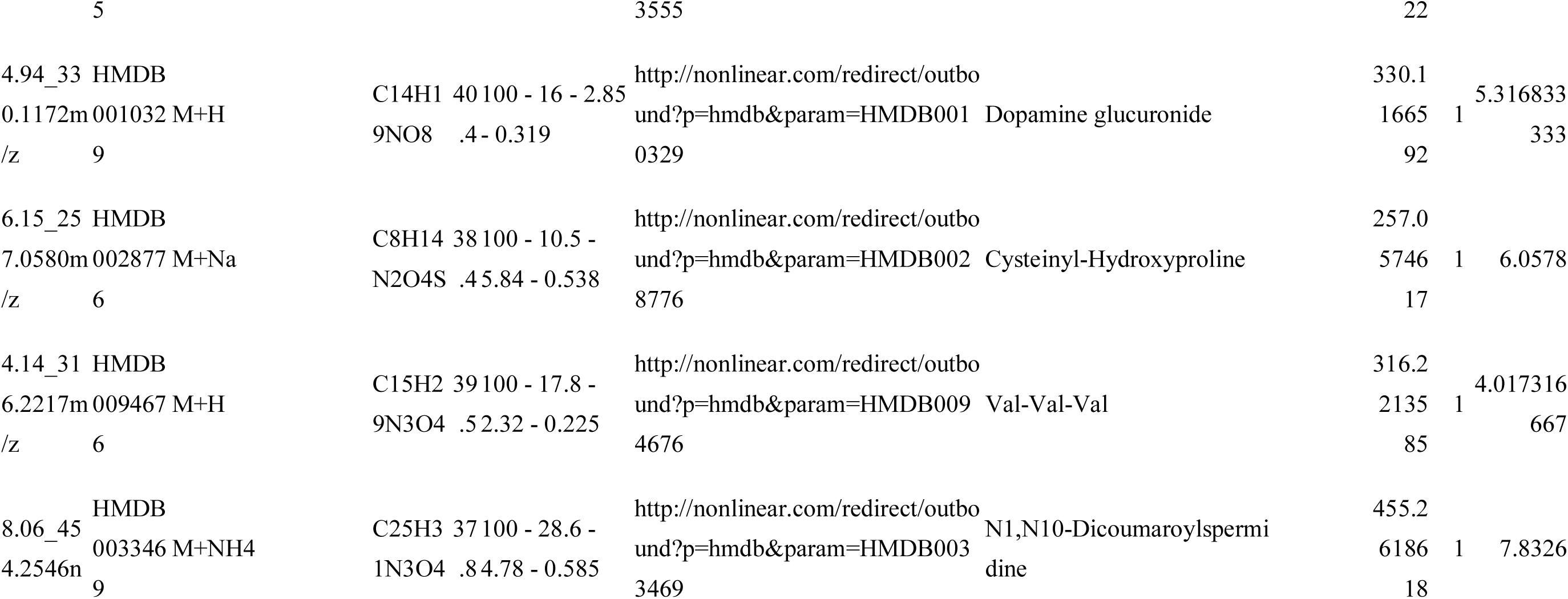
Detailed information of the identified 310 metabolites.

**Table S2.**
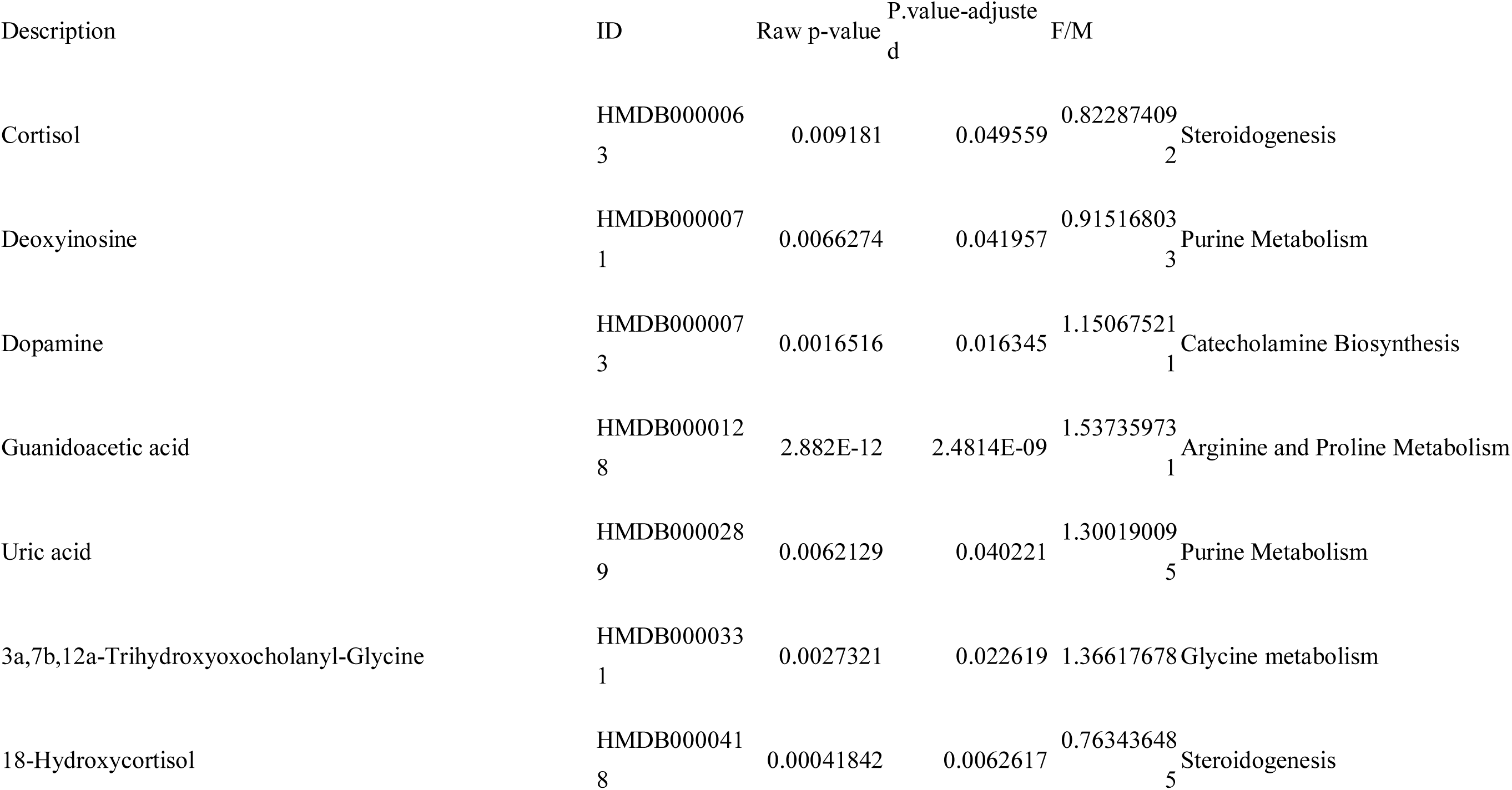
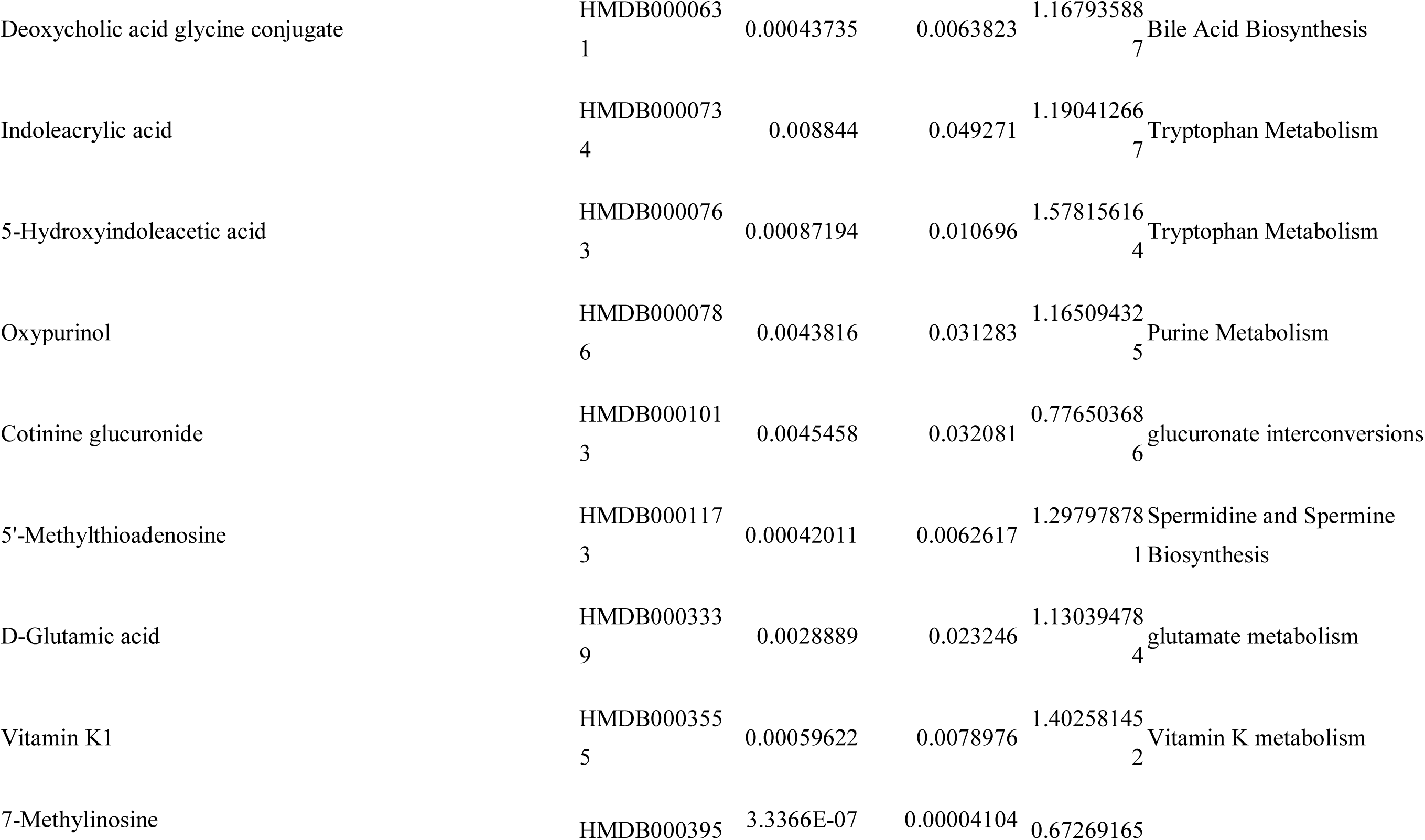
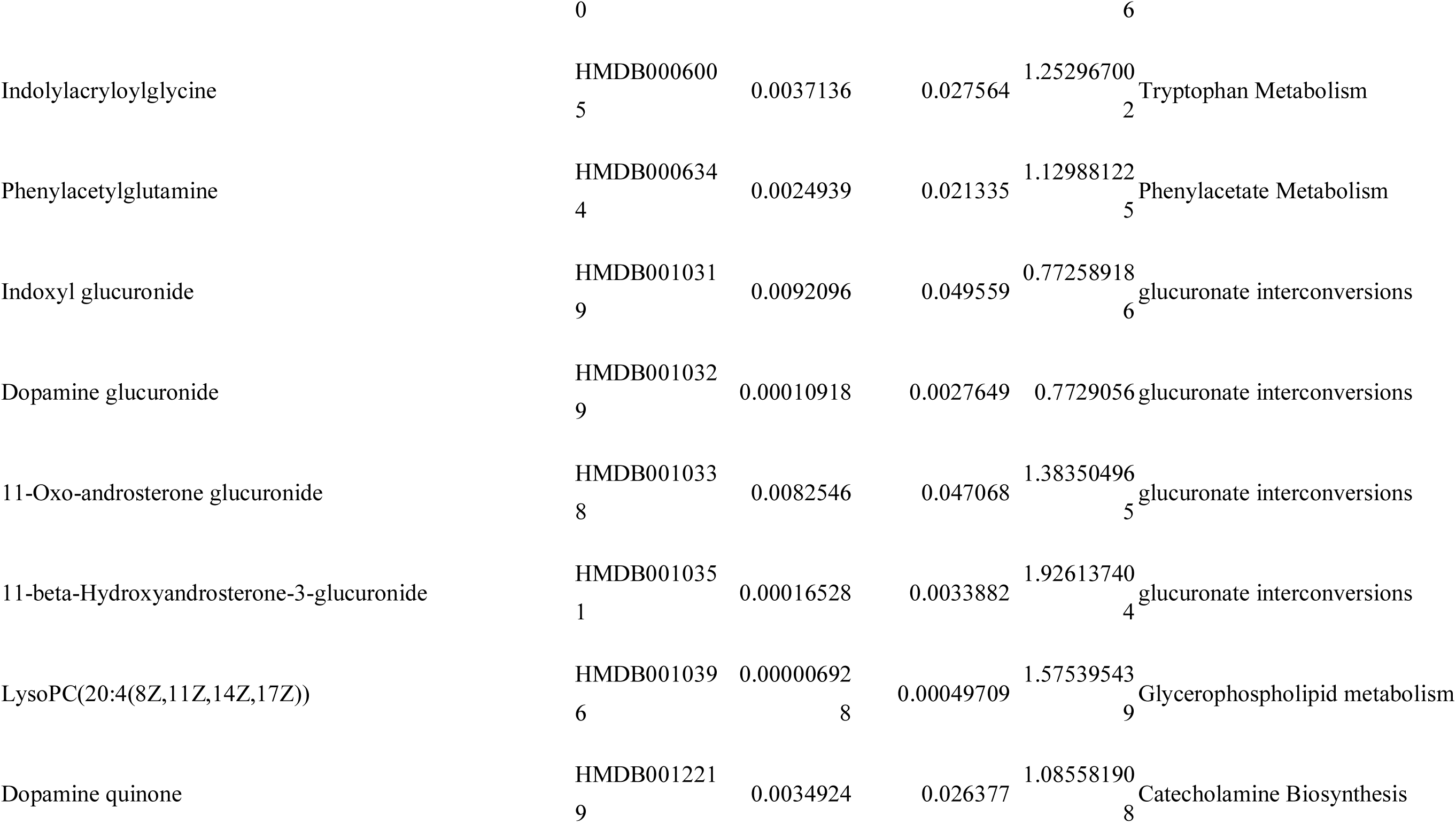
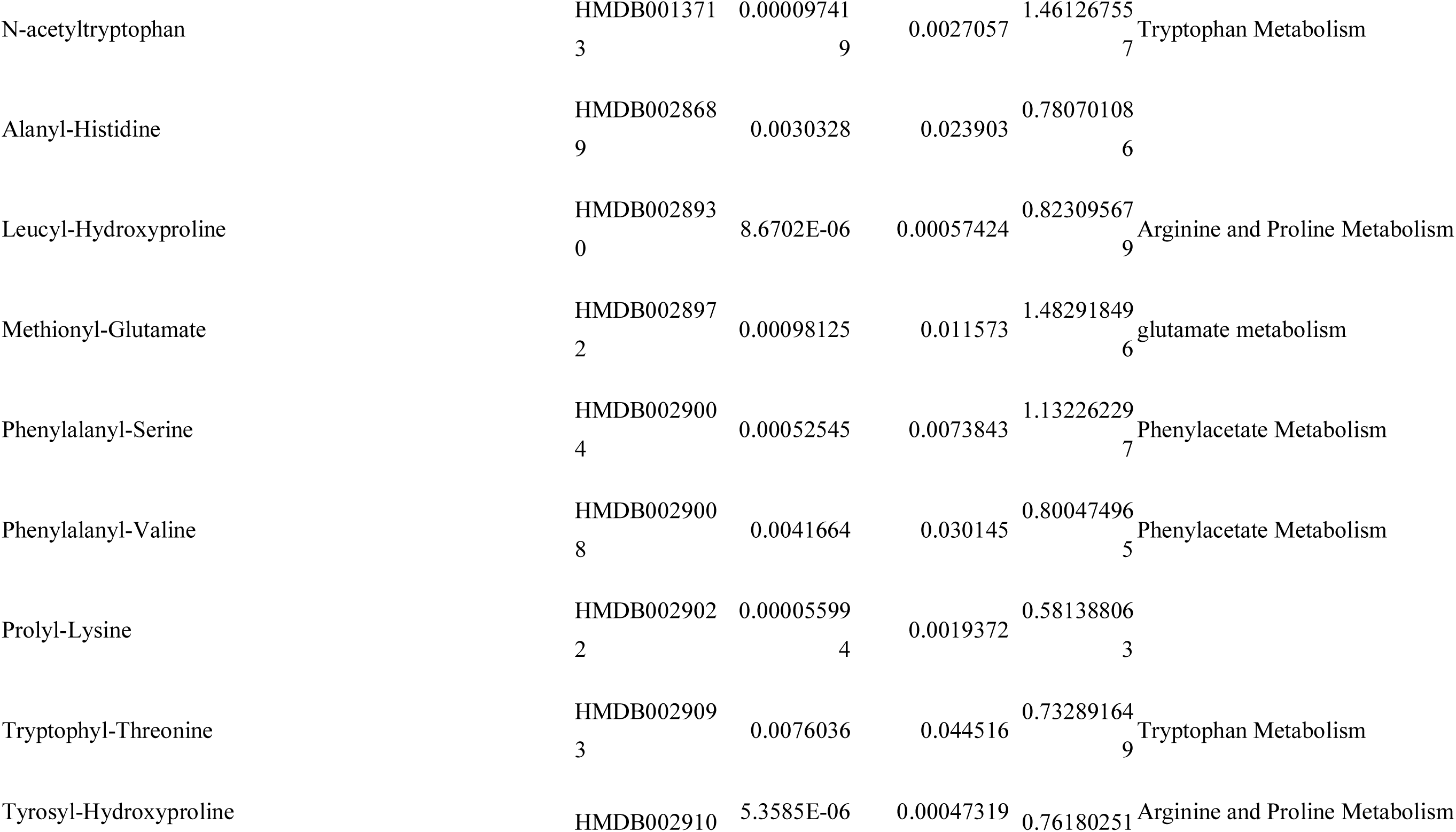
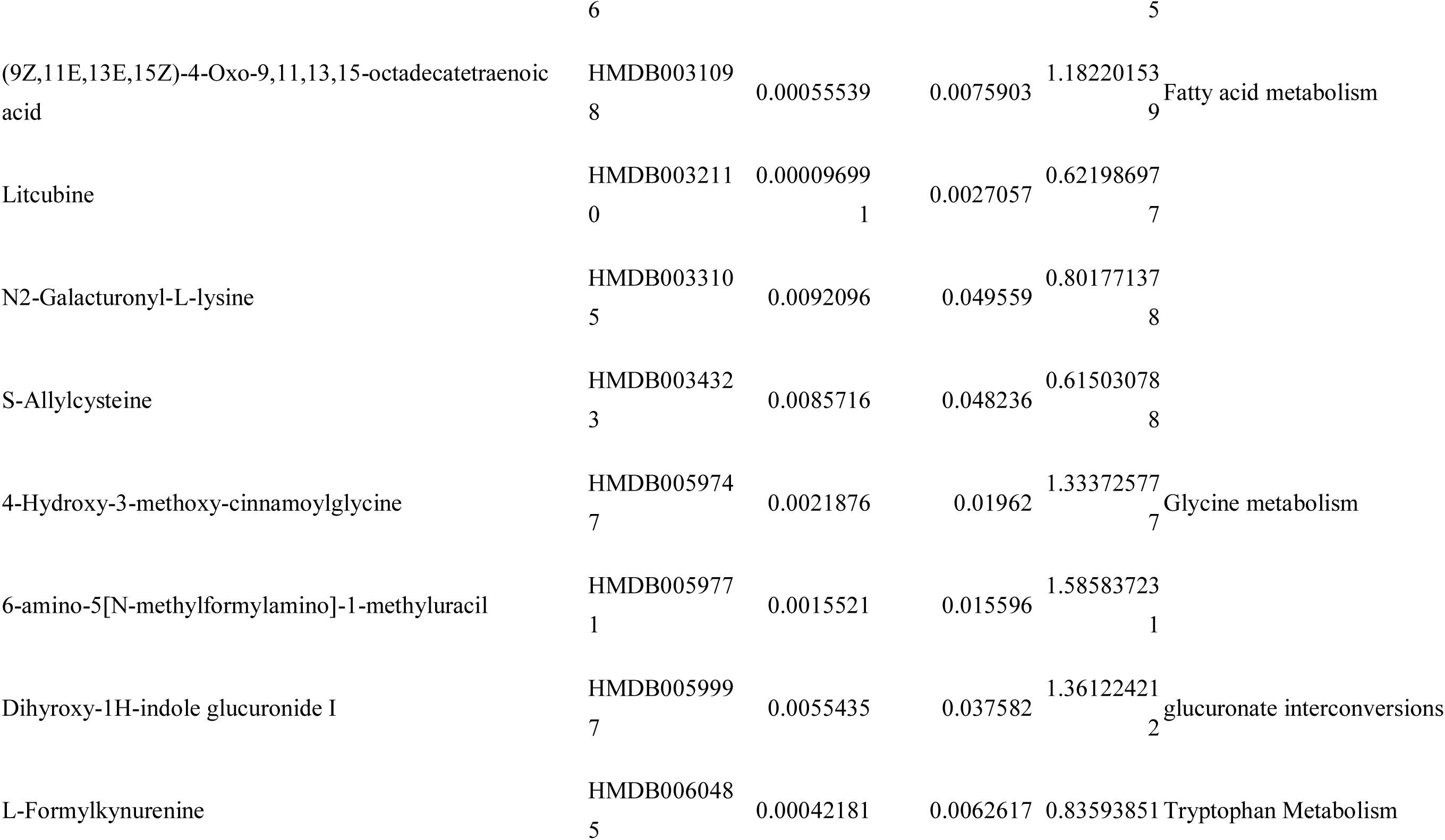
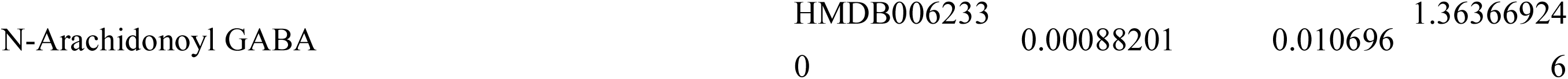
Gender differential metabolites in children population aged 1 to 18 years.

| Description | ID | Raw p-value | P.value-adjusted | F/M |  |
| --- | --- | --- | --- | --- | --- |
| Cortisol | HMDB000006<br>3 | 0.009181 | 0.049559 | 0.82287409<br>2 | Steroidogenesis |
| Deoxyinosine | HMDB000007<br>1 | 0.0066274 | 0.041957 | 0.91516803<br>3 | Purine Metabolism |
| Dopamine | HMDB000007<br>3 | 0.0016516 | 0.016345 | 1.15067521<br>1 | Catecholamine Biosynthesis |
| Guanidoacetic acid | HMDB000012<br>8 | 2.882E-12 | 2.4814E-09 | 1.53735973<br>1 | Arginine and Proline Metabolism |
| Uric acid | HMDB000028<br>9 | 0.0062129 | 0.040221 | 1.30019009<br>5 | Purine Metabolism |
| 3a,7b,12a-Trihydroxyoxocholanyl-Glycine | HMDB000033<br>1 | 0.0027321 | 0.022619 | 1.36617678 | Glycine metabolism |
| 18-Hydroxycortisol | HMDB000041<br>8 | 0.00041842 | 0.0062617 | 0.76343648<br>5 | Steroidogenesis |
| Deoxycholic acid glycine conjugate | HMDB000063<br>1 | 0.00043735 | 0.0063823 | 1.16793588<br>7 | Bile Acid Biosynthesis |
| Indoleacrylic acid | HMDB000073<br>4 | 0.008844 | 0.049271 | 1.19041266<br>7 | Tryptophan Metabolism |
| 5-Hydroxyindoleacetic acid | HMDB000076<br>3 | 0.00087194 | 0.010696 | 1.57815616<br>4 | Tryptophan Metabolism |
| Oxypurinol | HMDB000078<br>6 | 0.0043816 | 0.031283 | 1.16509432<br>5 | Purine Metabolism |
| Cotinine glucuronide | HMDB000101<br>3 | 0.0045458 | 0.032081 | 0.77650368<br>6 | glucuronate interconversions |
| 5'-Methylthioadenosine | HMDB000117<br>3 | 0.00042011 | 0.0062617 | 1.29797878<br>1 | Spermidine and Spermine Biosynthesis |
| D-Glutamic acid | HMDB000333<br>9 | 0.0028889 | 0.023246 | 1.13039478<br>4 | glutamate metabolism |
| Vitamin K1 | HMDB000355<br>5 | 0.00059622 | 0.0078976 | 1.40258145<br>2 | Vitamin K metabolism |
| 7-Methylinosine | HMDB000395 | 3.3366E-07 | 0.00004104 | 0.67269165 |  |
|  | 0 |  |  | 6 |  |
| Indolylacryloylglycine | HMDB000600<br>5 | 0.0037136 | 0.027564 | 1.25296700<br>2 | Tryptophan Metabolism |
| Phenylacetylglutamine | HMDB000634<br>4 | 0.0024939 | 0.021335 | 1.12988122<br>5 | Phenylacetate Metabolism |
| Indoxyl glucuronide | HMDB001031<br>9 | 0.0092096 | 0.049559 | 0.77258918<br>6 | glucuronate interconversions |
| Dopamine glucuronide | HMDB001032<br>9 | 0.00010918 | 0.0027649 | 0.7729056 | glucuronate interconversions |
| 11-Oxo-androsterone glucuronide | HMDB001033<br>8 | 0.0082546 | 0.047068 | 1.38350496<br>5 | glucuronate interconversions |
| 11-beta-Hydroxyandrosterone-3-glucuronide | HMDB001035<br>1 | 0.00016528 | 0.0033882 | 1.92613740<br>4 | glucuronate interconversions |
| LysoPC(20:4(8Z,11Z,14Z,17Z)) | HMDB001039<br>6 | 0.00000692<br>8 | 0.00049709 | 1.57539543<br>9 | Glycerophospholipid metabolism |
| Dopamine quinone | HMDB001221<br>9 | 0.0034924 | 0.026377 | 1.08558190<br>8 | Catecholamine Biosynthesis |
| N-acetyltryptophan | HMDB001371<br>3 | 0.00009741<br>9 | 0.0027057 | 1.46126755<br>7 | Tryptophan Metabolism |
| Alanyl-Histidine | HMDB002868<br>9 | 0.0030328 | 0.023903 | 0.78070108<br>6 |  |
| Leucyl-Hydroxyproline | HMDB002893<br>0 | 8.6702E-06 | 0.00057424 | 0.82309567<br>9 | Arginine and Proline Metabolism |
| Methionyl-Glutamate | HMDB002897<br>2 | 0.00098125 | 0.011573 | 1.48291849<br>6 | glutamate metabolism |
| Phenylalanyl-Serine | HMDB002900<br>4 | 0.00052545 | 0.0073843 | 1.13226229<br>7 | Phenylacetate Metabolism |
| Phenylalanyl-Valine | HMDB002900<br>8 | 0.0041664 | 0.030145 | 0.80047496<br>5 | Phenylacetate Metabolism |
| Prolyl-Lysine | HMDB002902<br>2 | 0.00005599<br>4 | 0.0019372 | 0.58138806<br>3 |  |
| Tryptophyl-Threonine | HMDB002909<br>3 | 0.0076036 | 0.044516 | 0.73289164<br>9 | Tryptophan Metabolism |
| Tyrosyl-Hydroxyproline | HMDB002910 | 5.3585E-06 | 0.00047319 | 0.76180251 | Arginine and Proline Metabolism |
|  | 6 |  |  | 5 |  |
| (9Z,11E,13E,15Z)-4-Oxo-9,11,13,15-octadecatetraenoic acid | HMDB003109<br>8 | 0.00055539 | 0.0075903 | 1.18220153<br>9 | Fatty acid metabolism |
| Litcubine | HMDB003211<br>0 | 0.00009699<br>1 | 0.0027057 | 0.62198697<br>7 |  |
| N2-Galacturonyl-L-lysine | HMDB003310<br>5 | 0.0092096 | 0.049559 | 0.80177137<br>8 |  |
| S-Allylcysteine | HMDB003432<br>3 | 0.0085716 | 0.048236 | 0.61503078<br>8 |  |
| 4-Hydroxy-3-methoxy-cinnamoylglycine | HMDB005974<br>7 | 0.0021876 | 0.01962 | 1.33372577<br>7 | Glycine metabolism |
| 6-amino-5[N-methylformylamino]-1-methyluracil | HMDB005977<br>1 | 0.0015521 | 0.015596 | 1.58583723<br>1 |  |
| Dihydroxy-1H-indole glucuronide I | HMDB005999<br>7 | 0.0055435 | 0.037582 | 1.36122421<br>2 | glucuronate interconversions |
| L-Formylkynurenine | HMDB006048<br>5 | 0.00042181 | 0.0062617 | 0.83593851 | Tryptophan Metabolism |

|  |  |  |  |  |
| --- | --- | --- | --- | --- |
| N-Arachidonoyl GABA | HMDB006233 | 0.00088201 | 0.010696 | 1.36366924 |
|  | 0 |  |  | 6 |

**Table S3.**
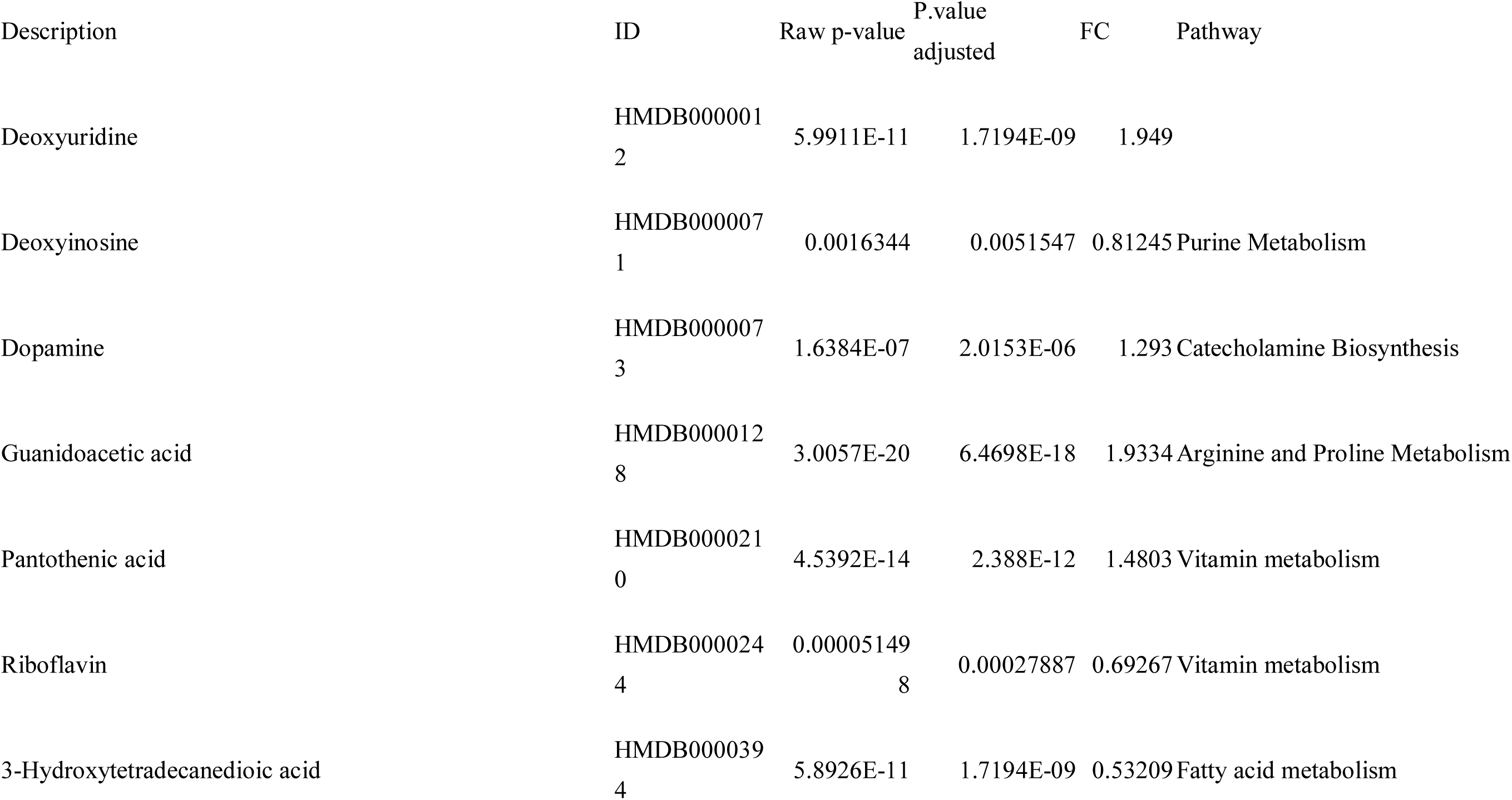
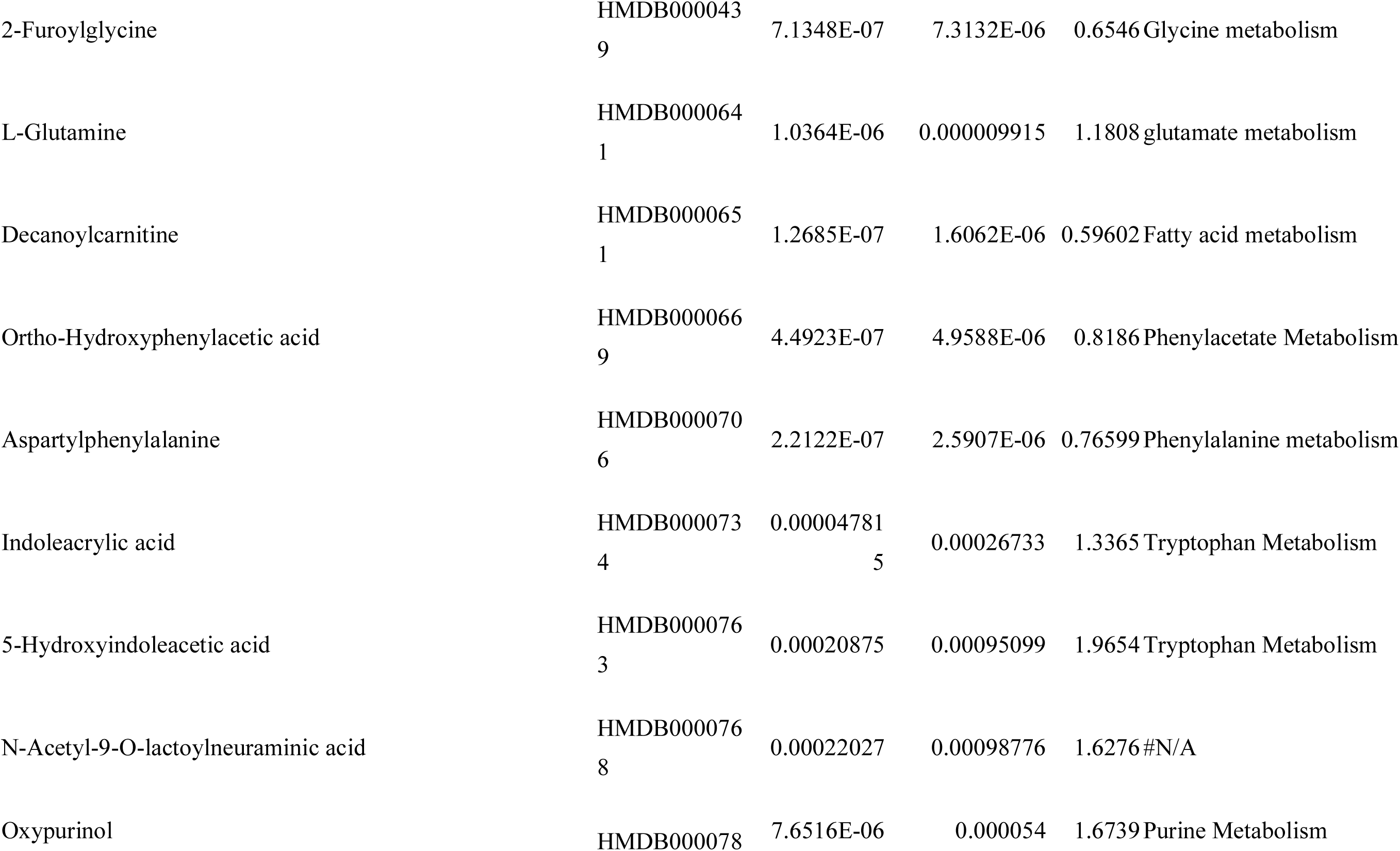
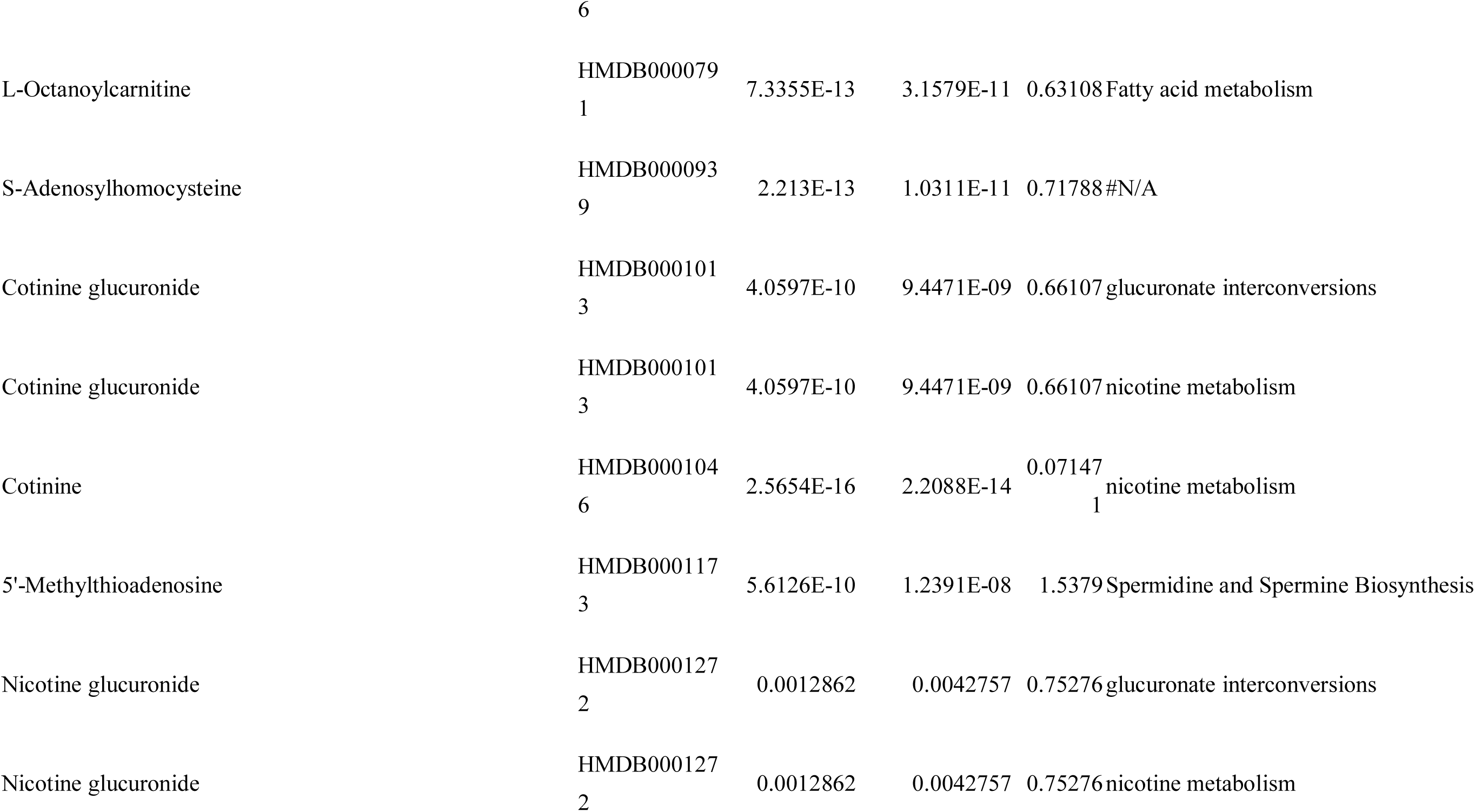
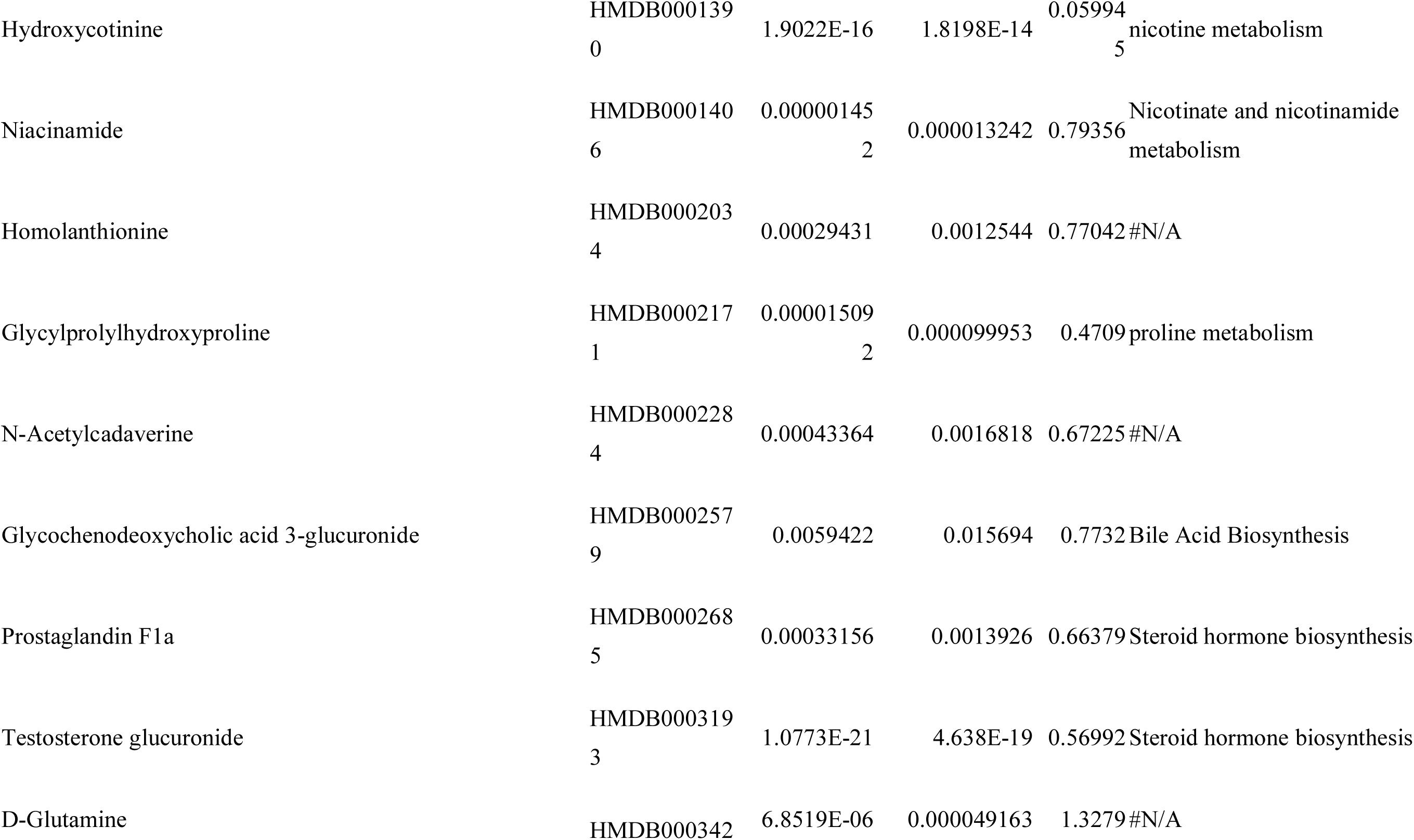
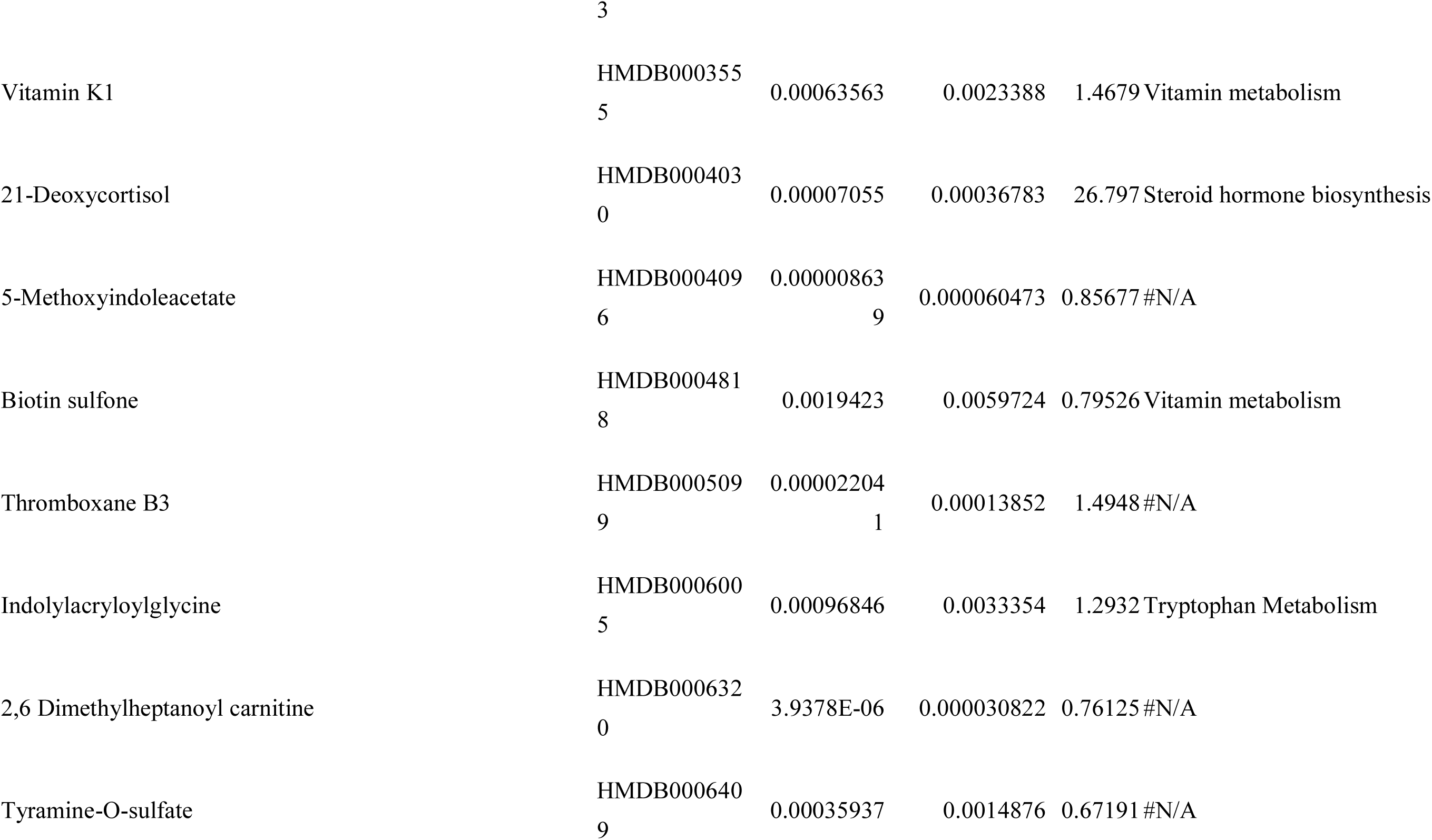
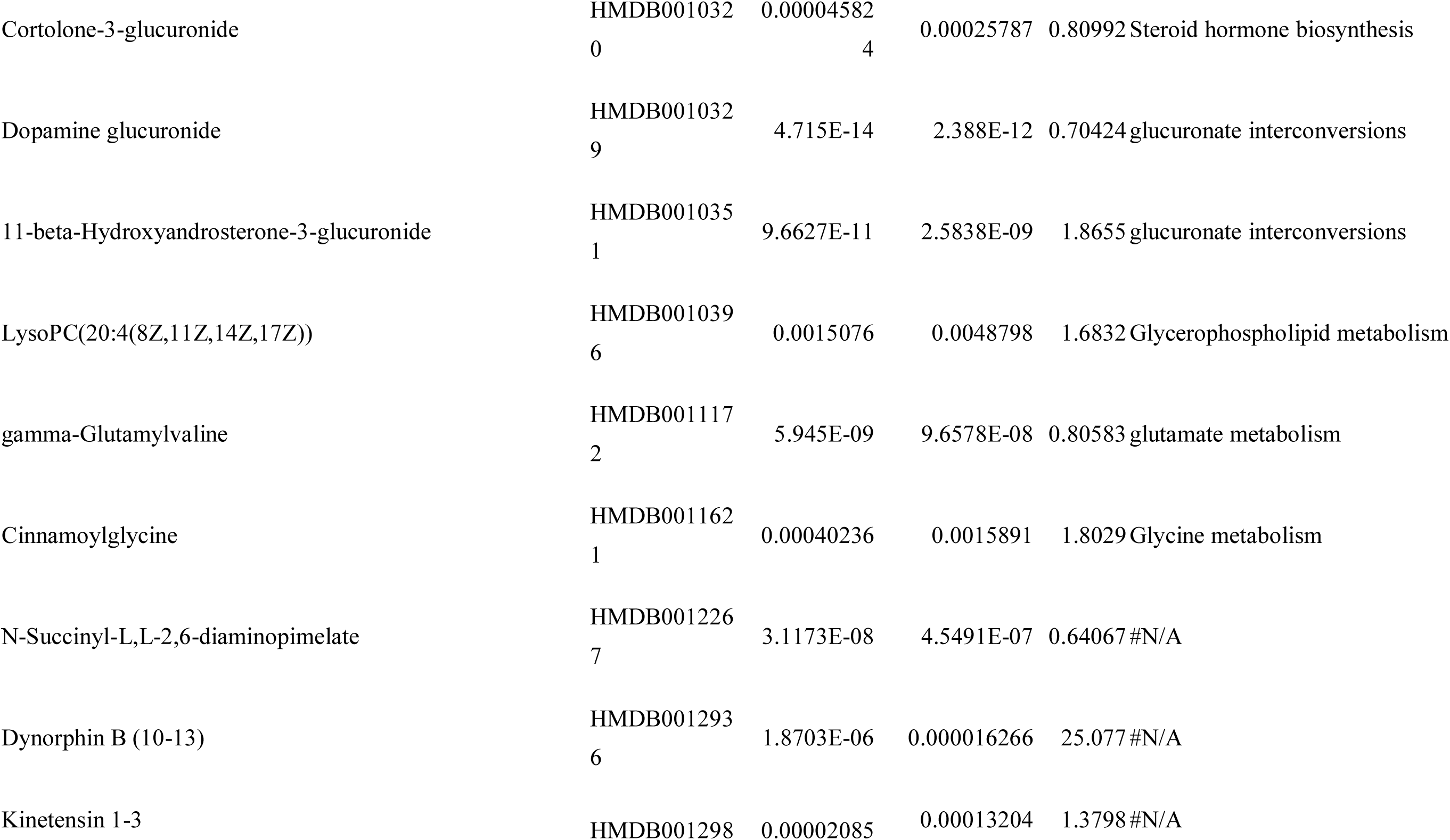
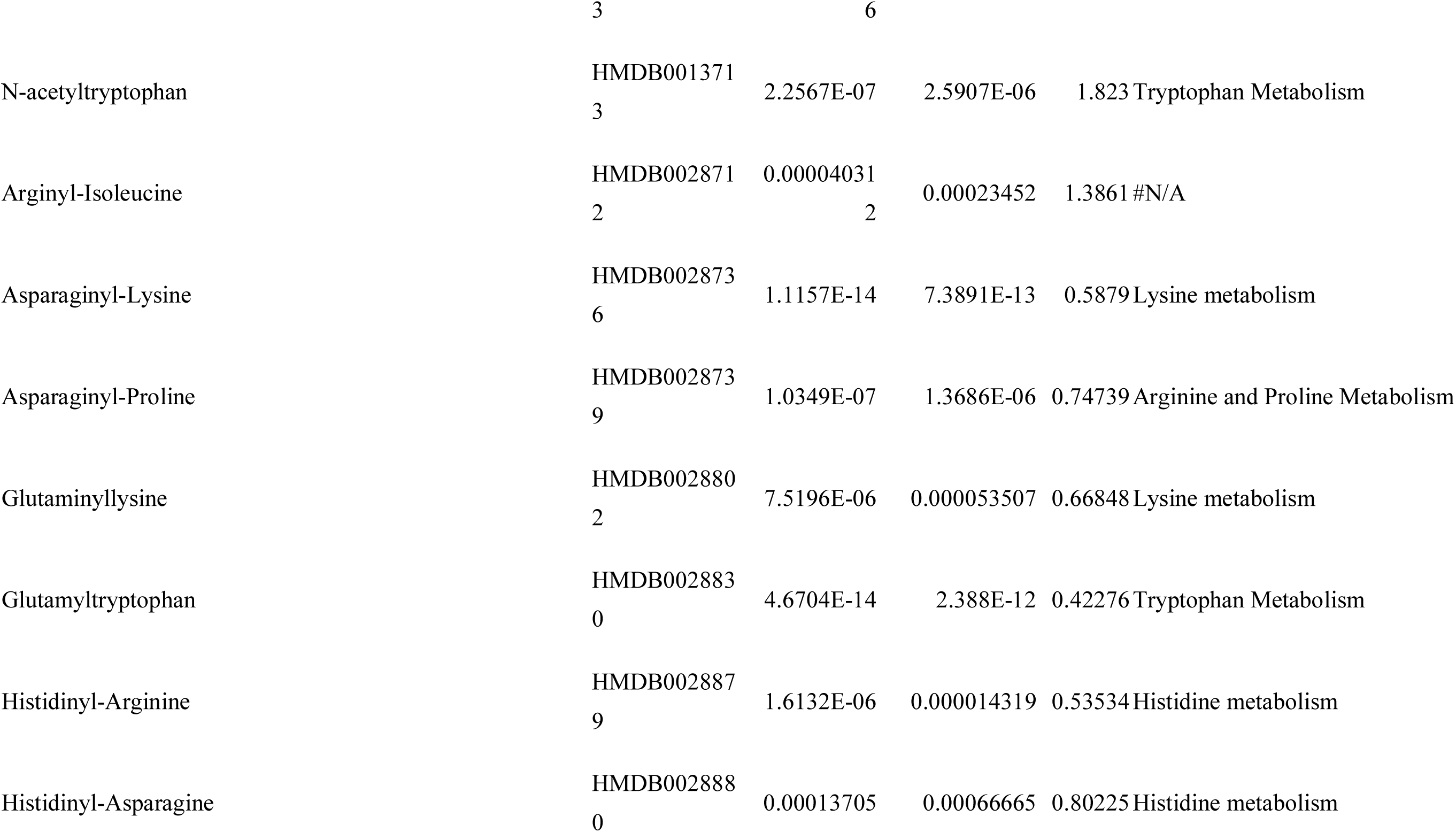
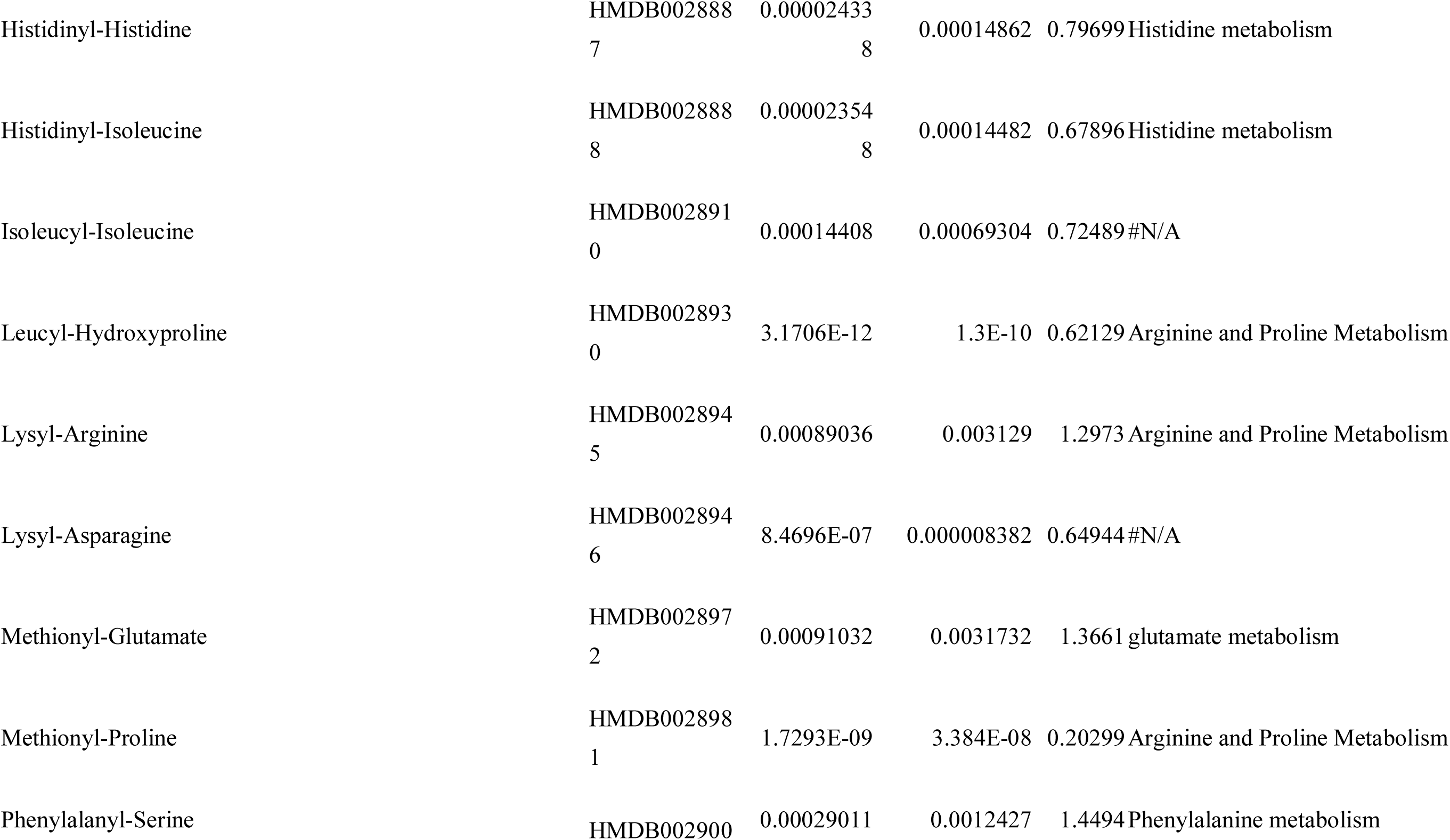
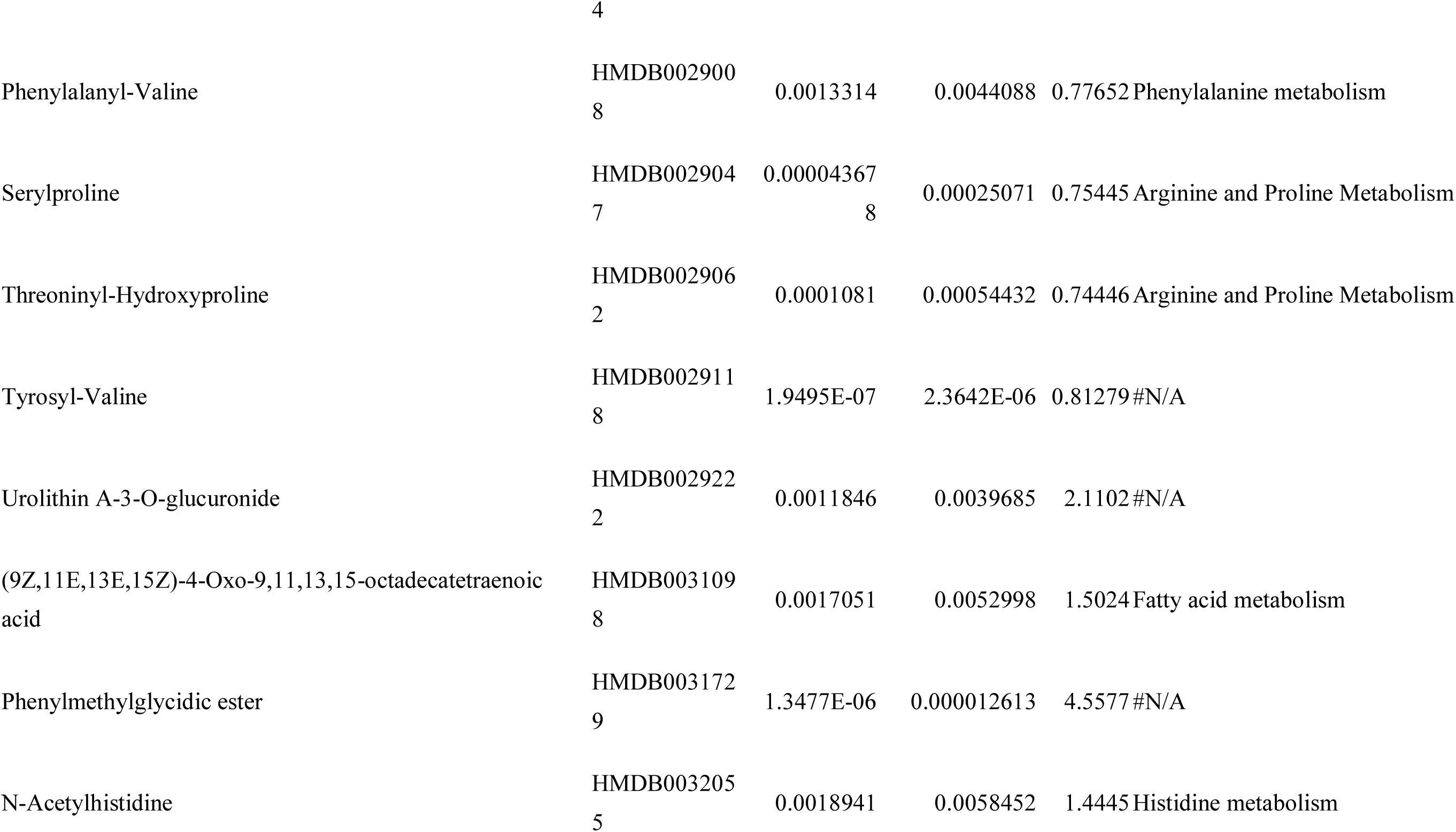
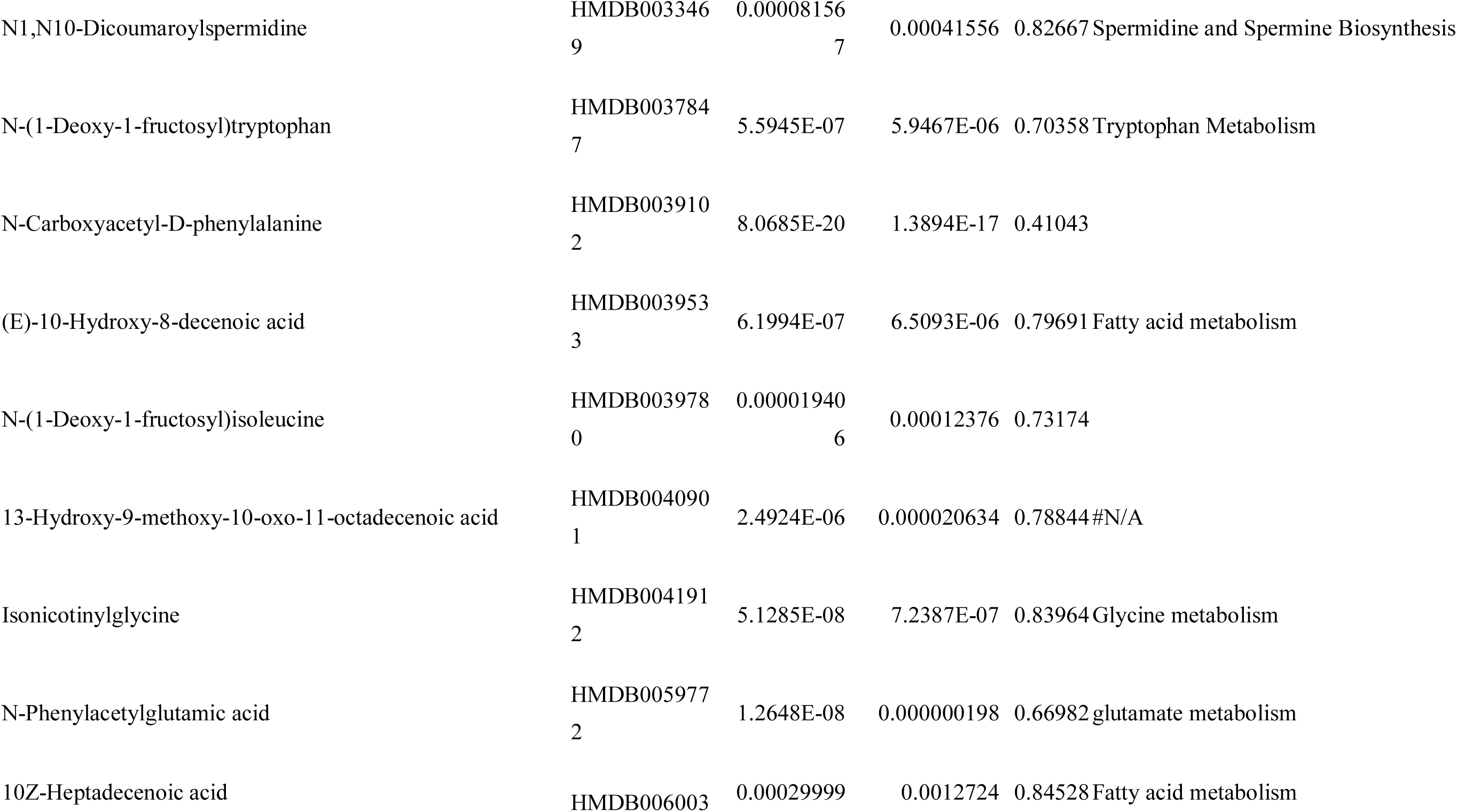
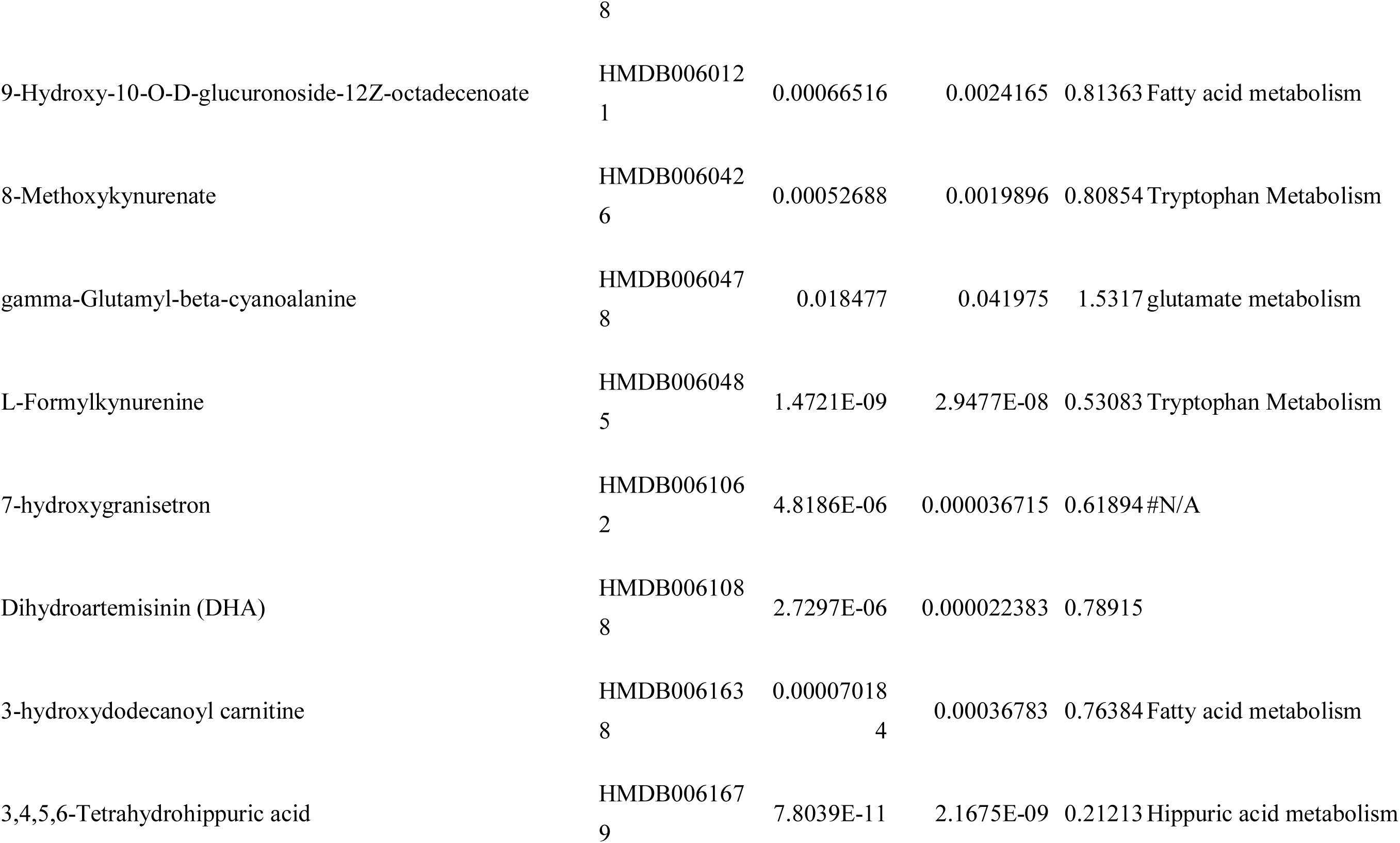
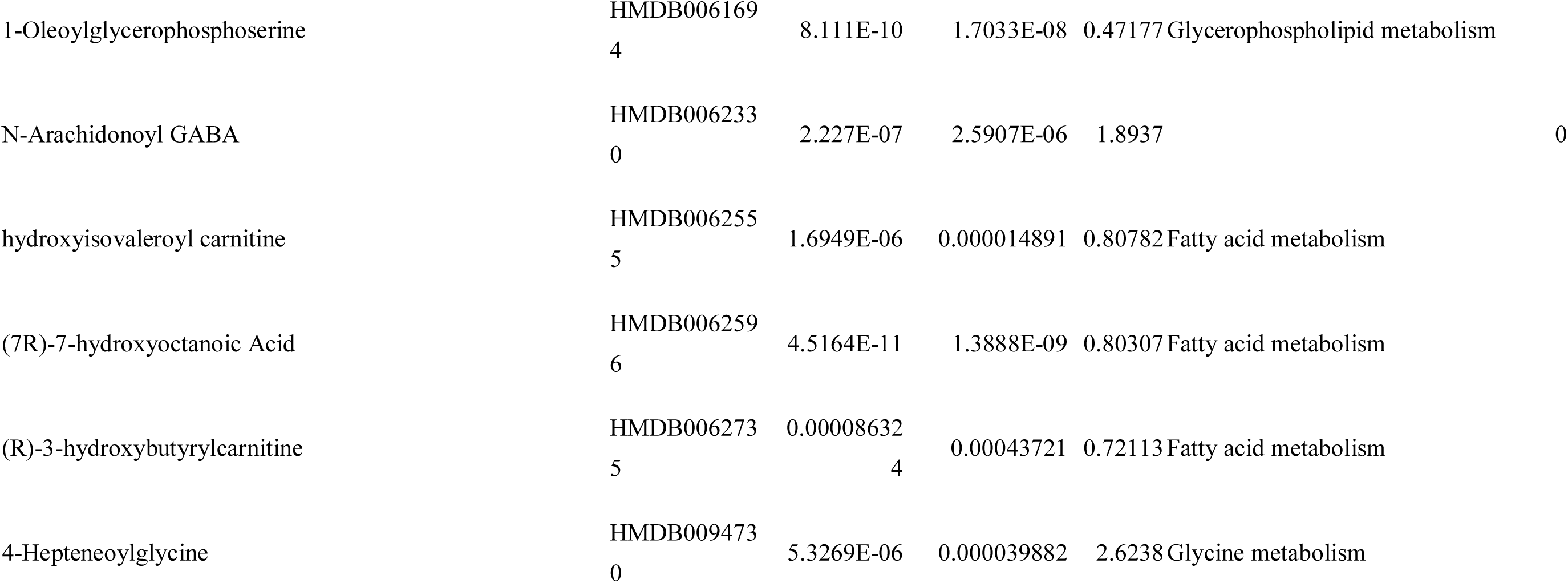
Gender differential metabolites in adults population.

**Table S4.** Gender and age dependent metabolism pathways.

| **Gender-dependent metabolism pathways** |  |
| --- | --- |
| **Factors** | **Differential metabolism pathways** |
| 1-6 | Vitamin k metabolism |
| 7-12 | Riboflavin metabolism, Steroidogenesis , Arginine and proline metabolism, glycine and serine metabolism |
| 13-18 | Riboflavin metabolism, Androgen and estrogen metabolism , Pantothenate and CoA biosynthesis, caffeine metabolism, Arginine and proline metabolism, Pterine biosynthesis, urea cycle |
| 20-30 | Methylhistine metabolism, Vitamine B6 metabolism, Biotin metabolism, Phosphatidylcholine biosynthesis, Ammonia recycling, Alanine metabolism, Nicotinate and nicotinamide metabolism , Vitamin k metabolism, Spermidine and spermine biosynthesis, Pterine biosynthesis |
| 31-50 | Riboflavin metabolism, Beta-oxidation of short chain fatty acids , Spermidine and spermine biosynthesis, Pantothenate and CoA biosynthesis, Pterine biosynthesis, beta-alanine metabolism |
| >50 | Beta-oxidation of short chain fatty acids, Androgen and estrogen metabolism , Vitamin k metabolism, Pterine biosynthesis, Arginine and proline metabolism |

| Age-dependent metabolism pathways |  |
| --- | --- |
| Age-boys | Tryptophan metabolism, Histidine metabolism, Riboflavin metabolism, Pantothenate and CoA biosynthesis, Alanine metabolism, Fatty acid biosynthesis |
| Age-girls | Histidine metabolism ,Purine metabolism ,Steroid hormone biosynthesis ,Riboflavin metabolism ,Pantothenate and CoA biosynthesis, Fatty acid biosynthesis |
| Age-males | Steroid hormone biosynthesis, Riboflavin metabolism, Tryptophan metabolism, Pantothenate and CoA biosynthesis, Amino acid metabolism, Purine metabolism, Pyrimidine metabolism |
| Age-females | Purine metabolism, Steroid hormone biosynthesis, Caffeine metabolism, Pyrimidine metabolism, Pantothenate and CoA biosynthesis, Amino acid metabolism |

**Table S5.** Age differential metabolites in boys.

| Compound | ID | VIP | Raw p.value | P-value adjusted |
| --- | --- | --- | --- | --- |
| 10-Hydroxy-octadec-12Z-enoate-9-beta-D-glucuronide | HMDB0060120 | 1.83762 | 4.84E-15 | 1.1906E-13 |
| 10Z-Heptadecenoic acid | HMDB0060038 | 2.26015 | 1.8146E-28 | 1.953E-26 |
| 11-beta-Hydroxyandosterone-3-glucuronide | HMDB0010351 | 1.5088 | 2.0918E-10 | 2.1699E-09 |
| 11-Oxo-androsterone glucuronide | HMDB0010338 | 2.06964 | 9.115E-18 | 3.27E-16 |
| 12-Hydroxydodecanoic acid | HMDB0002059 | 1.03311 | 0.000088107 | 0.00027386 |
| 12-Hydroxyhexadecanoic acid | HMDB0112190 | 1.5886 | 3.5655E-08 | 2.3615E-07 |
| 12-Methyltridecanoic acid | HMDB0031072 | 1.30581 | 2.6268E-09 | 2.2617E-08 |
| 12-oxo-20-dihydroxy-leukotriene B4 | HMDB0012551 | 1.46376 | 0.026547 | 0.04476 |
| 14,15-DiHETrE | HMDB0002265 | 1.33567 | 0.00024928 | 0.00069236 |
| 1-Methylhistidine | HMDB0000001 | 1.59163 | 1.4388E-18 | 5.6311E-17 |
| 1-Oleoylglycerophosphoserine | HMDB0061694 | 1.36393 | 1.1443E-13 | 2.3458E-12 |
| 2-Amino-4-ethoxy-3-hydroxybutanoic acid | HMDB0032862 | 1.04644 | 0.000085252 | 0.00026595 |
| 2-Aminomuconic acid | HMDB0001241 | 1.06575 | 0.0013876 | 0.0032733 |
| 2-Furoylglycine | HMDB0000439 | 1.177660 | 0.000024384 | 0.000088212 |
| 2-octenoylglycine | HMDB0094801 | 1.219020 | 0.000012341 | 0.00004608 |
| 3,4,5,6-Tetrahydrohippuric acid | HMDB0061679 | 1.02523 | 6.5219E-06 | 0.00002674 |
| 3a,7b,12a-Trihydroxyoxocholanyl-Glycine | HMDB0000331 | 1.00857 | 0.023923 | 0.040869 |
| 3-alpha-hydroxy-5-alpha-androstane-17-one 3-D-glucuronide | HMDB0010365 | 3.07168 | 5.5E-43 | 4.7355E-40 |
| 3-Hydroxydodecanedioic acid | HMDB0000413 | 1.04132 | 1.0099E-15 | 2.8357E-14 |
| 3-hydroxydodecanoyl carnitine | HMDB0061638 | 1.18864 | 0.0057302 | 0.011609 |
| 3-Hydroxytetradecanedioic acid | HMDB0000394 | 1.35272 | 8.6761E-16 | 2.5759E-14 |
| 3-Methoxytyrosine | HMDB0001434 | 1.02954 | 4.1996E-06 | 0.000018638 |
| 4-Hepteneoylglycine | HMDB0094730 | 1.15123 | 1.4833E-08 | 1.0732E-07 |
| 5-Hydroxyindoleacetic acid | HMDB0000763 | 1.05159 | 0.00013506 | 0.00040802 |
| 5-Methylcytidine | HMDB0000982 | 1.9013 | 2.8118E-09 | 2.3735E-08 |
| 5'-Methylthioadenosine | HMDB0001173 | 1.031350 | 0.000041448 | 0.00014105 |
| 5Z-Dodecenoic acid | HMDB0000529 | 1.17921 | 2.4877E-08 | 1.6999E-07 |
| 6-octenoylglycine | HMDB0094805 | 1.03805 | 0.00022408 | 0.00062844 |
| 7-Aminomethyl-7-carbaguanine | HMDB0011690 | 1.52211 | 1.681E-09 | 1.4921E-08 |
| 7-hydroxygranisetron | HMDB0061062 | 1.09752 | 8.9435E-06 | 0.000035161 |
| 7Z,10Z-Hexadecadienoic acid | HMDB0000477 | 1.27628 | 2.2169E-07 | 1.2236E-06 |
| 8-Hydroxy-5,6-octadienoic acid | HMDB0031101 | 1.20208 | 9.4714E-08 | 5.7027E-07 |
| 8-Hydroxy-7-methylguanine | HMDB0006037 | 1.08802 | 5.4212E-06 | 0.000022881 |
| 8-Methoxykynurenate | HMDB0060426 | 1.53238 | 2.9697E-06 | 0.000013601 |
| 9-Hydroxy-10-O-D-glucuronoside-12Z-octadecenoate | HMDB0060121 | 1.6162 | 1.2387E-12 | 2.2219E-11 |
| 9-peroxy-5Z,7E,11Z,14Z-eicosatetraenoate | HMDB0060098 | 1.14289 | 0.0015792 | 0.0036849 |
| alpha-CEHC glucuronide | HMDB0062445 | 1.17026 | 1.7382E-08 | 1.2167E-07 |
| Androstenedione | HMDB0000053 | 1.50982 | 3.319E-07 | 0.000001764 |
| Androstenol | HMDB0005935 | 2.08427 | 1.2664E-31 | 2.1808E-29 |
| AsparaginyL-Lysine | HMDB0028736 | 1.63496 | 1.1422E-21 | 7.3661E-20 |
| Beta-Citryl-L-glutamic acid | HMDB0013220 | 1.34012 | 4.7718E-11 | 6.0419E-10 |
| Cotinine | HMDB0001046 | 1.68957 | 7.8077E-07 | 3.8635E-06 |
| Cotinine glucuronide | HMDB0001013 | 1.17367 | 0.000010621 | 0.000040463 |
| Cytosine | HMDB0000630 | 1.83326 | 6.8114E-11 | 8.26E-10 |
| Deoxyuridine | HMDB0000012 | 1.25522 | 0.00067975 | 0.0017063 |
| Dihydroxy-1H-indole glucuronide I | HMDB0059997 | 1.1680 | 0.000008481 | 0.000033964 |
| DL-Dopa | HMDB0000609 | 1.40935 | 0.00019339 | 0.00055504 |
| Dodecanoic acid | HMDB0000638 | 1.17673 | 4.7593E-07 | 2.4247E-06 |
| Dopamine glucuronide | HMDB0010329 | 1.49323 | 6.5033E-22 | 4.6661E-20 |
| Eicosapentaenoic acid | HMDB0001999 | 1.48613 | 2.9274E-12 | 4.7557E-11 |
| gamma-Glutamyl-beta-cyanoalanine | HMDB0060478 | 1.2691 | 0.020285 | 0.03557 |
| Glutamate, gamma-methyl ester | HMDB0061715 | 1.02066 | 0.0002545 | 0.00070458 |
| Glutamylisoleucine | HMDB0028822 | 1.09779 | 2.5291E-15 | 6.4262E-14 |
| Glycochenodeoxycholic acid 3-glucuronide | HMDB0002579 | 1.59991 | 3.134E-11 | 4.2831E-10 |
| Glycyl-Histidine | HMDB0028843 | 2.08443 | 5.3763E-11 | 6.7086E-10 |
| HistidinyI-Arginine | HMDB0028879 | 1.18299 | 0.0010671 | 0.0025592 |
| HistidinyI-Asparagine | HMDB0028880 | 1.14463 | 0.00001869 | 0.000068478 |
| HistidinyI-Histidine | HMDB0028887 | 1.12599 | 3.4594E-11 | 4.5824E-10 |
| Hydroxycotinine | HMDB0001390 | 2.1527 | 7.5805E-10 | 7.1723E-09 |
| Isoleucyl-Leucine | HMDB0028911 | 1.47108 | 2.1018E-06 | 9.8349E-06 |
| Kinetensin 1-3 | HMDB0012983 | 1.24301 | 8.9858E-08 | 5.4871E-07 |
| Kynuramine | HMDB0012246 | 1.43135 | 5.8681E-06 | 0.000024302 |
| L-Octanoylcarnitine | HMDB0000791 | 1.33 | 1.021E-15 | 2.8357E-14 |
| Lysyl-Asparagine | HMDB0028946 | 1.29517 | 6.9111E-13 | 1.266E-11 |
| Methionyl-Glutamate | HMDB0028972 | 1.02282 | 0.00015888 | 0.00047498 |
| N1,N10-Dicoumaroylspermidine | HMDB0033469 | 1.90388 | 1.2594E-14 | 2.9306E-13 |
| N-Acetyl-9-O-lactoylneuraminic acid | HMDB0000768 | 1.35408 | 9.9726E-06 | 0.000038504 |
| N-Acetylaspartylglutamic acid | HMDB0001067 | 2.45464 | 1.2075E-16 | 3.8506E-15 |
| N-Acetylhistidine | HMDB0032055 | 1.41169 | 7.3513E-11 | 8.6705E-10 |
| N-acetyltryptophan | HMDB0013713 | 1.3743 | 0.00046284 | 0.0012039 |
| NAPQI | HMDB0060946 | 1.35511 | 3.0133E-08 | 2.0429E-07 |
| Nicotine glucuronide | HMDB0001272 | 1.15856 | 3.638E-08 | 2.3911E-07 |
| N-Methylserotonin | HMDB0004369 | 1.57520 | 0.000012363 | 0.00004608 |
| N-Succinyl-L,L-2,6-diaminopimelate | HMDB0012267 | 1.52039 | 0.00078595 | 0.0019279 |
| Ortho-Hydroxyphenylacetic acid | HMDB0000669 | 1.30534 | 9.1085E-19 | 3.7345E-17 |
| Oxypurinol | HMDB0000786 | 1.96538 | 9.0519E-18 | 3.27E-16 |
| Pantothenic acid | HMDB0000210 | 1.68881 | 1.2139E-30 | 1.7419E-28 |
| Phenylalanyl-Serine | HMDB0029004 | 1.16889 | 9.9709E-06 | 0.000038504 |
| Phenylmethylglycidic ester | HMDB0031729 | 1.2105 | 2.1928E-08 | 1.5104E-07 |
| Prolyl-Histidine | HMDB0029019 | 1.41617 | 7.7156E-09 | 5.9314E-08 |
| Prolyl-Lysine | HMDB0029022 | 3.47662 | 0.0001833 | 0.00053137 |
| Prostaglandin F1a | HMDB0002685 | 1.26371 | 0.00050203 | 0.0012864 |
| Riboflavin | HMDB0000244 | 1.66616 | 1.1469E-09 | 1.0395E-08 |
| Serylmethionine | HMDB0029045 | 1.21992 | 4.569E-07 | 2.3463E-06 |
| Serylproline | HMDB0029047 | 1.3227 | 1.9059E-06 | 8.9673E-06 |
| Succinyladenosine | HMDB0000912 | 1.13645 | 2.7627E-23 | 2.1624E-21 |
| Testosterone glucuronide | HMDB0003193 | 2.60747 | 3.9103E-39 | 1.6834E-36 |
| Tetradecanedioic acid | HMDB0000872 | 1.21149 | 1.7632E-07 | 9.8579E-07 |
| Theobromine | HMDB0002825 | 2.0576 | 1.4417E-08 | 1.052E-07 |
| Threoninyl-Asparagine | HMDB0029056 | 1.13838 | 1.2422E-10 | 1.389E-09 |
| Threoninyl-Proline | HMDB0029069 | 1.51386 | 4.256E-14 | 9.396E-13 |
| Tryptophyl-Threonine | HMDB0029093 | 1.15239 | 0.00006214 | 0.00020114 |
| Tyramine-O-sulfate | HMDB0006409 | 1.17301 | 0.00020927 | 0.00059467 |
| Tyrosyl-Hydroxyproline | HMDB0029106 | 1.36858 | 1.1272E-19 | 4.8527E-18 |
| Tyrosyl-Isoleucine | HMDB0029108 | 1.09731 | 9.8293E-08 | 5.8771E-07 |
| Uric acid | HMDB0000289 | 1.32663 | 1.0148E-10 | 1.1497E-09 |
| Urocanic acid | HMDB0000301 | 1.90705 | 2.2177E-20 | 1.1934E-18 |
| Valyl-Proline | HMDB0029135 | 1.27558 | 0.00041336 | 0.001093 |

**Table S6.** Age differential metabolites in girls.

| Compound | ID | VIP | Raw p.value | P-value adjusted |
| --- | --- | --- | --- | --- |
| 10-Hydroxy-octadec-12Z-enoate-9-beta-D-glucuronide | HMDB0060120 | 1.60914 | 1.8513E-08 | 3.6024E-07 |
| 10Z-Heptadecenoic acid | HMDB0060038 | 2.39228 | 1.3827E-11 | 4.7619E-10 |
| 11-beta-Hydroxyandrosterone-3-glucuronide | HMDB0010351 | 2.97832 | 2.3661E-12 | 9.7011E-11 |
| 11-Oxo-androsterone glucuronide | HMDB0010338 | 2.61005 | 3.6029E-15 | 2.5851E-13 |
| 12-Hydroxyhexadecanoic acid | HMDB0112190 | 1.48123 | 1.2547E-07 | 1.9478E-06 |
| 12-Methyltridecanoic acid | HMDB0031072 | 1.09854 | 0.00037271 | 0.0020056 |
| 18-Hydroxycortisol | HMDB0000418 | 1.02747 | 0.0010453 | 0.0046064 |
| 1-Methylhistidine | HMDB0000001 | 1.16109 | 0.000040572 | 0.00028077 |
| 1-Oleoylglycerophosphoserine | HMDB0061694 | 1.19824 | 7.6236E-06 | 0.000068374 |
| 21-Deoxycortisol | HMDB0004030 | 1.91262 | 0.0075737 | 0.02203 |
| 2-Amino-4-ethoxy-3-hydroxybutanoic acid | HMDB0032862 | 1.63016 | 0.0046924 | 0.015019 |
| 3,4-O-Dimethylgallic acid | HMDB0041662 | 1.07151 | 0.0061282 | 0.018294 |
| 3-alpha-hydroxy-5-alpha-androstane-17-one 3-D-glucuronide | HMDB0010365 | 3.18703 | 3.3365E-18 | 4.7878E-16 |
| 3-Hydroxytetradecanedioic acid | HMDB0000394 | 1.2878 | 0.000035271 | 0.00025307 |
| 3-Methoxytyrosine | HMDB0001434 | 1.89109 | 1.1758E-06 | 0.000013868 |
| 4-Hepteneoylglycine | HMDB0094730 | 1.10599 | 1.2425E-07 | 1.9478E-06 |
| 4-Pyridoxic acid | HMDB0000017 | 1.18179 | 0.013044 | 0.034136 |
| 5-Methoxytryptophan | HMDB0002339 | 1.27878 | 0.01423 | 0.03625 |
| 5'-Methylthioadenosine | HMDB0001173 | 1.39918 | 0.000034571 | 0.00025013 |
| 5Z-Dodecenoic acid | HMDB0000529 | 1.1343 | 2.3111E-06 | 0.000024266 |
| 7-Methylinosine | HMDB0003950 | 1.13161 | 0.017853 | 0.042213 |
| 8-Hydroxy-5,6-octadienoic acid | HMDB0031101 | 1.51476 | 0.0051912 | 0.01602 |
| 8-Hydroxy-7-methylguanine | HMDB0006037 | 1.10587 | 0.000029818 | 0.00022132 |
| 9-Hydroxy-10-O-D-glucuronoside-12Z-octadecenoate | HMDB0060121 | 1.33978 | 4.9577E-06 | 0.000046398 |
| alpha-CEHC glucuronide | HMDB0062445 | 1.43568 | 1.0988E-06 | 0.00001314 |
| Androstenedione | HMDB0000053 | 1.45314 | 0.000014246 | 0.0001191 |
| Androstenol | HMDB0005935 | 2.06736 | 6.0374E-18 | 7.426E-16 |
| Arginyl-Histidine | HMDB0028711 | 1.00018 | 0.00065604 | 0.0033423 |
| Arginyl-Proline | HMDB0028717 | 1.05334 | 0.000090549 | 0.00056907 |
| Asparaginy-Lysine | HMDB0028736 | 1.41849 | 1.4162E-12 | 6.4177E-11 |
| Aspartylphenylalanine | HMDB0000706 | 1.18854 | 0.000040762 | 0.00028077 |
| Aspartyl-Valine | HMDB0028766 | 1.063 | 0.013112 | 0.034211 |
| Beta-Citryl-L-glutamic acid | HMDB0013220 | 1.6593 | 0.0014283 | 0.0058727 |
| Deoxyinosine | HMDB0000071 | 1.1994 | 0.0007458 | 0.0036485 |
| Dihydroxy-1H-indole glucuronide I | HMDB0059997 | 1.36962 | 6.073E-09 | 1.4132E-07 |
| Diphthamide | HMDB0060473 | 1.29704 | 0.012893 | 0.033948 |
| Dodecanoic acid | HMDB0000638 | 1.18643 | 0.00018726 | 0.0010749 |
| Dopamine glucuronide | HMDB0010329 | 1.02192 | 0.000013132 | 0.00011325 |
| Glutamate, gamma-methyl ester | HMDB0061715 | 1.59439 | 0.0050013 | 0.015716 |
| Glutamylisoleucine | HMDB0028822 | 1.05414 | 1.6456E-12 | 7.0843E-11 |
| Glycochenodeoxycholic acid 3-glucuronide | HMDB0002579 | 1.54222 | 1.108E-11 | 4.0989E-10 |
| Glycylprolylhydroxyproline | HMDB0002171 | 1.93795 | 0.000005065 | 0.000046892 |
| Guanine | HMDB0000132 | 1.54704 | 1.8828E-08 | 3.6024E-07 |
| Histidinyl-Asparagine | HMDB0028880 | 1.41162 | 1.7823E-06 | 0.000019182 |
| Histidinyl-Histidine | HMDB0028887 | 1.14755 | 5.0887E-11 | 1.4605E-09 |
| Kynuramine | HMDB0012246 | 1.59884 | 0.005122 | 0.015921 |
| Leucyl-Hydroxyproline | HMDB0028930 | 1.35912 | 2.2557E-07 | 3.3486E-06 |
| L-Octanoylcarnitine | HMDB0000791 | 1.46439 | 2.5628E-07 | 0.00000374 |
| L-prolyl-L-proline | HMDB0011180 | 1.12994 | 0.014197 | 0.03625 |
| Lysyl-Asparagine | HMDB0028946 | 1.20734 | 8.6141E-13 | 4.3628E-11 |
| Methionyl-Glutamate | HMDB0028972 | 1.52222 | 2.3694E-06 | 0.000024579 |
| N1,N10-Dicoumaroylspermidine | HMDB0033469 | 1.73211 | 2.3074E-09 | 5.6761E-08 |
| N2,N2-Dimethylguanosine | HMDB0004824 | 1.14419 | 0.00059672 | 0.0031138 |
| N-Acetyl-9-O-lactoylneuraminic acid | HMDB0000768 | 1.17607 | 0.000044645 | 0.0002994 |
| N-Acetylaspartylglutamic acid | HMDB0001067 | 2.10141 | 0.0052314 | 0.016087 |
| N-Acetylhistidine | HMDB0032055 | 1.84236 | 0.000096347 | 0.0005968 |
| NAPQI | HMDB0060946 | 1.1814 | 1.6278E-08 | 3.3371E-07 |
| Niacinamide | HMDB0001406 | 1.21 | 0.00082676 | 0.0038675 |
| Nicotine glucuronide | HMDB0001272 | 1.04692 | 0.00066969 | 0.0033918 |
| Ortho-Hydroxyphenylacetic acid | HMDB0000669 | 1.43558 | 2.6968E-11 | 8.2925E-10 |
| Oxypurinol | HMDB0000786 | 1.29738 | 0.000094858 | 0.00059183 |
| Pantothenic acid | HMDB0000210 | 1.28721 | 1.8378E-11 | 6.086E-10 |
| Phenylalanyl-Gamma-glutamate | HMDB0029009 | 1.28121 | 3.9878E-15 | 2.6411E-13 |
| Pivaloylcarnitine | HMDB0041993 | 1.01361 | 0.013907 | 0.035851 |
| Prostaglandin B2 | HMDB0004236 | 1.09062 | 0.0076578 | 0.0222 |
| Riboflavin | HMDB0000244 | 1.31822 | 0.0005718 | 0.0030019 |
| Serylproline | HMDB0029047 | 1.72547 | 0.01063 | 0.029019 |
| Succinyladenosine | HMDB0000912 | 1.12975 | 8.9784E-18 | 9.574E-16 |
| Testosterone glucuronide | HMDB0003193 | 2.63919 | 7.1437E-17 | 5.5916E-15 |
| Tetradecanedioic acid | HMDB0000872 | 1.10707 | 0.0096715 | 0.026876 |
| Threoninyl-Asparagine | HMDB0029056 | 1.2968 | 1.229E-10 | 3.4134E-09 |
| Threoninyl-Proline | HMDB0029069 | 1.74715 | 0.0010133 | 0.0044971 |
| Tyrosyl-Hydroxyproline | HMDB0029106 | 2.49219 | 1.2567E-20 | 3.6066E-18 |
| Uric acid | HMDB0000289 | 1.14038 | 0.010646 | 0.029019 |
| Urocanic acid | HMDB0000301 | 2.9307 | 1.0008E-17 | 9.574E-16 |
| Vitamin K1 | HMDB0003555 | 1.09069 | 0.000041184 | 0.00028142 |

**Table S7.**
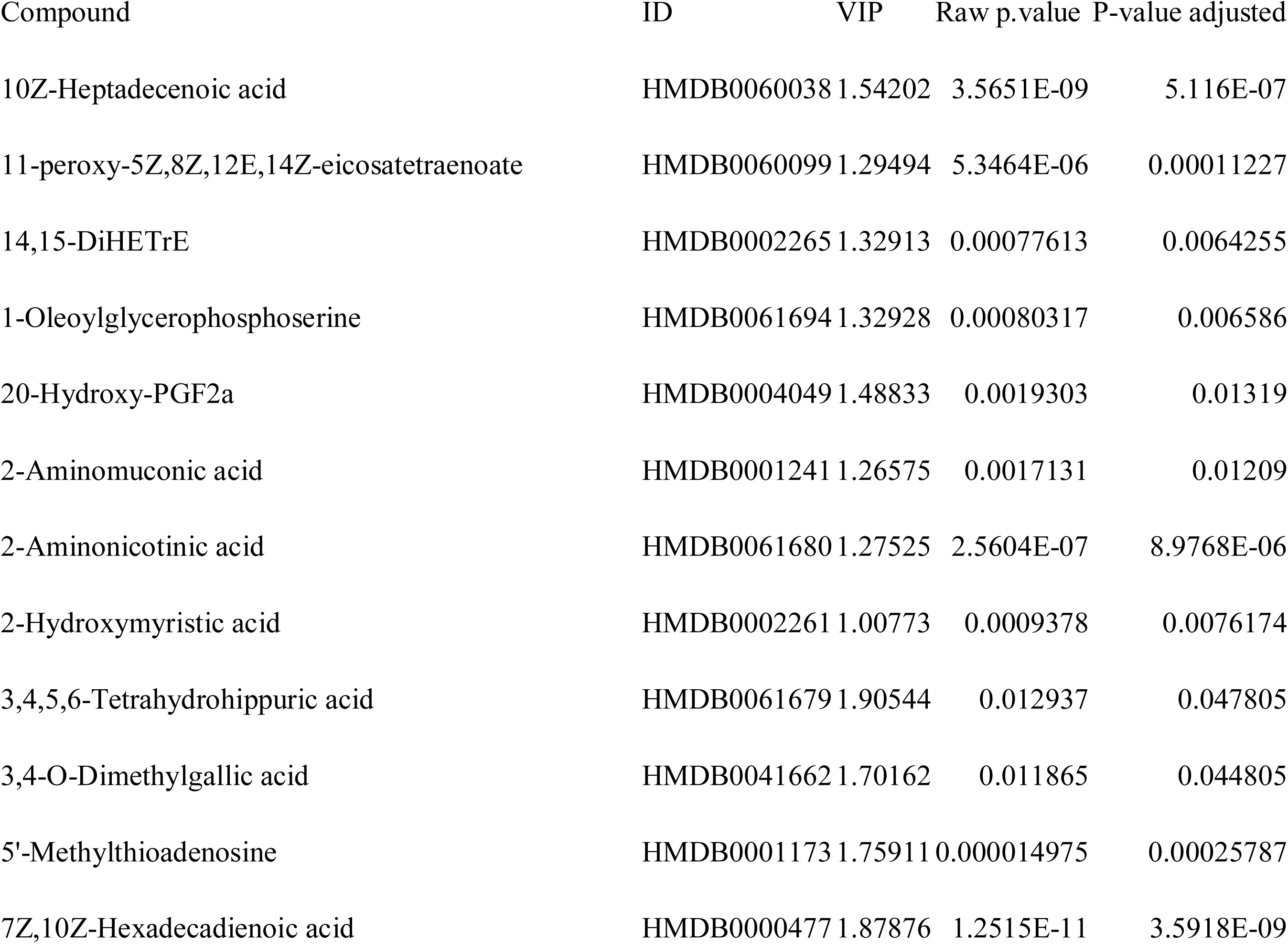
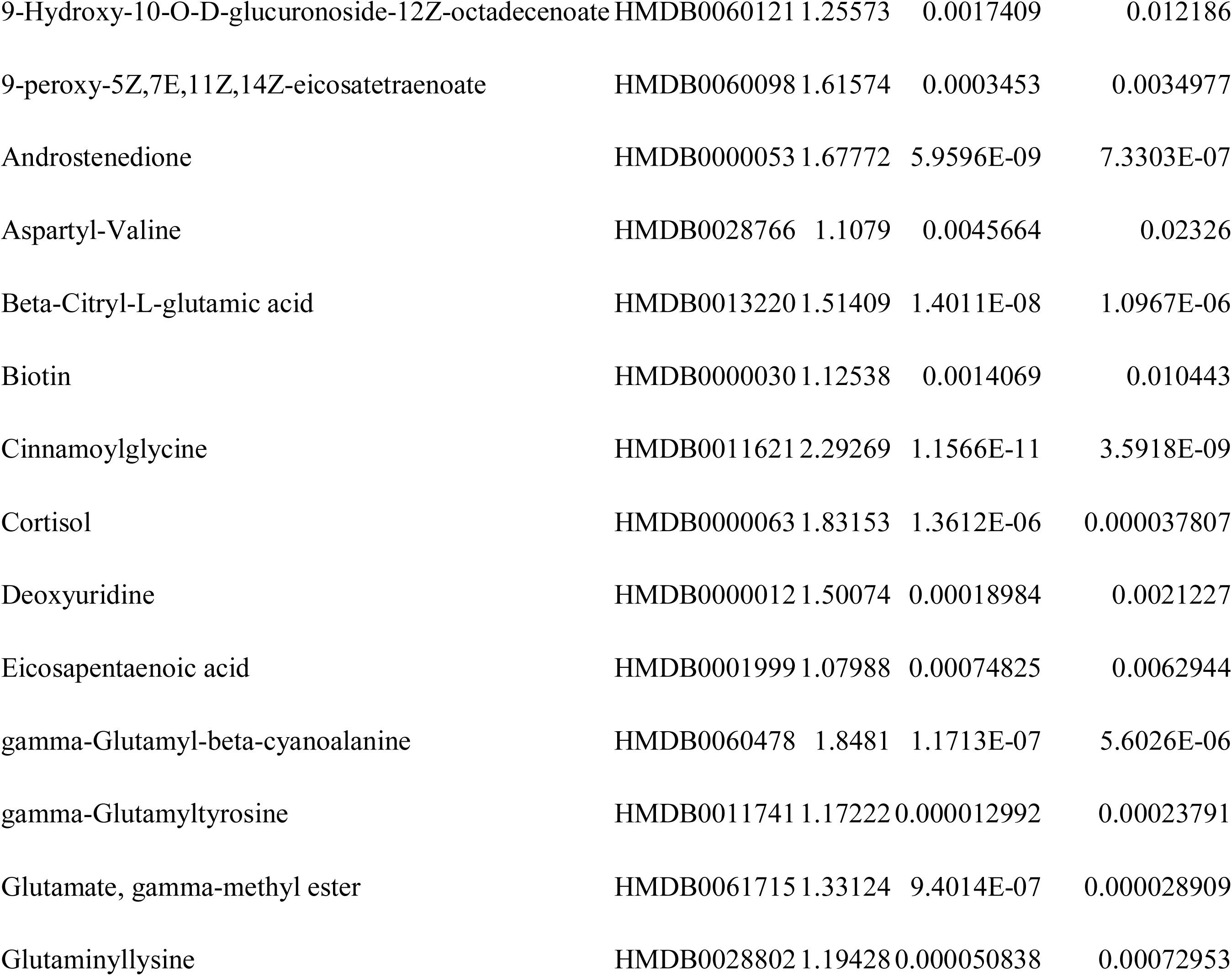
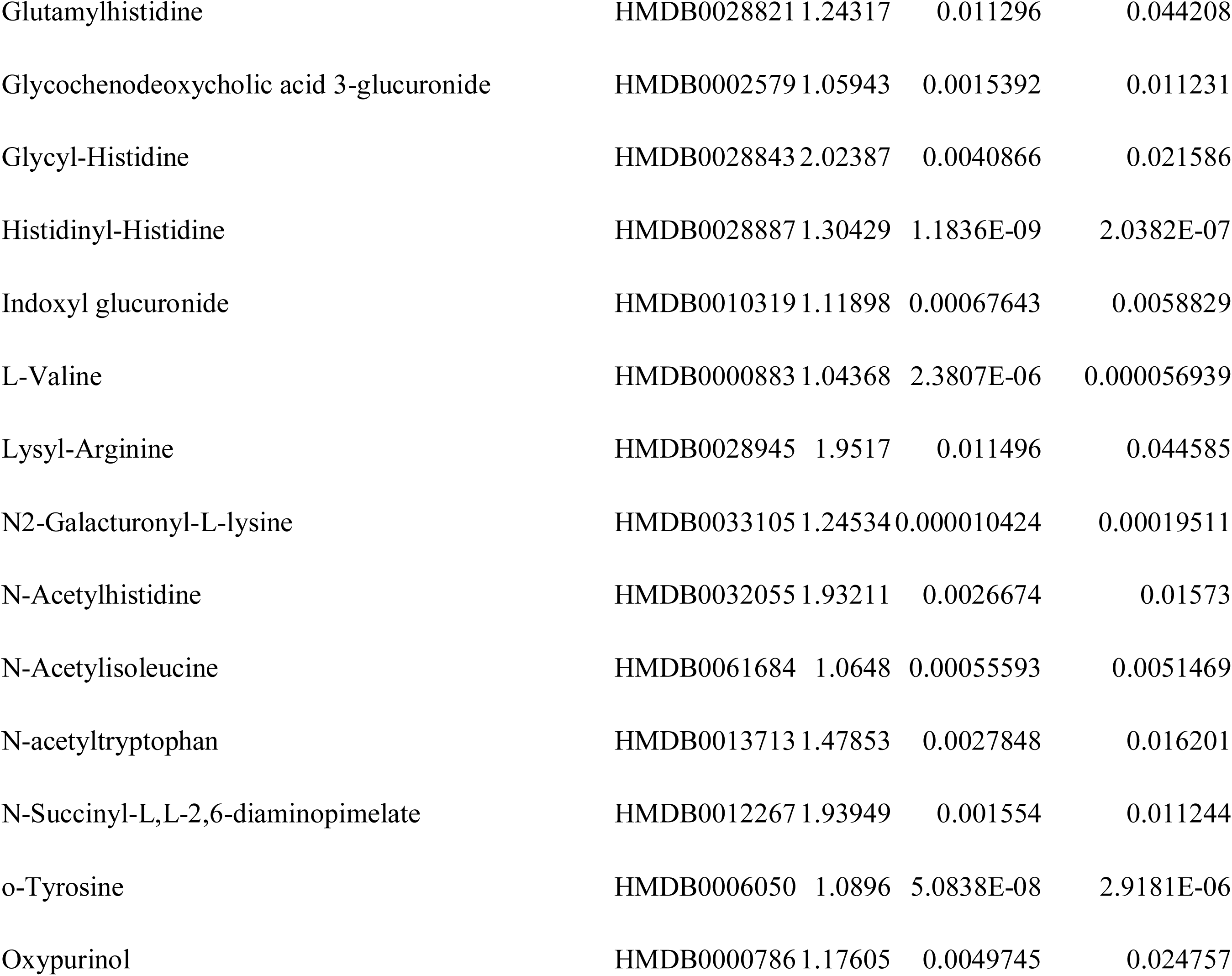
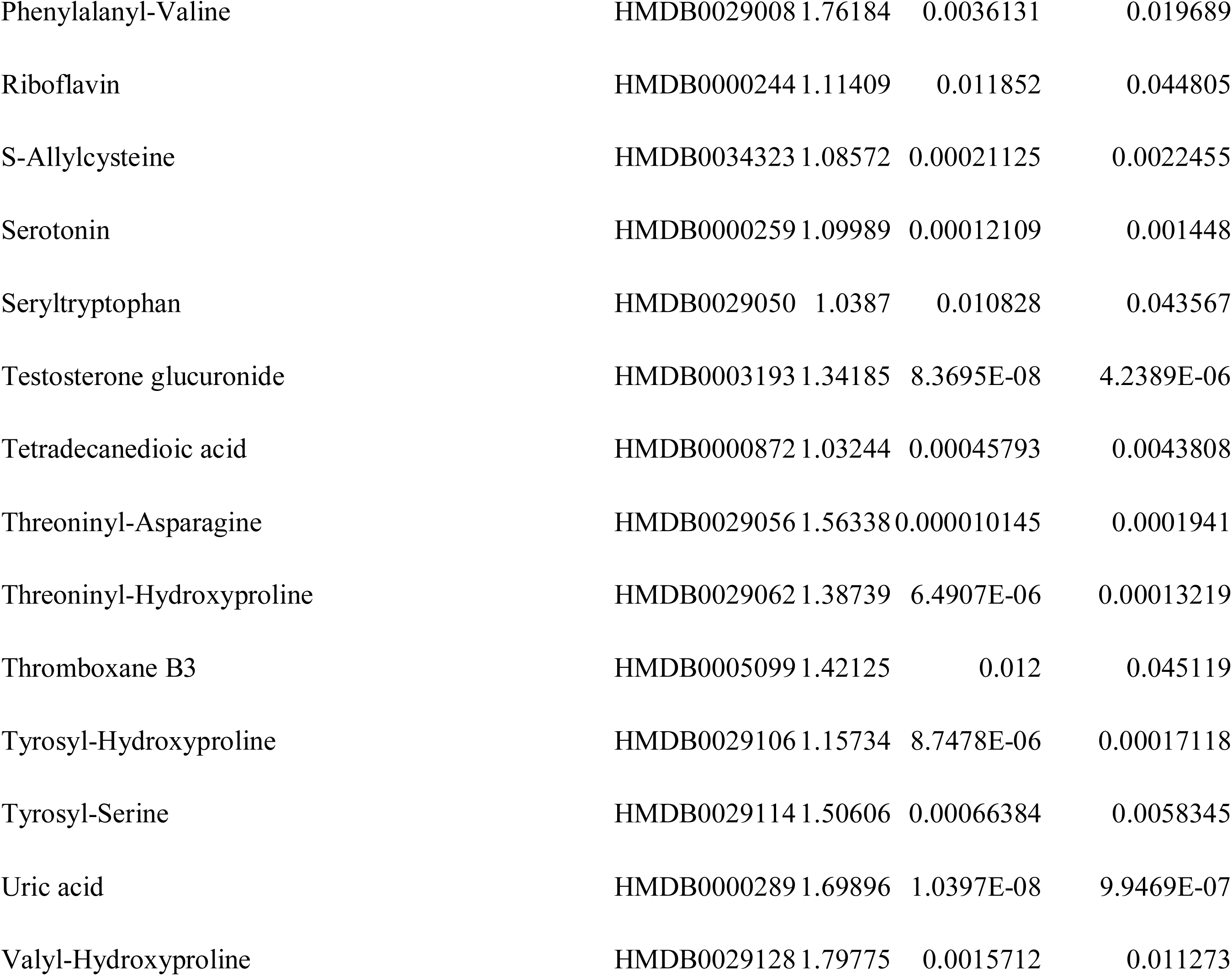
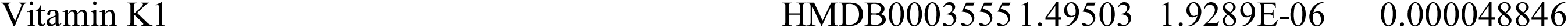
Age differential metabolites in males.

| Compound | ID | VIP | Raw p.value | P-value adjusted |
| --- | --- | --- | --- | --- |
| 10Z-Heptadecenoic acid | HMDB0060038 | 1.54202 | 3.5651E-09 | 5.116E-07 |
| 11-peroxy-5Z,8Z,12E,14Z-eicosatetraenoate | HMDB0060099 | 1.29494 | 5.3464E-06 | 0.00011227 |
| 14,15-DiHETrE | HMDB0002265 | 1.32913 | 0.00077613 | 0.0064255 |
| 1-Oleoylglycerophosphoserine | HMDB0061694 | 1.32928 | 0.00080317 | 0.006586 |
| 20-Hydroxy-PGF2a | HMDB0004049 | 1.48833 | 0.0019303 | 0.01319 |
| 2-Aminomuconic acid | HMDB0001241 | 1.26575 | 0.0017131 | 0.01209 |
| 2-Aminonicotinic acid | HMDB0061680 | 1.27525 | 2.5604E-07 | 8.9768E-06 |
| 2-Hydroxymyristic acid | HMDB0002261 | 1.00773 | 0.0009378 | 0.0076174 |
| 3,4,5,6-Tetrahydrohippuric acid | HMDB0061679 | 1.90544 | 0.012937 | 0.047805 |
| 3,4-O-Dimethylgallic acid | HMDB0041662 | 1.70162 | 0.011865 | 0.044805 |
| 5'-Methylthioadenosine | HMDB0001173 | 1.75911 | 0.000014975 | 0.00025787 |
| 7Z,10Z-Hexadecadienoic acid | HMDB0000477 | 1.87876 | 1.2515E-11 | 3.5918E-09 |
| 9-Hydroxy-10-O-D-glucuronoside-12Z-octadecenoate | HMDB0060121 | 1.25573 | 0.0017409 | 0.012186 |
| 9-peroxy-5Z,7E,11Z,14Z-eicosatetraenoate | HMDB0060098 | 1.61574 | 0.0003453 | 0.0034977 |
| Androstenedione | HMDB0000053 | 1.67772 | 5.9596E-09 | 7.3303E-07 |
| Aspartyl-Valine | HMDB0028766 | 1.1079 | 0.0045664 | 0.02326 |
| Beta-Citryl-L-glutamic acid | HMDB0013220 | 1.51409 | 1.4011E-08 | 1.0967E-06 |
| Biotin | HMDB0000030 | 1.12538 | 0.0014069 | 0.010443 |
| Cinnamoylglycine | HMDB0011621 | 2.29269 | 1.1566E-11 | 3.5918E-09 |
| Cortisol | HMDB0000063 | 1.83153 | 1.3612E-06 | 0.000037807 |
| Deoxyuridine | HMDB0000012 | 1.50074 | 0.00018984 | 0.0021227 |
| Eicosapentaenoic acid | HMDB0001999 | 1.07988 | 0.00074825 | 0.0062944 |
| gamma-Glutamyl-beta-cyanoalanine | HMDB0060478 | 1.8481 | 1.1713E-07 | 5.6026E-06 |
| gamma-Glutamyltyrosine | HMDB0011741 | 1.17222 | 0.000012992 | 0.00023791 |
| Glutamate, gamma-methyl ester | HMDB0061715 | 1.33124 | 9.4014E-07 | 0.000028909 |
| Glutaminyllysine | HMDB0028802 | 1.19428 | 0.000050838 | 0.00072953 |
| Glutamylhistidine | HMDB0028821 | 1.24317 | 0.011296 | 0.044208 |
| Glycochenodeoxycholic acid 3-glucuronide | HMDB0002579 | 1.05943 | 0.0015392 | 0.011231 |
| Glycyl-Histidine | HMDB0028843 | 2.02387 | 0.0040866 | 0.021586 |
| Histidinyl-Histidine | HMDB0028887 | 1.30429 | 1.1836E-09 | 2.0382E-07 |
| Indoxyl glucuronide | HMDB0010319 | 1.11898 | 0.00067643 | 0.0058829 |
| L-Valine | HMDB0000883 | 1.04368 | 2.3807E-06 | 0.000056939 |
| Lysyl-Arginine | HMDB0028945 | 1.9517 | 0.011496 | 0.044585 |
| N2-Galacturonyl-L-lysine | HMDB0033105 | 1.24534 | 0.000010424 | 0.00019511 |
| N-Acetylhistidine | HMDB0032055 | 1.93211 | 0.0026674 | 0.01573 |
| N-Acetylisoleucine | HMDB0061684 | 1.0648 | 0.00055593 | 0.0051469 |
| N-acetyltryptophan | HMDB0013713 | 1.47853 | 0.0027848 | 0.016201 |
| N-Succinyl-L,L-2,6-diaminopimelate | HMDB0012267 | 1.93949 | 0.001554 | 0.011244 |
| o-Tyrosine | HMDB0006050 | 1.0896 | 5.0838E-08 | 2.9181E-06 |
| Oxypurinol | HMDB0000786 | 1.17605 | 0.0049745 | 0.024757 |
| Phenylalanyl-Valine | HMDB0029008 | 1.76184 | 0.0036131 | 0.019689 |
| Riboflavin | HMDB0000244 | 1.11409 | 0.011852 | 0.044805 |
| S-Allylcysteine | HMDB0034323 | 1.08572 | 0.00021125 | 0.0022455 |
| Serotonin | HMDB0000259 | 1.09989 | 0.00012109 | 0.001448 |
| Seryltryptophan | HMDB0029050 | 1.0387 | 0.010828 | 0.043567 |
| Testosterone glucuronide | HMDB0003193 | 1.34185 | 8.3695E-08 | 4.2389E-06 |
| Tetradecanedioic acid | HMDB0000872 | 1.03244 | 0.00045793 | 0.0043808 |
| Threoninyl-Asparagine | HMDB0029056 | 1.56338 | 0.000010145 | 0.0001941 |
| Threoninyl-Hydroxyproline | HMDB0029062 | 1.38739 | 6.4907E-06 | 0.00013219 |
| Thromboxane B3 | HMDB0005099 | 1.42125 | 0.012 | 0.045119 |
| Tyrosyl-Hydroxyproline | HMDB0029106 | 1.15734 | 8.7478E-06 | 0.00017118 |
| Tyrosyl-Serine | HMDB0029114 | 1.50606 | 0.00066384 | 0.0058345 |
| Uric acid | HMDB0000289 | 1.69896 | 1.0397E-08 | 9.9469E-07 |
| Valyl-Hydroxyproline | HMDB0029128 | 1.79775 | 0.0015712 | 0.011273 |

**Table S8.**
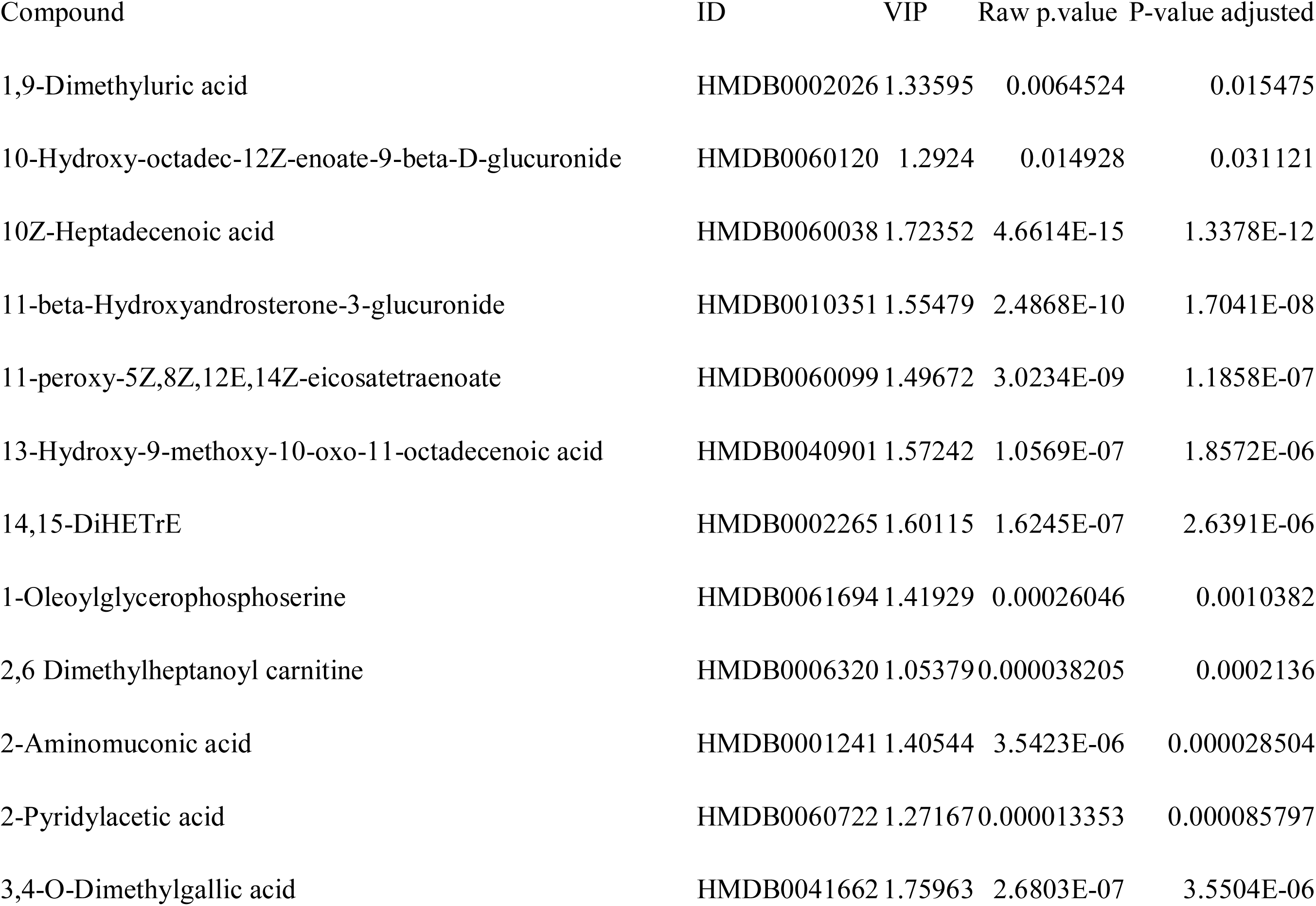
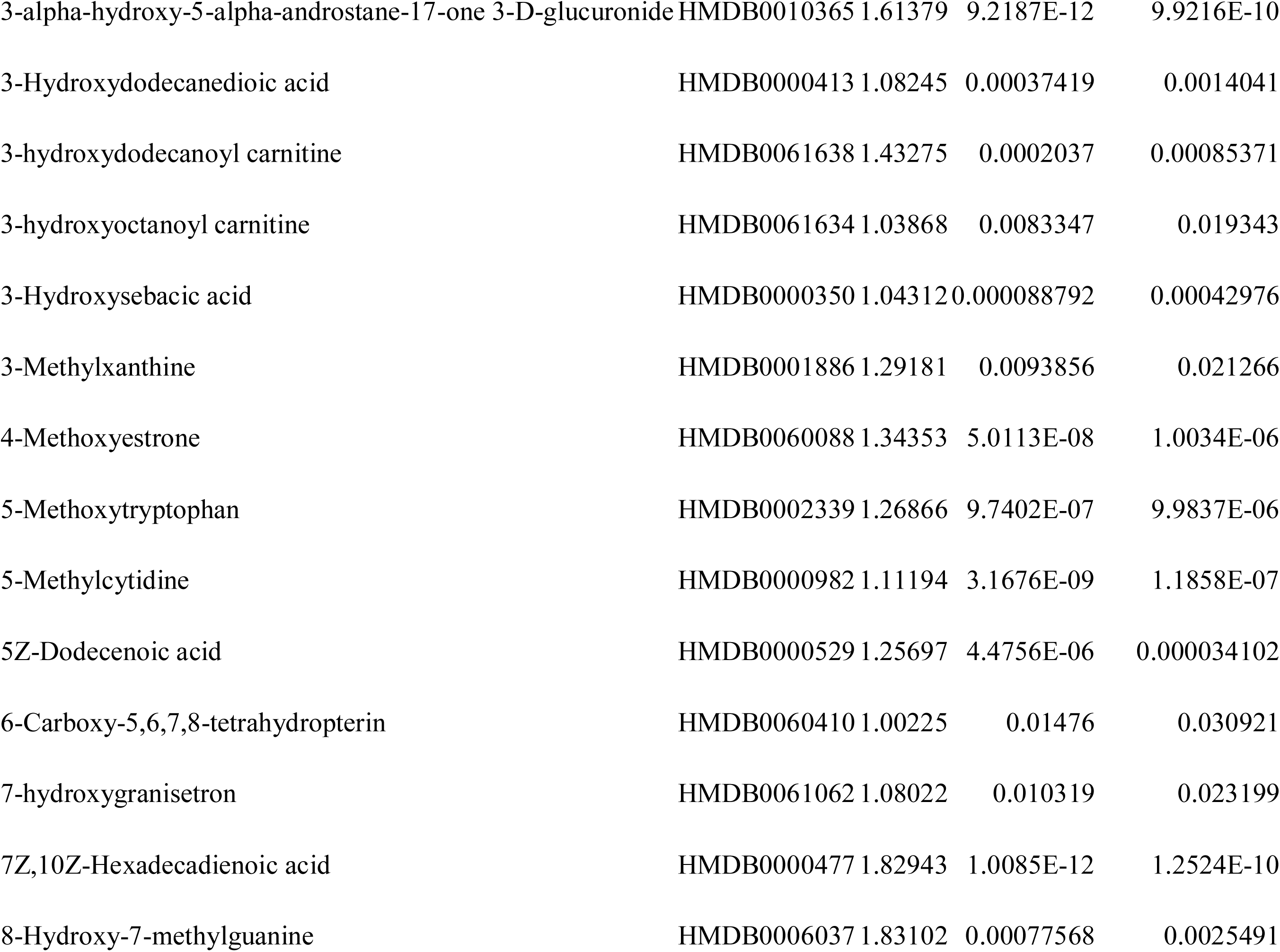
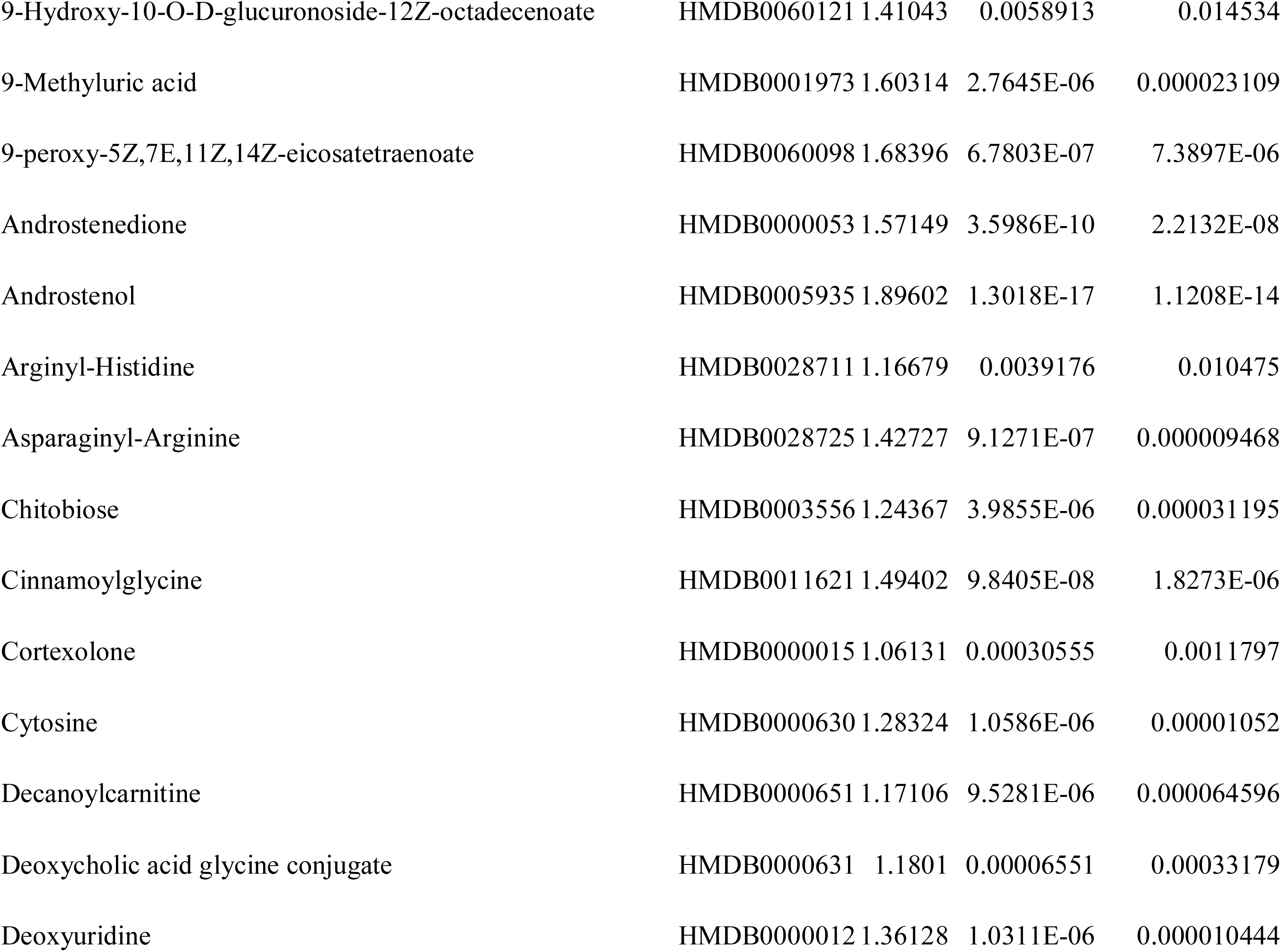
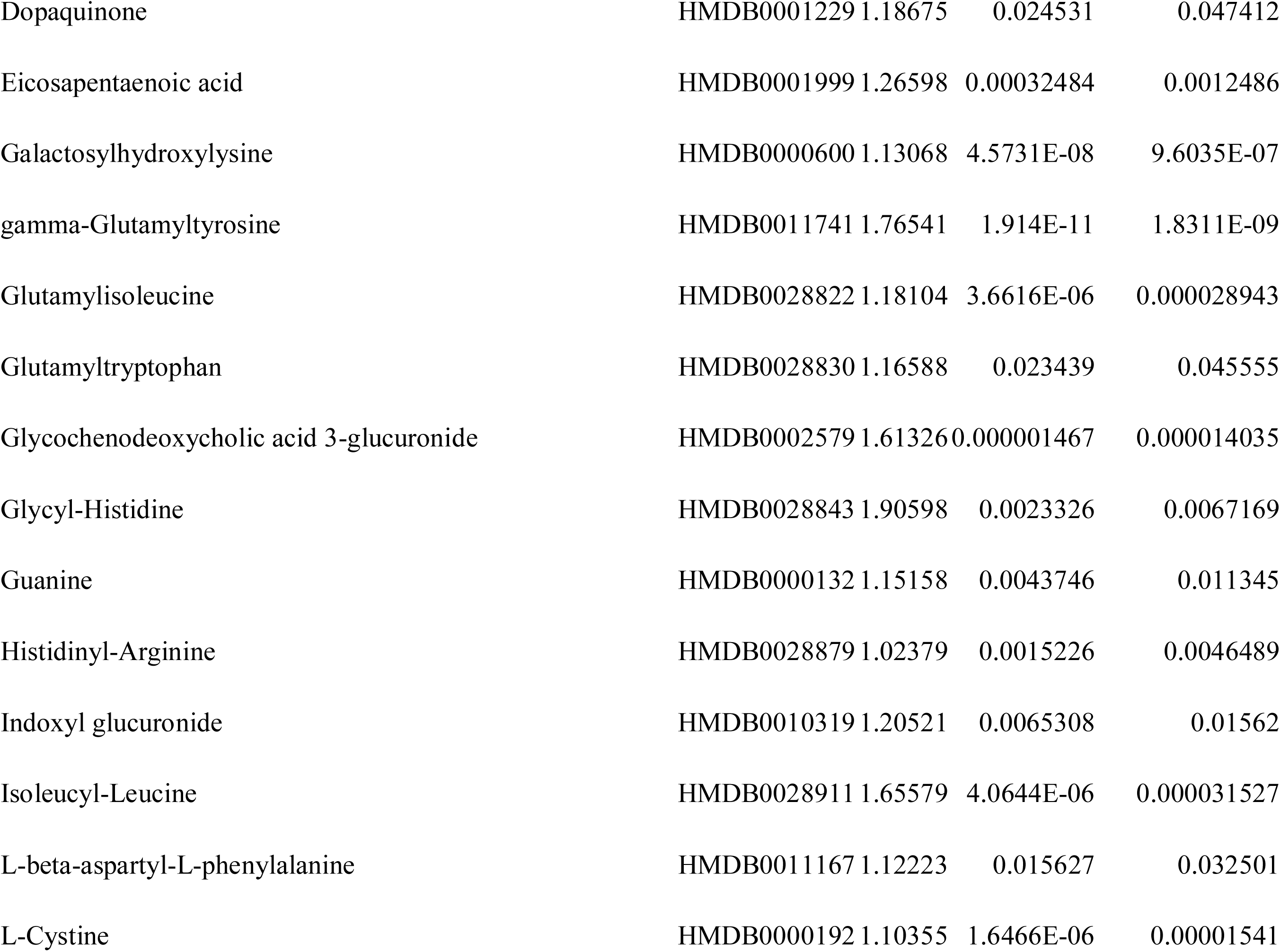
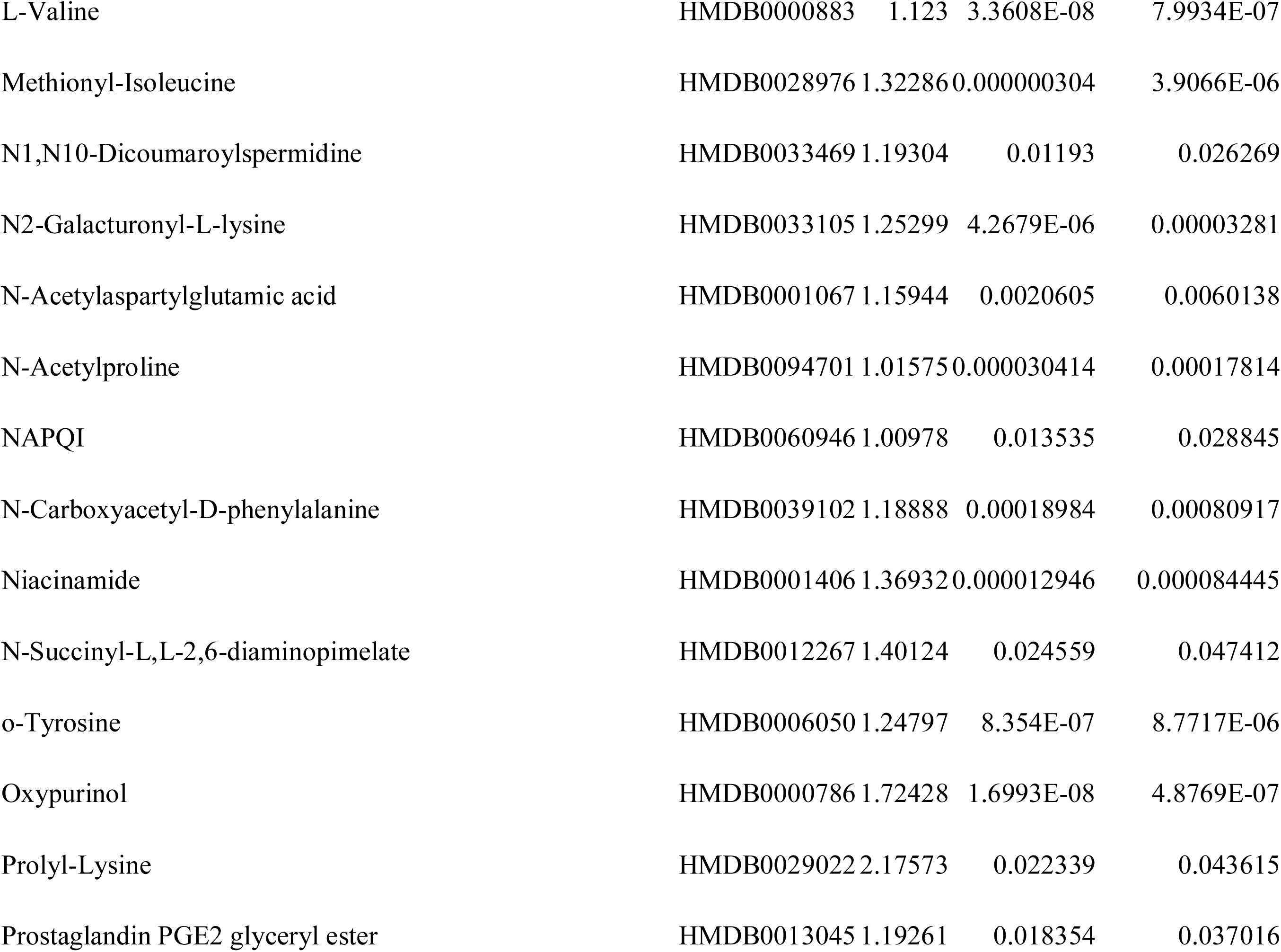
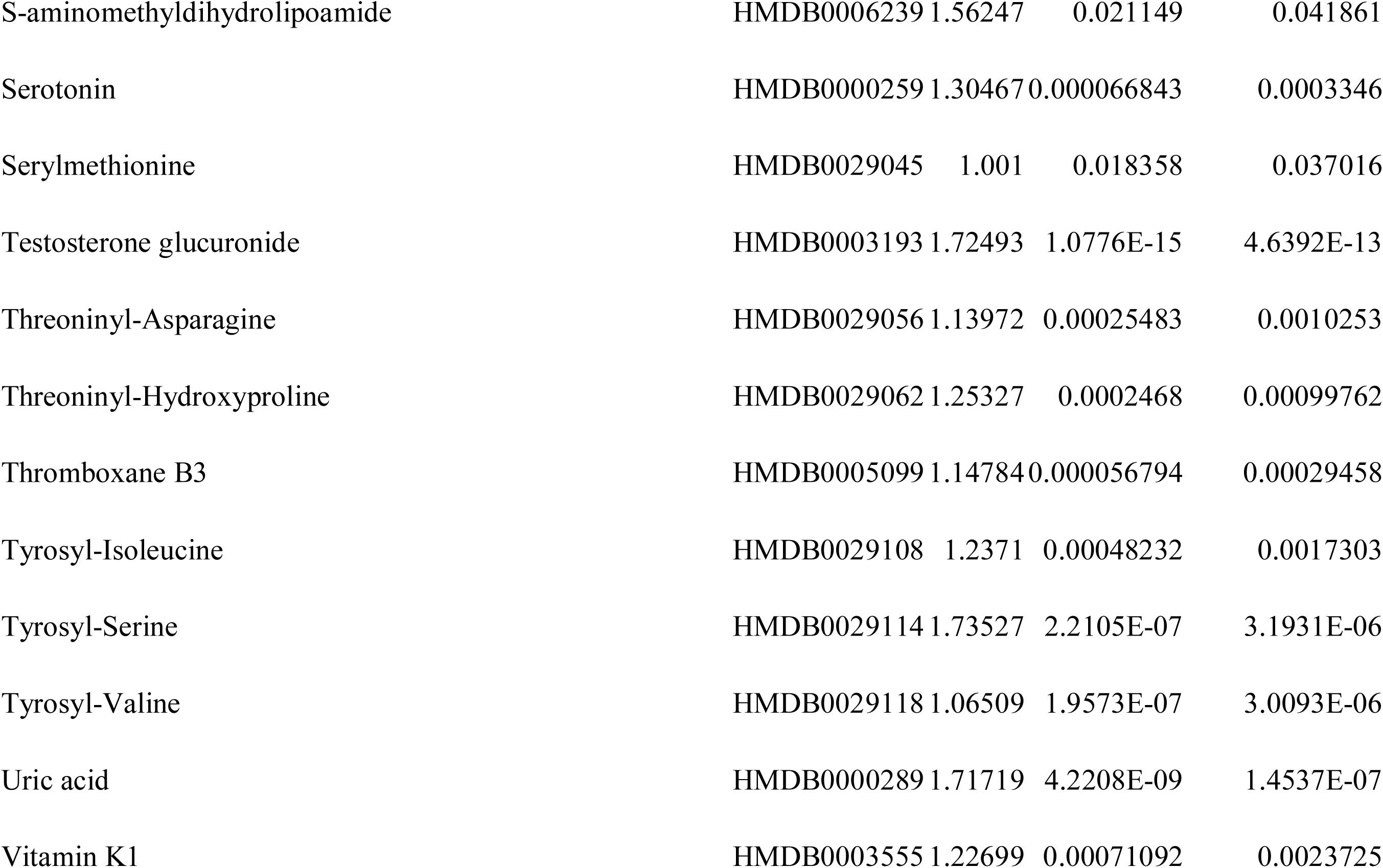
Age differential metabolites in females.

| Compound | ID | VIP | Raw p.value | P-value adjusted |
| --- | --- | --- | --- | --- |
| 1,9-Dimethyluric acid | HMDB0002026 | 1.33595 | 0.0064524 | 0.015475 |
| 10-Hydroxy-octadec-12Z-enoate-9-beta-D-glucuronide | HMDB0060120 | 1.2924 | 0.014928 | 0.031121 |
| 10Z-Heptadecenoic acid | HMDB0060038 | 1.72352 | 4.6614E-15 | 1.3378E-12 |
| 11-beta-Hydroxyandrosterone-3-glucuronide | HMDB0010351 | 1.55479 | 2.4868E-10 | 1.7041E-08 |
| 11-peroxy-5Z,8Z,12E,14Z-eicosatetraenoate | HMDB0060099 | 1.49672 | 3.0234E-09 | 1.1858E-07 |
| 13-Hydroxy-9-methoxy-10-oxo-11-octadecenoic acid | HMDB0040901 | 1.57242 | 1.0569E-07 | 1.8572E-06 |
| 14,15-DiHETrE | HMDB0002265 | 1.60115 | 1.6245E-07 | 2.6391E-06 |
| 1-Oleoylglycerophosphoserine | HMDB0061694 | 1.41929 | 0.00026046 | 0.0010382 |
| 2,6 Dimethylheptanoyl carnitine | HMDB0006320 | 1.05379 | 0.000038205 | 0.0002136 |
| 2-Aminomuconic acid | HMDB0001241 | 1.40544 | 3.5423E-06 | 0.000028504 |
| 2-Pyridylacetic acid | HMDB0060722 | 1.27167 | 0.000013353 | 0.000085797 |
| 3,4-O-Dimethylgallic acid | HMDB0041662 | 1.75963 | 2.6803E-07 | 3.5504E-06 |
| 3-alpha-hydroxy-5-alpha-androstane-17-one 3-D-glucuronide | HMDB0010365 | 1.61379 | 9.2187E-12 | 9.9216E-10 |
| 3-Hydroxydodecanedioic acid | HMDB0000413 | 1.08245 | 0.00037419 | 0.0014041 |
| 3-hydroxydodecanoyl carnitine | HMDB0061638 | 1.43275 | 0.0002037 | 0.00085371 |
| 3-hydroxyoctanoyl carnitine | HMDB0061634 | 1.03868 | 0.0083347 | 0.019343 |
| 3-Hydroxysebacic acid | HMDB0000350 | 1.04312 | 0.000088792 | 0.00042976 |
| 3-Methylxanthine | HMDB0001886 | 1.29181 | 0.0093856 | 0.021266 |
| 4-Methoxyestrone | HMDB0060088 | 1.34353 | 5.0113E-08 | 1.0034E-06 |
| 5-Methoxytryptophan | HMDB0002339 | 1.26866 | 9.7402E-07 | 9.9837E-06 |
| 5-Methylcytidine | HMDB0000982 | 1.11194 | 3.1676E-09 | 1.1858E-07 |
| 5Z-Dodecenoic acid | HMDB0000529 | 1.25697 | 4.4756E-06 | 0.000034102 |
| 6-Carboxy-5,6,7,8-tetrahydropterin | HMDB0060410 | 1.00225 | 0.01476 | 0.030921 |
| 7-hydroxygranisetron | HMDB0061062 | 1.08022 | 0.010319 | 0.023199 |
| 7Z,10Z-Hexadecadienoic acid | HMDB0000477 | 1.82943 | 1.0085E-12 | 1.2524E-10 |
| 8-Hydroxy-7-methylguanine | HMDB0006037 | 1.83102 | 0.00077568 | 0.0025491 |
| 9-Hydroxy-10-O-D-glucuronoside-12Z-octadecenoate | HMDB0060121 | 1.41043 | 0.0058913 | 0.014534 |
| 9-Methyluric acid | HMDB0001973 | 1.60314 | 2.7645E-06 | 0.000023109 |
| 9-peroxy-5Z,7E,11Z,14Z-eicosatetraenoate | HMDB0060098 | 1.68396 | 6.7803E-07 | 7.3897E-06 |
| Androstenedione | HMDB0000053 | 1.57149 | 3.5986E-10 | 2.2132E-08 |
| Androstenol | HMDB0005935 | 1.89602 | 1.3018E-17 | 1.1208E-14 |
| Arginyl-Histidine | HMDB0028711 | 1.16679 | 0.0039176 | 0.010475 |
| Asparaginy-Arginine | HMDB0028725 | 1.42727 | 9.1271E-07 | 0.000009468 |
| Chitobiose | HMDB0003556 | 1.24367 | 3.9855E-06 | 0.000031195 |
| Cinnamoylglycine | HMDB0011621 | 1.49402 | 9.8405E-08 | 1.8273E-06 |
| Cortexolone | HMDB0000015 | 1.06131 | 0.00030555 | 0.0011797 |
| Cytosine | HMDB0000630 | 1.28324 | 1.0586E-06 | 0.00001052 |
| Decanoylcarnitine | HMDB0000651 | 1.17106 | 9.5281E-06 | 0.000064596 |
| Deoxycholic acid glycine conjugate | HMDB0000631 | 1.1801 | 0.00006551 | 0.00033179 |
| Deoxyuridine | HMDB0000012 | 1.36128 | 1.0311E-06 | 0.000010444 |
| Dopaquinone | HMDB0001229 | 1.18675 | 0.024531 | 0.047412 |
| Eicosapentaenoic acid | HMDB0001999 | 1.26598 | 0.00032484 | 0.0012486 |
| Galactosylhydroxylysine | HMDB0000600 | 1.13068 | 4.5731E-08 | 9.6035E-07 |
| gamma-Glutamyltyrosine | HMDB0011741 | 1.76541 | 1.914E-11 | 1.8311E-09 |
| Glutamylisoleucine | HMDB0028822 | 1.18104 | 3.6616E-06 | 0.000028943 |
| Glutamyltryptophan | HMDB0028830 | 1.16588 | 0.023439 | 0.045555 |
| Glycochenodeoxycholic acid 3-glucuronide | HMDB0002579 | 1.61326 | 0.000001467 | 0.000014035 |
| Glycyl-Histidine | HMDB0028843 | 1.90598 | 0.0023326 | 0.0067169 |
| Guanine | HMDB0000132 | 1.15158 | 0.0043746 | 0.011345 |
| HistidinyL-Arginine | HMDB0028879 | 1.02379 | 0.0015226 | 0.0046489 |
| Indoxyl glucuronide | HMDB0010319 | 1.20521 | 0.0065308 | 0.01562 |
| Isoleucyl-Leucine | HMDB0028911 | 1.65579 | 4.0644E-06 | 0.000031527 |
| L-beta-aspartyl-L-phenylalanine | HMDB0011167 | 1.12223 | 0.015627 | 0.032501 |
| L-Cystine | HMDB0000192 | 1.10355 | 1.6466E-06 | 0.00001541 |
| L-Valine | HMDB0000883 | 1.123 | 3.3608E-08 | 7.9934E-07 |
| Methionyl-Isoleucine | HMDB0028976 | 1.32286 | 0.000000304 | 3.9066E-06 |
| N1,N10-Dicoumaroylspermidine | HMDB0033469 | 1.19304 | 0.01193 | 0.026269 |
| N2-Galacturonyl-L-lysine | HMDB0033105 | 1.25299 | 4.2679E-06 | 0.00003281 |
| N-Acetylaspartylglutamic acid | HMDB0001067 | 1.15944 | 0.0020605 | 0.0060138 |
| N-Acetylproline | HMDB0094701 | 1.01575 | 0.000030414 | 0.00017814 |
| NAPQI | HMDB0060946 | 1.00978 | 0.013535 | 0.028845 |
| N-Carboxyacetyl-D-phenylalanine | HMDB0039102 | 1.18888 | 0.00018984 | 0.00080917 |
| Niacinamide | HMDB0001406 | 1.36932 | 0.000012946 | 0.000084445 |
| N-Succinyl-L,L-2,6-diaminopimelate | HMDB0012267 | 1.40124 | 0.024559 | 0.047412 |
| o-Tyrosine | HMDB0006050 | 1.24797 | 8.354E-07 | 8.7717E-06 |
| Oxypurinol | HMDB0000786 | 1.72428 | 1.6993E-08 | 4.8769E-07 |
| Prolyl-Lysine | HMDB0029022 | 2.17573 | 0.022339 | 0.043615 |
| Prostaglandin PGE2 glyceryl ester | HMDB0013045 | 1.19261 | 0.018354 | 0.037016 |
| S-aminomethyldihydrolipoamide | HMDB0006239 | 1.56247 | 0.021149 | 0.041861 |
| Serotonin | HMDB0000259 | 1.30467 | 0.000066843 | 0.0003346 |
| Serylmethionine | HMDB0029045 | 1.001 | 0.018358 | 0.037016 |
| Testosterone glucuronide | HMDB0003193 | 1.72493 | 1.0776E-15 | 4.6392E-13 |
| Threoninyl-Asparagine | HMDB0029056 | 1.13972 | 0.00025483 | 0.0010253 |
| Threoninyl-Hydroxyproline | HMDB0029062 | 1.25327 | 0.0002468 | 0.00099762 |
| Thromboxane B3 | HMDB0005099 | 1.14784 | 0.000056794 | 0.00029458 |
| Tyrosyl-Isoleucine | HMDB0029108 | 1.2371 | 0.00048232 | 0.0017303 |
| Tyrosyl-Serine | HMDB0029114 | 1.73527 | 2.2105E-07 | 3.1931E-06 |
| Tyrosyl-Valine | HMDB0029118 | 1.06509 | 1.9573E-07 | 3.0093E-06 |
| Uric acid | HMDB0000289 | 1.71719 | 4.2208E-09 | 1.4537E-07 |
| Vitamin K1 | HMDB0003555 | 1.22699 | 0.00071092 | 0.0023725 |

